# Cold-adapted live attenuated SARS-CoV-2 vaccine completely protects human ACE2 transgenic mice from SARS-CoV-2 infection

**DOI:** 10.1101/2020.08.04.235689

**Authors:** Sang Heui Seo, Yunyueng Jang

## Abstract

Severe acute respiratory syndrome coronavirus (SARS-CoV-2) has infected more than 16,000,000 people and has caused the death of more than 650,000 individuals since December 2019. A safe and effective vaccine that can provide herd immunity against SARS-CoV-2 is urgently needed to stop the spread of this virus among humans. Many human viral vaccines are live attenuated forms of viruses that elicit humoral and cellular immunity. Here, we describe the development of a cold-adapted live attenuated vaccine (SARS-CoV-2/human/Korea/CNUHV03-CA22**°**C/2020) by gradually adapting the growth of SARS-CoV-2 from 37**°**C to 22**°**C in Vero cells. This vaccine can be potentially administered to humans through nasal spray. Its single dose was observed to strongly induce the neutralising antibody (>640), cellular immunity, and mucosal IgA antibody in intranasally immunised K18-hACE2 mice, which are very susceptible to SARS-CoV-2 and SARS-CoV infection. The one-dose vaccinated mice were completely protected from SARS-CoV-2 infection and did not show loss of body weight, death, and the presence of virus in tissues, such as the nasal turbinates, brain, lungs, and kidneys. Taken together, the cold-adapted live attenuated SARS-CoV-2 vaccine developed by us may contribute to saving of human lives from the threat of SARS-CoV-2.

---

In December 2019, human cases of mysterious severe pneumonia were reported from the city of Wuhan in eastern China (1, 2). The initial symptoms were found to be similar to those of patients infected with Severe Acute Respiratory Syndrome (SARS) virus. Sequencing of the causative agent showed that its genome was similar to that of SARS, leading to its designation as Severe Acute Respiratory Syndrome Coronavirus-2 (SARS-CoV-2) (1, 2). The World Health Organization (WHO) declared coronavirus disease 2019 (COVID-19), the disease caused by SARS-CoV-2, as a pandemic on the 11 March 2020.

SARS-CoV-2 belongs to the family of coronaviruses, which are enveloped, positive-sense single-stranded RNA viruses (3). The genome size of SARS-CoV-2 is about 30 kb. SARS-CoV-2 consists of four structural proteins, namely nucleocapsid (N), membrane (M), envelope (E), and spike (S) proteins, which form the structural backbone of the virus; sixteen non-structural proteins (nsp1–nsp16); and several accessory proteins (4). The S protein is located on the surface of SARS-CoV-2 and binds to human angiotensin converting enzyme 2 (hACE2) receptor to initiate the infection (5, 6).

Patients infected with SARS-CoV-2 commonly show fever, cough, myalgia, and fatigue, and some patients might also develop acute respiratory distress syndrome (ARDS) (**7**). Among the 99 patients infected with SARS-CoV-2 in Wuhan, China, 74 showed bilateral pneumonia, 14 showed multiple mottling and ground-grass opacity in the lungs, and one patient had pneumothorax; 17 patients had ARSD, and 11 of these patients died of multiple organ failure (8). In Washington, USA, ARDS was observed in 15 of 21 patients, and mechanical ventilation was required for these patients (9). In addition to pneumonia and ARDS, SARS-CoV-2 is responsible for clinical signs related to the affliction of the central nervous system (CNS); these include loss of taste and smell, headaches, twitching, seizures, vision impairment, nerve pain, dizziness, impaired consciousness, nausea, vomiting, hemiplegia, ataxia, stroke, and cerebral haemorrhage (10, 11).

As of date, there is no effective licenced vaccine for SARS-CoV-2. Therefore, there is an urgent need to develop a safe and effective SARS-CoV-2 vaccine to protect humans from the COVID-19 pandemic. Various types of SARS-CoV-2 vaccines are under development; these include DNA-and mRNA-based vaccines, encoding the S protein of SARS-CoV-2 (12–14), adenovirus-, measles virus-, and vesicular stomatitis virus-based vectors expressing the S gene (15–18), and purified inactivated vaccine (19). Most of the licenced human viral vaccines, such as those against measles, mumps, rubella, rotavirus, smallpox, chickenpox, yellow fever, and influenza virus (nasal inoculation), are live attenuated forms of the respective virus (20–27). Cold-adapted live influenza vaccines for seasonal influenza viruses are produced in primary chick kidney cells or embryonated eggs at 25°C and are administered intranasally to humans (26, 27). Live attenuated vaccines are similar to natural infectious agents; they elicit strong and long-lasting immune response, and thereby, have good protective effects in humans.

In this study, we developed a cold-adapted live attenuated SARS-CoV-2 vaccine strain by gradually adapting the growth of SARS-CoV-2 virus from 37°C to 22°C in Vero cells. The attenuation of the SARS-CoV-2 vaccine strain (designated as SARS-CoV-2/human/Korea/CNUHV03-CA22°C/2020) and its efficacy as a vaccine was confirmed in hACE-2 transgenic mice (K18-hACE2 mice), which were found to be lethal to SARS-CoV and SARS-CoV-2 (28–32).

To develop a live attenuated vaccine for SARS-CoV-2, amenable to intranasal delivery, we gradually adapted SARS-CoV-2 isolated from a human patient (SARS-CoV-2/human/Korea/CNUHV03/2020; referred to as CoV-2-CNUHV03 in this study) (GenBank accession number: MT678839) (33) to a temperature from 37°C to 22°C in Vero cells cultured in an atmosphere of 5% CO_2_ in a humidified incubator. When cells infected with SARS-CoV-2 showed complete cytopathic effects (CPE) at the set temperature, they were incubated at the next lower temperature. SARS-CoV-2 viruses that were successfully passaged more than five times (>p = 5) at 22°C were used for the vaccine efficacy study. SARS-CoV-2 adapted at 22°C was designated as SARS-CoV-2/human/Korea/CNUHV03-CA22°C/2020 (herein referred to as CoV-2-CNUHV03-CA22°C).

Vero cells grown in 6-well plates were infected with wild-type SARS-CoV-2 (CoV-2-CNUHV03) or cold-adapted vaccine SARS-CoV-2 (CoV-2-CNUHV03-CA22°C) in a humidified 5% CO_2_ incubator at 37°C and 41°C to determine the temperature sensitivity of the vaccine strain (**fig. S1**). At 37°C and 0.00001 multiplicity of infections (m.o.i) (**fig. S1A**), the viral titers of CoV-2-CNUHV03-CA22°C and CoV-2-CNUHV03 were 2.6 × 10^5^ plaque forming units (pfu)/ml and 7.9 × 10^5^ pfu/ml, respectively. At 37°C and 0.000001 m.o.i (**fig S1A**), those of CoV-2-CNUHV03-CA22°C and CoV-2-CNUHV03 were 0 pfu/ml and 7.9 × 10^4^ pfu/ml, respectively. At 41°C and 0.00001 m.o.i (**fig. S1B**), the viral titers of CoV-2-CNUHV03-CA22°C and CoV-2-CNUHV03 were 0 pfu/ml and 10 × 10^3^ pfu/ml, respectively. At 41°C and 0.000001 m.o.i. (**fig. S1B**), those of CoV-2-CNUHV03-CA22°C and CoV-2-CNUHV03 were 0 pfu/ml and 8 × 10^3^ pfu/ml, respectively.

To confirm the attenuation of CoV-2-CNUHV03-CA22°C in an animal, we intranasally (i.n.) infected hACE-2 transgenic mice (K18-hACE2), which are very susceptible to SARS-CoV-2, with CoV-2-CNUHV03-CA22°C (2 × 10^4^ pfu) (**Fig. 1**). The infected mice were monitored for mortality (**Fig. 1A**) and change in body weight (**Fig. 1B**) for 14 days. All these mice survived and did not show any loss of body weight, whereas all the K18-hACE2 mice infected with CoV-2-CNUHV03 showed loss of body weight (5.8%) until 8 days post-infection (p.i.) and eventually died.

**Fig. 1.**
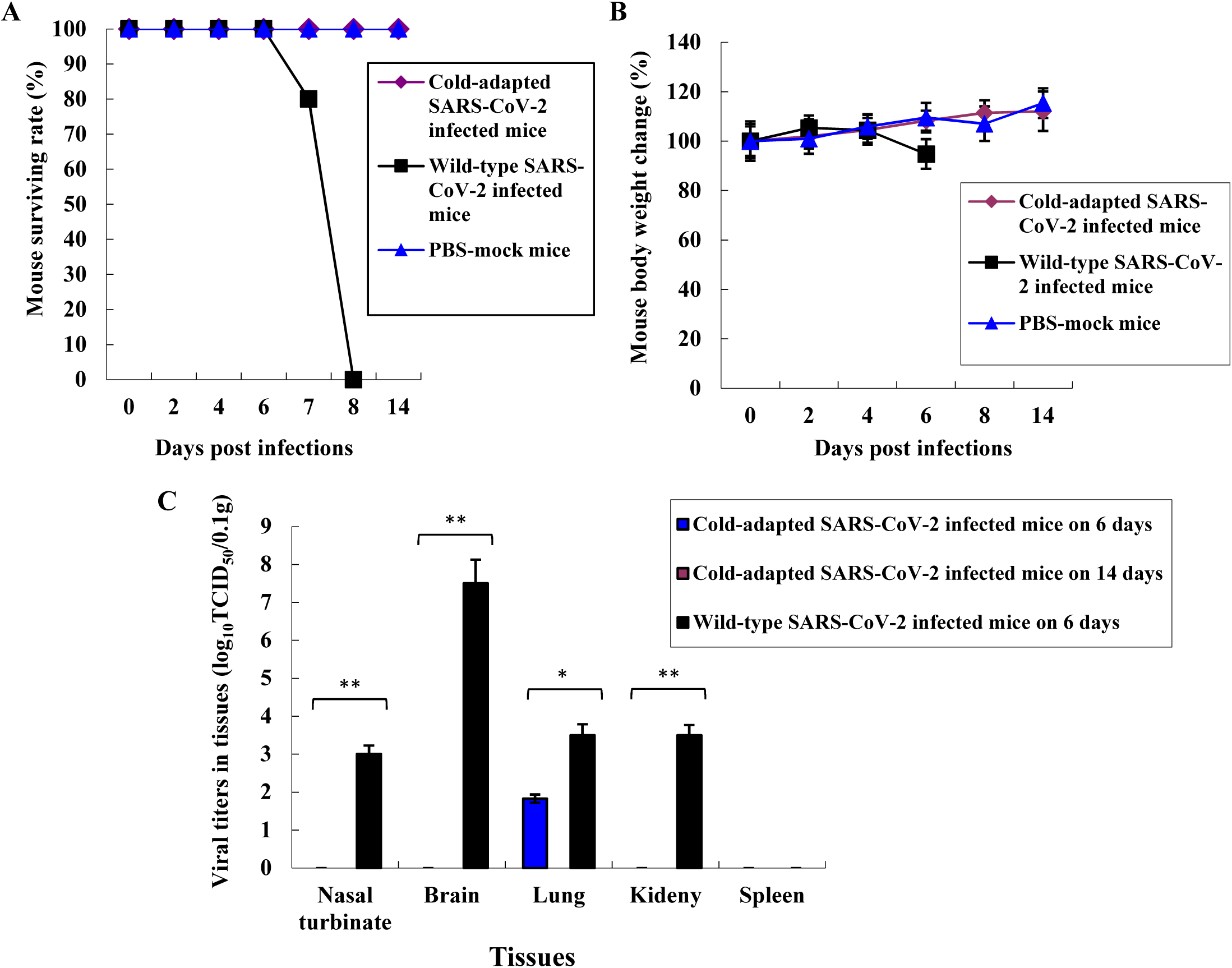
Attenuation of cold-adapted SARS-CoV-2 vaccine strain in hACE2 transgenic mice. K18-ACE2 mice were i.n. infected with cold adapted SARS-CoV-2 (CoV-2-CNUHV03-CA22°C) (2X10^4^ pfu) (n=14) or were i.n. infected with wild-type SARS-CoV-2 (CoV-2-CNUHV03) (2X10^4^ pfu) (n=6). Mice were monitored for mortality and the change of body weights for 14 days. On day 6 p.i., mice (n=3 per group) were euthanized for viral titers in tissues. One day 14 p.i., mice (n=3) infected with CoV-2-CNUHV03-CA22°C were also euthanized. PBS-mock infected K18-ACE-2 mice (n=4) were used as a control group and among them, one mice were euthanized on day 6. **A**, mice mortality rate (%); **B,** the change of mouse body weights (%) compared to those before infections, **C,** Viral titers in mouse tissues (n=3 per group) of nasal turbinate, brain, lung, kidney, and spleen by log_10_ TCID_50_/0.1g. Viral titers are the mean of 3 tissues ± standard deviations. Detection limit of virus is 1 TCID_50_/0.1g.

We measured the virus titres in different tissues (nasal turbinates, brain, lungs, kidneys, spleen) of infected mice by determining the log_10_ tissue culture infectious dose 50 (log_10_TCID_50_) values in Vero cells as well as by performing real-time quantitative polymerase chain reaction (RT-qPCR) with SARS-CoV-2 N primers and probe on day 6 p.i. The virus titres were lower in the tissues of K18-hACE2 mice infected with CoV-2-CNUHV03-CA22°C than in the tissues of K18-hACE2 mice infected with CoV-2-CNUHV03 (**Fig. 1C, & fig. S2**). When we measured the virus titres in terms of the log_10_TCID_50_ value, viruses were detected only in the lungs of K18-hACE2 mice infected with CoV-2-CNUHV03-CA22°C with a titre of 1.83 TCID_50_/0.1g; however, the virus was detected in the nasal turbinates (3.0 TCID_50_/0. g), brain (7.5 TCID_50_/0.1 g), lungs (3.5 TCID_50_/0.1 g), and kidneys (3.5 TCID_50_/0.1 g) of K18-hACE2 mice infected with CoV-2-CNUHV03 (**Fig. 1C**). Using RT-qPCR, the virus was detected in the nasal turbinates (5.9 × 10^3^ pfu/0.1 g) and lungs (11 × 10^3^ pfu/0.1 g) of K18-hACE2 mice infected with CoV-2-CNUHV03-CA22°C, whereas it was detected in the nasal turbinates (10 × 10^3^ pfu/0.1g), brain (2.5 × 10^6^ pfu/0.1g), lungs (14 × 10^3^ pfu/0.1 g), and kidneys (1.3 × 10^3^ pfu/0.1 g) of K18-hACE2 mice infected with CoV-2-CNUHV03 (**fig. S2A & S2B**). On day 14 p.i., no virus was detected in the tissues of K18-hACE2 mice infected with CoV-2-CNUHV03-CA22°C (**Fig. 1C & fig. S2A & S2B**).

We stained the lung tissue sections of K18-hACE2 mice with haematoxylin and eosin (H&E) and brain, lungs, and kidneys with the SARS-CoV-2 NP antibody (**Fig. 2, fig. S3, & fig. S4**). The lung tissue of K18-hACE2 mice infected with CoV-2-CNUHV03-CA22°C (**Fig. 2B**) showed much milder pneumonia than that of K18-hACE2 mice infected with CoV-2-CNUHV03 (**Fig. 2C**) on day 6 p.i. The antigen staining in the lung of K18-hACE2 mice infected with CoV-2-CNUHV03-CA22°C (**Fig. 2E**) was much sparse than that in mice infected with CoV-2-CNUHV03 **(Fig. 2F**) on day 6 p.i. No antigen staining was observed in the brain (**fig. S3B**) and kidneys (**fig. S3E**) of K18-hACE2 mice infected with CoV-2-CNUHV03-CA22°C on day 6 p.i., whereas profuse antigen staining was observed in the brain (**fig. S3C**) and kidneys (**fig. S3F**) of K18-hACE2 mice infected with CoV-2-CNUHV03 on day 6 p.i. No antigen staining was observed in the lungs (**fig. S4A**), brain (**fig. S4B**), and kidney (**fig. S4C**) of K18-hACE2 mice infected with CoV-2-CNUHV03-CA22°C on day 14 p.i.

**Fig. 2.**
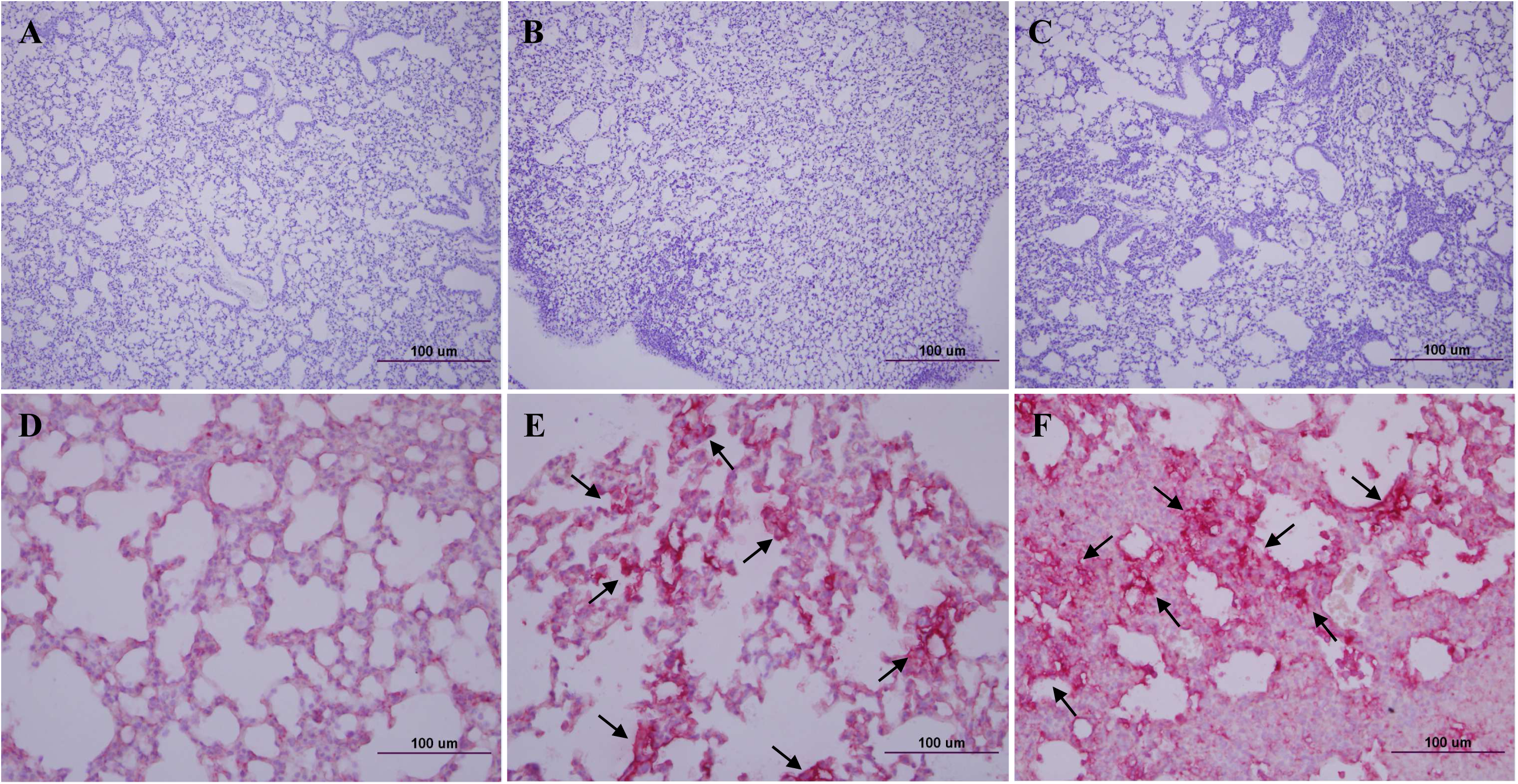
Lung histopathology and antigen staining of cold-adapted SARS-CoV-2 vaccine strain in hACE2 transgenic mice. Lung tissues (Fig. 1C) of K18-ACE2 mice on 6 days p.i. were stained with hematoxylin & eosin (A,B,C) (X100), and SARS-CoV-2 NP antibody (D,E,F) (X400). **A&D**, lung tissues of PBS-mock mice, **B&E**, lung tissues of mice i.n. infected with cold adapted SARS-CoV-2 (CoV-2-CNUHV03-CA22°C) (2X10^4^ pfu), **C&F**, lung tissues of mice infected with wild-type SARS-CoV-2 (CoV-2-CNUHV03) (2X10^4^ pfu). Arrow: positive antigen staining.

K18-hACE2 mice were immunised by i.n. administration of 2 × 10^4^ or 2 × 10^3^ pfu of CoV-2-CNUHV03-CA22°C, and sera were collected 19 days post vaccination (p.v.). The titre of the neutralising antibody (NA) was measured using CoV-2-CNUHV03 and CoV-2-KCDC03 in Vero cells. Strong NA titres, in the range of 640–4960, were induced in K18-hACE2 mice immunised with 2 × 10^4^ (**Fig. 3A**) or 2 × 10^3^ (**Fig. 3B**) pfu of CoV-2-CNUHV03-CA22°C. No NA was detected in the sera of K18-hACE2 mice, collected before vaccination (**fig. S5**). We measured the levels of IgA antibody, which is responsible for mucosal immunity, in the different tissues (nasal turbinates, lungs, and kidneys) using purified inactivated SARS-CoV-2 antigen (CoV-2-CNUHV03) and goat horseradish peroxidase (HRP)-labelled anti-mouse IgA antibody (**fig. S6A**) and T cells expressing IFN-γ (**fig. S6B**). The detection of IgA indicates the induction of cellular immunity in splenocytes in K18-hACE2 mice immunised with of CoV-2-CNUHV03-CA22°C (2 × 10^4^ pfu). IgA was detected in all the tissues that were assessed, with the highest amount detected in nasal turbinates (OD: 0.298) (**fig. S6A**). The number of IFN-γ expressing T cells in the immunised and PBS (mock)-immunised K18-hACE2 mice was 1682/250,000 and 249/250,000 splenocytes, respectively (**fig. S6B**).

**Fig. 3.**
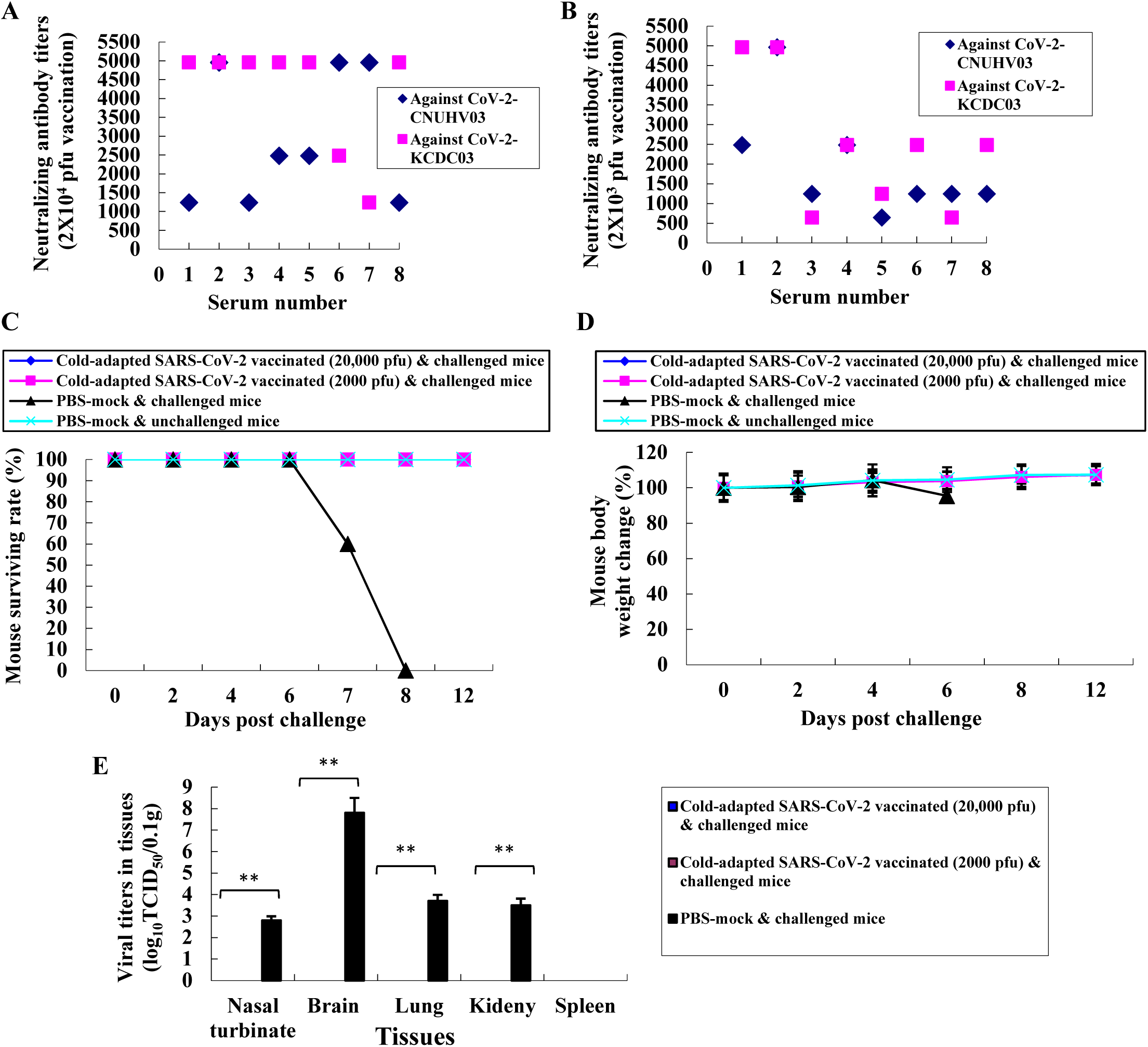
Neutralizing antibodies and challenge of immunized hACE2 transgenic mice with cold-adapted SARS-CoV-2 vaccine strain. Sera collected from K18-ACE2 mice (n=8 per group) immunized with 2X10^4^ pfu or 2X10^3^ pfu of cold adapted SARS-CoV-2 (CoV-2-CNUHV03-CA22°C) 19 days ago, and their neutralizing antibody titers were determined against wild-type SARS-CoV-2 viruses, CoV-2-CNUHV03 and CoV-2-KCDC03 in Vero cells. **A**, Sera from immunized mice with 2X10^4^ pfu of CoV-2-CNUHV03-CA22°C; **B**, Sera from immunized mice with 2X10^3^ pfu of CoV-2-CNUHV03-CA22°C. Sera (n=16) collected before immunization were used as a control. Detection limit of neutralizing antibody is 10. *p<0.05, **p<0.001. K18-ACE2 mice (n=8 per group) immunized with 2X10^4^ pfu or 2X10^3^ pfu of cold adapted SARS-CoV-2 (CoV-2-CNUHV03-CA22°C) 21 days ago were i.n. challenged with 2X10^4^ pfu of wild-type SARS-CoV-2 (CoV-2-KCDC03). PBS-mock vaccinated mice (n=6) were also i.n. challenged with 2X10^4^ pfu of CoV-2-KCDC03 and PBS-mock and unchallenged mice (n=3) were used as a control. The challenged mice were monitored for mortality and the change of body weights for 12 days. On 6 days post challenge, the challenged mice (n=3) were euthanized for viral titers in tissues. One mouse of PBS-mock and unchallenged mice was euthanized on day 6. **C**, mice mortality rate (%); **D,** the change of mouse body weights (%) compared to those before challenge, **E,** Viral titers in mouse tissues (n=3 per group) of nasal turbinate, brain, lung, kidney, and spleen by log_10_TCID_50_/0.1g. Viral titers are the mean of 3 tissues ± standard deviations. Detection limit of virus is 1 TCID_50_/0.1g.*p<0.05, **p<0.001

K18-hACE2 mice were i.n. immunised with 2 × 10^4^ or 2 × 10^3^ pfu of CoV-2-CNUHV03-CA22°C and i.n. challenged with 2 × 10^4^ pfu of CoV-2-KCDC03 on 21 days p.v. The challenged K18-hACE2 mice were monitored for mortality (**Fig. 3C**), and change in body weight (**Fig. 3D**) for 12 days p.i.; the virus titres in different tissues (nasal turbinates, lungs, brain, kidneys, and spleen) were measured 6 days post-challenge (p.c.) by determining the log10TCID50 values (**Fig. 3E**) and by performing RT-qPCR (**fig. S7A & S7B**). All the immunised and challenged K18-hACE2 mice survived **(Fig. 3C**) and did not show loss of body weight (**Fig. 3D**), whereas all the PBS (mock)-immunised and challenged K18-hACE2 mice died (**Fig. 3C**) within 8 days p.c., and showed loss of body weight (4.6%) (**Fig. 3D**). No virus titre was detected in the nasal turbinates, brain, lungs, kidneys, and spleen of immunised K18-hACE2 mice determined in terms of the log10TCID50 value (**Fig. 3E**) as well as by RT-qPCR (**fig. S7A & S7B**). Considerable virus titres were detected in the nasal turbinates (2.8TCID_50_/0.1g, 12.0 × 10^3^ pfu/0.1 g), brain (7.8TCID_50_/0.1 g, 2.7 × 10^6^ pfu/0.1 g), lungs (3.7TCID_50_/0.1 g, 15.0 × 10^3^ pfu/0.1 g), and kidneys (3.5TCID_50_/0.1 g, 1.4 × 10^3^ pfu/0.1g) in PBS (mock)-immunised and challenged K18-hACE2 mice (**Fig. 3E & fig. S7A & S7B**). As in the case of PBS (mock)-immunised and unchallenged K18-hACE2 mice (**Fig. 4A**), H&E staining of lung tissue samples from challenged K18-hACE2 mice immunised with 2 × 10^3^ pfu (**Fig. 4B**) and 2 × 10^4^ pfu (**Fig. 4C**) of CoV-2-CNUHV03-CA22°C showed mild pneumonia and no pneumonia, respectively. The lung tissue of PBS (mock)-immunised and challenged K18-hACE2 mice exhibited severe interstitial pneumonia with infiltration of inflammatory cells (**Fig. 4D**). As in the case of PBS (mock)-immunised and unchallenged K18-hACE2 mice (**Fig. 4E**), no positive staining was detected with SARS-CoV-2 NP antibody in the lung tissues of challenged K18-hACE2 mice immunised with 2 × 10^3^ pfu (**Fig. 4F**) or 2 × 10^4^ (**Fig. 4G**) pfu of CoV-2-CNUHV03-CA22°C. Numerous positively stained regions were observed in the lung tissue of PBS (mock)-immunised and challenged K18-hACE2 mice (**Fig. 4H**). No antigen staining was observed in the brain (**fig. S8B**) and kidneys (**fig. S8F**) of challenged K18-hACE2 mice (immunised with 2 × 10^3^ pfu) as well as in the brain (**fig. S8C**) and kidneys (**fig. S8G**) of challenged K18-hACE2 mice (immunised with 2 × 10^4^ pfu) as was observed for the brain (**fig. S8A**) and kidneys (Extended Data Fig. 8e) of PBS (mock)-immunised and unchallenged K18-hACE2 mice. Strong positive antigen staining was found in the brain (**fig. S8D**) and kidneys (**fig. S8H**) of PBS (mock)-immunised and challenged K18-hACE2 mice.

**Fig. 4.**
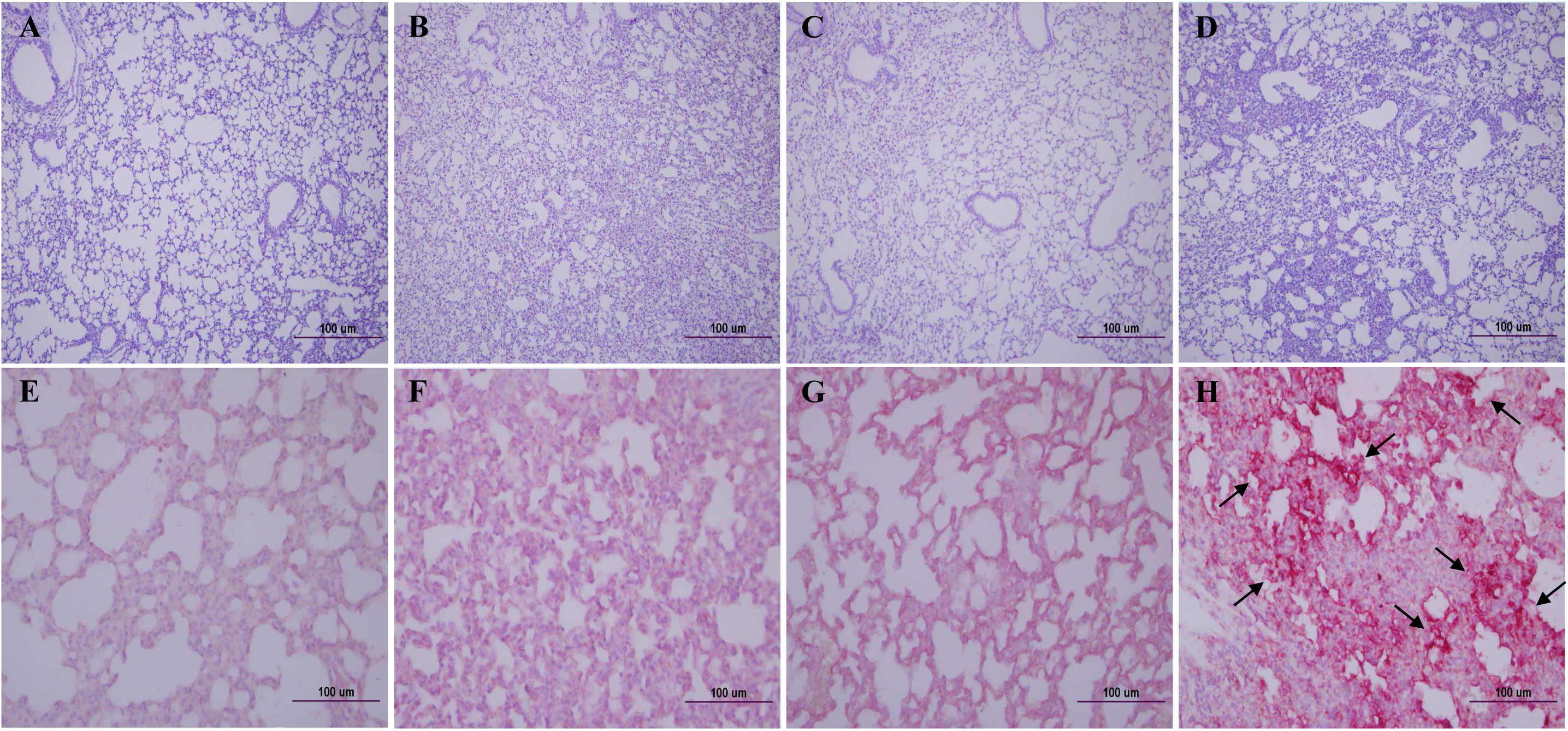
Lung histopathology and antigen-staining of the immunized and challenged hACE2 transgenic mice. Lung tissues (Fig. 3E) of the immunized and challenged K18-ACE2 were stained with hematoxylin & eosin (A,B,C,D) (X100) and SARS-CoV-2 NP antibody (E,F,G,H) (X400). **A&E**, Lung tissue of PBS-mock mouse; **B&F** Lung tissue of the immunized mouse (2X10^3^ pfu) and challenged with CoV-2-KCDC03 (2X10^4^ pfu); **C&G**, Lung tissue of the immunized mouse (2X10^4^ pfu) and challenged with CoV-2-KCDC03 (2X10^4^ pfu); **D&H**, Lung tissue of PBS-mock and challenged mouse with CoV-2-KCDC03 (2X10^4^ pfu). Arrow: positive antigen staining.

We fully sequenced the genes of CoV-2-CNUHV03-CA22°C and compared the sequences with those of wild-type SARS-CoV-2 (CoV-2-CNUHV03) (**Table 1, Table S1, Table S2, fig. S9, fig. S10**). Among the 29,874 nucleotides, 59 including 37 nonsynonymous substitutions and 22 synonymous substitutions occurred in CoV-2-CNUHV03-CA22°C compared to those of CoV-2-CNUHV03 (**Table S1 & fig. S9**). Among the 9,755 amino acid residues, 31 were mutated in CoV-2-CNUHV03-CA22°C compared to that in CoV-2-CNUHV03 (**Table S2, fig. S10**). To identify the possible unique mutations in CoV-2-CNUHV03-CA22°C, we compared the changed amino acids in CoV-2-CNUHV03-CA22°C with those in the genes of SARS-CoV-2 present in the GenBank (https://www.ncbi.nlm.nih.gov/nuccore) and GISAID (https://www.gisaid.org/), and found 12 such amino acids out of 9755 amino acids in CoV-2-CNUHV03-CA22°C (**Table 1**). In nsp2 (non-structural protein 2), with no known function (34), amino acid residues from 82 to 84 [glycine (G), histidine (H), and valine (V)] were deleted, and there was one mutation (M (methionine) 85V). In nsp6, which functions as a potential transmembrane scaffold protein (35), two mutations [N (asparagine) 3609K (lysine), D (aspartic acid) 3671T (threonine)] were present, and in nsp7, which functions as a processivity clamp for RNA polymerase (36), one mutation [D3926A (alanine)] was present. In helicase (nsp13) acting as RNA 5′-triphosphatase (37), one mutation [L (leucine)5604F (phenylalanine)] was present, and in S protein, which binds to the receptors (38), three mutations [T95I (isoleucine), N185K, S (Serine)968A] were present. When we sequenced the genome of cold-adapted live attenuated SARS-CoV-2 vaccine strain (CoV-2-CNUHV03-CA22°C) in lung tissues of infected K18-ACE2 mice on day 6 p.i., no reverted changes of genes were found (data not shown). Viral growth titer of cold-adapted live attenuated SARS-CoV-2 vaccine strain in Vero cells in tissue culture flask (75cm^2^) at 22°C is about 2.7 × 10^6^ pfu/ml, which is very comparable to that (3.0 × 10^6^ pfu/ml) of wild type SARS-CoV-2 (CoV-2-CNUHV03) at 37°C (data not shown).

**Table 1.**
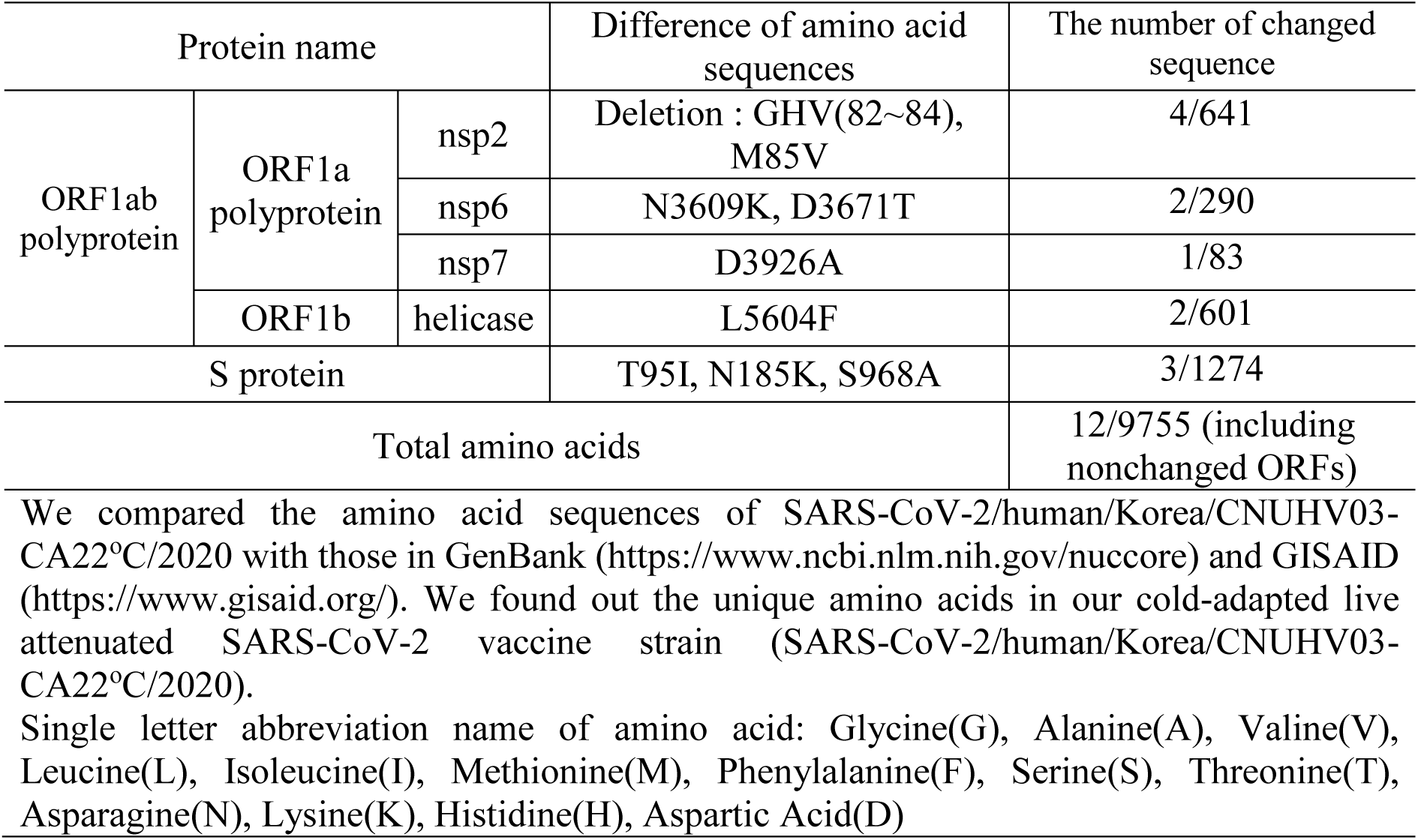
Unique sequences of amino acids in cold-adapted live attenuated SARS-CoV-2 vaccine strain (SARS-CoV-2/human/Korea/CNUHV03-CA22°C/2020)

SARS-CoV-2 has been continuously spreading in humans since December 2019. Several efforts have been made by scientists around the world to develop safe and effective vaccines to prevent SARS-CoV-2 infection in humans (39). Since March 2020, we have been in the process of developing a cold-adapted live attenuated SARS-CoV-2 vaccine that can elicit mucosal and cellular immunity and can be administered through the nasal route in humans. Our strategy had been to gradually adapt SARS-CoV-2 virus to temperatures from 37°C to 22°C in Vero cells. We confirmed the efficacy of the developed cold-adapted live attenuated SARS-CoV-2 vaccine (CoV-2-CNUHV03-CA22°C) in K18-hACE2 mice, which can be readily infected with SARS-CoV-2 and succumb to death upon infection.

We developed a cold-adapted live attenuated vaccine for SARS-CoV-2 by adapting it to grow in Vero cells at 22°C. This vaccine strain was found to be sensitive to temperatures of 37°C and 41 °C. The currently used live attenuated vaccine (nasal spray vaccine) for seasonal influenza viruses (influenza A and B viruses) was developed by adapting the virus to multiply in primary chicken kidney cells and embryonated eggs at temperature up to 25°C^10^. Nasal spray influenza vaccines for the 2019–2020 season contained four influenza viruses (influenza A (H1N1) virus, influenza A (H3N2) virus, and two influenza B viruses). Nasal spray influenza vaccines are approved for use in non-pregnant individuals who are 2 years to 49 years of age (40).

A strong neutralising antibody (640 to 4960) was induced in all the 16 immunised K18-hACE2 mice, which were i.n. inoculated with one dose (2 × 10^4^ or 2 × 10^3^ pfu) of CoV-2-CNUHV03-CA22°C. SARS-CoV-2-specific IgA antibody was also induced in the nasal turbinates, lungs, and kidneys. In addition, T lymphocytes expressing IFN-γ, which are specific for SARS-CoV-2, were strongly induced in the spleen of vaccinated mice at 19 days p.i. In a study on a measles virus (MeV)-based vaccine expressing the spike protein of SARS-CoV-2 (MeVvac2-SARS-S (H) (16), it was observed that after the second immunisation, the neutralising antibody titre for measles virus in all the immunised mice was from 257 to 800, but the titre against SARS-CoV-2 was from 15 to 80 in three out of six immunised mice.

We show that all the K18-hACE2 mice i.n. immunised with one dose (2 × 10^4^ or 2 × 10^3^ pfu) of CoV-2-CNUHV03-CA22°C were completely protected from the infection of wild-type SARS-CoV-2 (CoV-2-KCDC03) and did not show any loss of body weight, detection of virus in different tissues (nasal turbinates, brain, lungs, kidneys) in terms of the log_10_TCID_50_ values or as determined by RT-qPCR. The study on the adenovirus-based ChAdOX1 nCoV-19 vaccine, expressing the spike protein of SARS-CoV-2, showed significantly reduced virus titres in bronchoalveolar lavage fluid and respiratory tract tissues of vaccinated rhesus macaques that were challenged with SARS-CoV-2 (41).

In conclusion, cold-adapted live attenuated SARS-CoV-2 vaccine (CoV-2-CNUHV03-CA22°C) is safe in K18-hACE2 mice, and one-dose vaccination can completely protect K18-hACE2 mice from the challenge of SARS-CoV-2.

## Author information

SHS designed the study, developed cold-adapted live attenuated SARS-CoV-2 vaccine strain, performed the cold sensitivity of vaccine strain, all animal study and antigen staining of tissues, analyzed the data, and wrote the manuscript. YJ sequenced the genome of cold-adapted live attenuated SARS-CoV-2 vaccine and performed the staining of tissues with H&E. All authors approved the submitted version of manuscript.

## Data and materials availability

All data supporting the results of this study are within the manuscript, and its Extended Data tables and figures, and are available from the corresponding author when they are requested. The sequence of cold-adapted live attenuated SARS-CoV-2 vaccine strain (CoV-2-CNUHV03-CA22°C) is available in GenBank: MT810119.

## SUPPLEMENTARY MATERIALS

### Materials and Methods

#### Viruses and cells

The SARS-CoV-2 strain (SARS-CoV-2/human/Korea/CNUHV03/2020) (referred to as CoV-2-CNUHV03 in this paper) (GenBank accession number: MT678839), isolated in our laboratory from a human clinical sample collected at the Chungnam National University Hospital (Daejeon, South Korea), and BetaCoV/South Korea/KCDC03/2020 (referred to as CoV-2-KCDC03 in this paper), which was provided by the Korean Centers for Disease Control and Prevention (KCDC), were propagated in Vero-E6 cells obtained from American Type Culture Collection (Manassas, Virginia, USA). Minimal essential medium (MEM), supplemented with 10% foetal bovine serum (FBS) and 1× antibiotic–antimycotic solution (Sigma, St. Louis, USA), was used for the culture of cells. All experimental procedures involving potential contact with SARS-CoV-2 were conducted in a biosafety level 3 laboratory certified by the Korean government.

#### Animals

Female (5–6-week-old) human angiotensin converting enzyme 2 (ACE-2) transgenic mice (B6.Cg-Tg(K18-ACE2)2Prlmn/J) (referred to as K18-hACE2 in this paper) were kindly provided by The Jackson Laboratory (Bar Harbor, Maine, USA). The mice were fed a standard chow diet and water.

#### Development of the cold-adapted live attenuated SARS-CoV-2 vaccine strain

The SARS-CoV-2 strain (SARS-CoV-2/human/Korea/CNUHV03/2020) was gradually adapted from 37°C to 22 °C in Vero cells in MEM with 200 mM L-glutamine (Hyclone, South Logan, Utah, USA), supplemented with 1.5% bovine serum albumin (BSA; Rocky Mountain Biologicals, Missoula, MT, USA) and 1× antibiotic–antimycotic solution (Sigma). Vero cells were cultured in MEM with 10% FBS in a humidified 5% CO_2_ incubator (37°C) and washed twice with warm PBS (pH 7.4). The washed Vero cells were inoculated with SARS-CoV-2 virus (SARS-CoV-2/human/Korea/CNUHV03/2020) and incubated in a humidified 5% CO_2_ incubator (from 37°C to 22°C). When the infected Vero cells showed cytopathic effects (CPE), the next lower temperature was used to adapt the virus. When SARS-CoV-2 virus was successfully passaged at 22°C more than five times (>passage = 5), it was used for the vaccine study, and for sequencing of the whole genome. The cold-adapted live attenuated vaccine strain was designated as SARS-CoV-2/human/Korea/CNUHV03-CA22°C/2020 (referred to as CoV-2-CNUHV03-CA22°C in this paper). At passage 5, at 22 °C, the SARS-CoV-2 cold-adapted vaccine virus was cloned by limited-dilution infection in Vero cells in 96-well plates three times. Virus titres for the cold passaged SARS-CoV-2 were determined by RT-qPCR using SARS-CoV-2 N primers and Taqman probe and by plaque assay at 22 °C in terms of pfu.

#### Confirmation of temperature sensitivity of the cold-adapted live attenuated SARS-CoV-2 vaccine strain

Vero cells grown to confluence in 6-well plates were infected with 0.00001 or 0.000001 multiplicity of infections (m.o.i) of CoV-2-CNUHV03-CA22°C and wild-type SARS-CoV-2 (CoV-2-CNUHV03). The infected cells were incubated in a humidified 5% CO_2_ incubator at 37°C or 41°C, and virus titres in the supernatants were quantified 3 days later by RT-qPCR using the SARS-CoV-2 N primers and probe.

#### Measurement of plaque forming units by plaque assay

Stocks of SARS-CoV-2 (CoV-2-CNUHV03 or CoV-2-CNUHV03-CA22°C) were serially 10-fold diluted in MEM with 1.5% BSA. Confluent Vero cells growing in 24-well plates were infected with the diluted virus suspensions for 4h in a humidified 5% CO_2_ incubator (37 °C for CoV-2-CNUHV03 and 22 °C for CoV-2-CNUHV03-CA22°C). After removing the inoculum, Vero cells were overlaid with 1% electrophoretic agar (LPS Solution, Korea) in MEM and incubated for 4 or 7 days in a humidified 5% CO_2_ incubator (37 °C for CoV-2-CNUHV03 and 22 °C for CoV-2-CNUHV03-CA22°C). The cells were then stained with 0.1% crystal violet (Sigma-Aldrich, St. Louis, MO, USA) prepared in 37% formaldehyde solution, or with SARS-CoV-2 NP antibody and fluorescent-labelled secondary antibody. After removal of agar, the cells were fixed and permeabilised with 80% cold acetone (Samchun Pure Chemical Co., Gyeonggi-do, Korea). The cells were treated with SARS-CoV-2 nucleocapsid rabbit polyclonal antibody (Thermo Fisher Scientific, Waltham, MA, USA) and subsequently with fluorescent-labelled goat anti-rabbit antibody (Thermo Fisher Scientific). The number of plaques was counted under a fluorescence microscope (Olympus, Tokyo, Japan).

#### Measurement of virus titres using real-time quantitative PCR

RNA from virus samples was isolated using the RNeasy Mini Kit (QIAGEN, Hilden, Germany). Briefly, 100 μL of supernatant containing the virus was disrupted in 350 μL Buffer RLT, and then 550 μL of 70% ethanol was added. The sample (700 μL) was transferred to the RNeasy Mini spin column and subjected to centrifugation for 15s at 13,500 rpm. After discarding the flow-through, 700 μL of RW1 buffer was added to the spin column and it was centrifuged for 15s at 13,500 rpm. The flow-through was discarded and 500 μL of RPE buffer was added to the spin column before it was centrifuged for 15s at 13,500 rpm. The spin column was then placed in a new 1.5 mL collection tube and viral RNA was eluted using 40 μL of RNAse-free water.

To detect the virus, we used TaqMan real-time fluorescent PCR with TOPreal^TM^ One-step RT qPCR Kit (Enzynomics, Daejeon, Korea) and SARS-CoV-2 N primers and probe. In a total volume of 20 μL, the following components were mixed: 5 μL of TOPreal^TM^ One-step RT qPCR Kit (TaqMan probe), 1 μL of 10 pmol primers containing N_Sarbeco_F (5′-CACATTGGCACCCGCAAT-3′), N_Sarbeco-R (5′-GAGGAACGAGAAGAGGCTTG-3′), and N_Sarbeco_P (5′FAM-ACTTCCTCAAGGAACAACATTGCCA-3′BHQ1) (1), 10 μL of viral RNA, and 2 µ L of nuclease-free water. Real-time amplification was performed on a Rotor-Gene 6000 system (QIAGEN, Hilden, Germany) using the following temperature profile: initial incubation at 50°C for 30 min and at 95°C for 10 min, followed by 45 cycles of 95°C for 5s and 60°C for 30s. Standard curves were generated using data for stock viruses with known pfu titres determined by plaque assay.

#### Confirmation of attenuation of the cold-adapted live attenuated SARS-CoV-2 vaccine strain in hACE-2 transgenic mice

K18-hACE2 mice were i.n. immunised with 50 μL (2 × 10^4^ pfu) of the cold-adapted vaccine strain (n = 14) or wild-type virus (n = 6) after they were lightly anaesthetized with isoflurane USP (Gujarat, India). PBS (mock)-infected mice (n = 4) were used as controls. The infected mice were monitored for body weight change, and mortality. Six days p.i., three mice per virus (vaccine strain or wild-type virus)-infected group and one PBS (mock)-infected mouse were euthanized for determining virus titres in different tissues (nasal turbinates, brain, lungs, and kidneys) and for histopathology. Tissues (0.1 g per sample) were homogenised using a BeadBlaster homogeniser (Benchmark Scientific, Edison, New Jersey, USA) in 1 mL of PBS (pH 7.4) to measure virus titres by RT-qPCR and by determining the log_10_TCID_50_/0.1 g values. The remaining portions of tissues were used for histopathology and antibody staining.

#### Staining of tissues by haematoxylin and eosin

Mouse tissues were fixed in 10% neutral buffered formalin (10%) and then embedded in paraffin. The lung tissue was cut into 5 μm sections, which were stained with haematoxylin (H) solution for 4 min. The stained tissue sections were washed with tap water for 10 min and then stained with eosin (E) solution. The stained sections were visualised under an Olympus DP70 microscope and photographed (Olympus Corporation, Tokyo, Japan).

#### Staining of tissues with the SARS-CoV-2 NP antibody

Tissue sections were stained with SARS-CoV-2 nucleocapsid rabbit polyclonal antibody (Thermo Fisher Scientific). The sections were treated with antigen retrieval solution in a microwave oven and blocked with normal rabbit serum in PBS (pH 7.4). They were then incubated with the rabbit antibody against SARS-CoV-2 NP (1:100 dilution), and subsequently treated with biotin-labelled goat anti-rabbit immunoglobulin (Vector Laboratories, Burlingame, CA, USA), and Vectastain ABC-AP (Vector Laboratories) and Vector Red alkaline phosphatase substrate (Vector Laboratories). The labelled lung sections were counterstained with haematoxylin QS (Vector Laboratories) and observed under an Olympus DP70 microscope (Olympus Corporation).

#### Measurement of virus titres in terms of log_10_TCID_50_/mL

Vero cells grown in tissue culture flasks were detached by treatment with trypsin-EDTA and were seeded in 96-well tissue culture plates with MEM containing 10% FBS and 1× antibiotic-antimycotic solution. When confluent, the cells were washed with warm PBS (pH 7.4) and infected with virus samples, which were 10-fold diluted in MEM with 1.5% BSA. The cells in four wells were infected with the diluted virus samples for 4 days in a humidified incubator at 37°C (wild-type SARS-CoV-2 strain) or 22°C (cold-adapted SARS-CoV-2 vaccine strain). The cells were then fixed and permeabilised with 80% cold acetone (Samchun Pure Chemical Co.). They were subsequently incubated with SARS-CoV-2 nucleocapsid rabbit polyclonal antibody (Thermo Fisher Scientific) and fluorescent-labelled goat anti-rabbit antibody (Thermo Fisher Scientific). The titre was calculated using the method described by Muench and Reed (2).

#### Assessment of the efficacy of vaccine in hACE-2 transgenic mice

On 21 days p.v., the hACE-2 mice (n = 8 per group) immunised with 2 × 10^4^ or 2 × 10^3^ pfu of CoV-2-CNUHV03-CA22°C were i.n. challenged with 50µL (2 × 10^4^ pfu) of CoV-2-KCDC03. PBS (mock)-immunised hACE-2 mice (n = 6) were also i.n. challenged with 50 µ L (2 × 10^4^ pfu) of CoV-2-KCDC03. PBS (mock)-vaccinated and uninfected hACE-2 mice (n = 3) were used as controls. The infected mice were monitored body weight change, and mortality. Six days post-challenge, three mice per virus (vaccine strain or wild-type virus)-infected group and one PBS (mock)-infected mouse were euthanized for determining virus titres in different tissues (nasal turbinate, brain, lung, and kidney) and for histopathology. Virus titres were quantified by RT-qPCR and by determining the log_10_TCID_50_/0.1 g values.

#### Measurement of neutralising antibody titres

Sera (n = 8 per group) collected from hACE-2 mice that were i.n. immunised with 50 μL (2 × 10^4^ or 2 × 10^3^ pfu) of CoV-2-CNUHV03-CA22°C on 19 day after vaccination were 10-fold diluted in MEM with 1.5% BSA and then serially two-fold diluted before they were incubated with 100TCID_50_/mL (100 μL:100 μL) of wild-type SARS-CoV-2 virus, CoV-2-CNUHV03 or CoV-2-KCDC03, for 1 h in a humidified 5% CO_2_ incubator (37°C). Vero cells grown in a 96-well cell culture plate were washed with warm PBS (pH 7.4) and were inoculated with a mixture of serum and virus. Cells were incubated for 4 days and checked for CPE. The titre of the neutralising antibody was determined as the reciprocal of the highest dilution of serum at which the infectivity was neutralised in 100% of the cell in wells. The assay was performed in four replicates. Sixteen sera samples collected from hACE-2 mice before vaccination were used as controls.

#### Enzyme-linked immunosorbent spot assay (ELISpot) assay for mouse IFN**−**γ

The immunised hACE-2 transgenic mice (n = 3) i.n. inoculated with 50 μL (2 × 10^4^ pfu) of CoV-2-CNUHV03-CA22°C were euthanized to collect the spleen on 19 day p.v. The spleen samples were homogenised in PBS (pH 7.4) and the cells were collected. The nasal turbinates, lungs, and kidneys were homogenised in 10% PBS (pH 7.4) and the homogenates were used for detection of IgA specific for SARS-CoV-2 by enzyme-linked immunosorbent spot (ELISpot) assay. The collected cells were overlaid on HISTOPAQUE-1077 (Sigma) and centrifuged for 30 min at 1500 rpm at 4°C. The lymphocyte layer was collected for the IFN-γ ELISpot assay performed using the Mouse IFN-γ ELISpot^Plus^ kit (MABTECH, Nacka Strand, Sweden). The plate was removed from the sealed package and washed four times with sterile PBS (200 μL/well). The plate was conditioned with RPMI 1640 medium (200 μL/well) containing 10% FBS for 30 min at room temperature. The purified lymphocytes (250,000/well) mixed with 0.01 m.o.i of CoV-2-CNUHV03-CA22°C were added to the wells and the plate was incubated in a humidified 5% CO_2_ incubator at 37°C for 24 h. Thereafter, the solution was decanted from the plate and the cells were washed five times with PBS (pH 7.4) (200 μL PBS/well for each wash). The detection antibody (R4-6A2-biotin) diluted to 1 g/mL in PBS (pH 7.4) containing 0.5% FBS (200 μL/well) was added and the plate was incubated for 2 h at room temperature. The plate was washed five times with PBS (pH 7.4) (200 μL PBS/well for each wash) and streptavidin-ALP (1:1000) in PBS-0.5% FBS (100 μL/well) was added to the wells. The cells were incubated for 1 h at room temperature. The plate was washed five times with PBS as described above and the substrate solution (BCIP/NBT-plus) was added (100 μL/well). The plate was developed until distinct spots emerged. The colour development was stopped by extensive washing under tap water. Spots were inspected and counted under a microscope (Olympus). Three PBS (mock)-immunised hACE-2 mice were used as controls.

#### Detection of IgA antibody specific for SARS-CoV-2 in tissues of immunised mice by enzyme-linked immunosorbent assay (ELISA)

The purified and inactivated SARS-CoV-2 antigen (SARS-CoV-2/human/Korea/CNUHV03/2020) was diluted to a final concentration of 100 μg/mL in a coating buffer (carbonate–bicarbonate buffer, pH 9.6). The diluted antigen (100 μL) was coated onto the wells of a Nunc-Immuno^™^ MicroWell^™^ 96 well solid plate (Sigma-Aldrich) by incubated overnight at 4 °C. After removing the coating buffer, the plate was washed twice by filling the wells with 400 μL of washing buffer (0.05% Tween 20 PBS (pH 7.4) containing 4% horse serum). To block the remaining protein-binding sites, 400 μL of blocking buffer (PBS containing 4% skimmed milk) was added to the plate and the plate was kept overnight at 4 °C. The buffer was removed, and the supernatant (100 μL) of the homogenised tissue (nasal turbinates, lungs, and kidneys) from vaccinated and non-vaccinated mice 19 days after vaccination, diluted 10-fold in blocking buffer, was added to the plate and incubated for 1 h at room temperature. The plate was washed four times with the washing buffer. Goat anti-mouse IgA cross-adsorbed secondary antibody HRP (Invitrogen, MA, USA) (100 μL), diluted 1:5000 in blocking buffer, was added to each well and incubated for 1 h at room temperature. After washing the plate four times with the washing buffer, 100 μL TMB ELISA substrate (MABTECH) was dispensed in each well and the plate was incubated for 30 min at 4 °C. ABTS^®^ Peroxidase Stop Solution (KPL, MD, USA) (100 μL) was subsequently added to each well. The absorbance of the solution in each well was measured at 450 nm using an iMARK^™^ Microplate Absorbance Reader (Bio-Rad, CA, USA).

#### Sequencing of the full genome of cold-adapted live attenuated SARS-CoV-2 vaccine strain

The genome of CoV-2-CNUHV03-CA22°C was fully sequenced using overlapping primers (**Table S3**). Viral RNA was extracted using the RNeasy Mini Kit (QIAGEN, Venlo, Netherlands). Tissue culture supernatant (200 μL) containing the virus was disrupted in 350 μL Buffer RLT, and then 500 μL of 70% ethanol was added to the mixture. The disrupted samples (700 μL) were transferred to the RNeasy Mini spin column, and the column was centrifuged for 15 s at 13,500 rpm. After discarding the flow-through, 700 μL of RW1 buffer was added to the spin column and it was centrifuged for 15 s at 13,500 rpm. The flow-through was again discarded and 500 μL of RPE buffer was added to the spin column, which was then centrifuged for 2 min at 13,500 rpm. The spin column was placed in a new 1.5 mL collection tube and viral RNA was eluted with 50 μL of RNAse-free water. The extracted RNA was reverse transcribed to cDNAs using GoScript^™^ Reverse Transcription System (Promega, Madison, USA) and 12 reverse primers (covid2500R, covid5000R, covid7500R, covid10000R, covid12500R, covid15000R, covid17500R, covid20000R, covid22500R, covid25000R, covid27500R, covid29843R) (**Table S4**). Twelve viral genes were amplified by PCR with GoTaq Hot Start Green Master Mix (Promega) and a segment-specific primer set. Amplicons were separated by gel electrophoresis and purified using the QIAquick Gel Extraction Kit (QIAGEN). The purified genes were cloned into the pGEM-T Easy vector (Promega) and the vector construct was used for transformation of chemically competent *Escherichia coli* DH5α cells (Enzynomics, Daejeon, Korea). The plasmids were extracted using the HiGene Plasmid Mini Prep Kit (BIOFACT, Daejeon, Korea) and the sequences were determined by Macrogen (Seoul, Korea). Three clones per segment were sequenced. The sequenced genes were arranged using DNASTAR Lasergene (Madison, Wisconsin, USA). The sequence of CoV-2-CNUHV03-CA22°C was deposited in GenBank under the accession number MT810119.

#### Ethical approval

The protocol (202003-CNU-023) for the study of SARS-CoV-2 vaccine efficacy in mice was approved by the Internal Animal Use Committee at Chungnam National Univer sity (CNU). All the studies were approved and were conducted in accordance with the relevant legal guidelines and regulations prescribed by CNU, Republic of Korea.

#### Statistical analysis

Differences between mice infected with cold-adapted live attenuated SARS-CoV-2 vaccine strain and wild-type SARS-CoV-2 virus or between vaccinated and PBS (mock)-infected mice were analysed by Student’s *t*-test with IBM SPSS Statistics version 20. A value of P < 0.05 was considered statistically significant. The statistically significant differences are indicated with * (P < 0.05) or ** (P < 0.001).

---

**Table S1.** The changed nucleotide sequences in cold-adapted live attenuated SARS-CoV-2 vaccine strain (SARS-CoV-2/human/Korea/CNUHV03-CA22°C/2020)

| Gene name |  |  | Changed nucleotide sequences | The number of changed sequence |
| --- | --- | --- | --- | --- |
| ORF1ab | ORF1a | nsp2 | Deletion : GGTCATGTT(244~252) (NS), A253G (NS), G698A (NS), T2271C (S) | 12 [11 (NS) + 1 (S)]/1923 |
|  |  | nsp3 | G3288A(S), A3550G (NS), C3855T (S), G4754A (NS), G5765A (NS), G6126A (S), G6240A (S), C6933T (S), C7115T (NS). C7311T (S), A7419G (S), C8031T (S), C8174T (NS), C8258T (NS) | 14 [6 (NS) + 8 (S)]/5835 |
|  |  | nsp4 | A8611G (NS), C8914T (NS), G8953A (NS) | 3 (NS)/1500 |
|  |  | nsp6 | T10818G (NS), T10827A (NS), T10851G (S) | 3 [2(NS)+ 1 (S)]/870 |
|  |  | nsp7 | A11777C (NS) | 1 (NS)/249 |
|  |  | nsp10 | C12882T (S), G13050A (S) | 2 (S)/417 |
|  | ORF1b | RNA-dependent RNA polymerase | C13822T (NS), G14417A (NS), C15464T (NS) | 3 (NS)/2796 |
|  |  | helicase | C16810T (NS), C17508T (S), G17545A (NS) | 3[2(NS)+ 1 (S)]/1803 |
|  |  | 3'-to-5' exonuclease | C18349T (S) | 1(S)/1581 |
|  |  | endoRNase | T19506G (S) | 1(S)/1038 |
|  |  | 2'-0-ribose methyl transferase | G21113C (NS), C21168T (S), C21175T (NS) | 3 [2(NS)+1 (S)]/894 |
| S |  |  | G113A (NS), C257T (NS), G287A (NS), G451A (NS), C2511T (S), T2902G (NS), C3003T (S) | 7 [5 (NS)+ 2 (S)]/3822 |
| M |  |  | C537T (S) | 1 (S)/669 |
| ORF7a |  |  | C141T (S), A201G (NS) | 2 [1(NS)+ 1(S)]/366 |
| ORF8 |  |  | C63T (S) | 1 (S)/366 |
| N |  |  | N243T (S), T1082A (NS) | 2[1(NS)+1 (S)]/1260 |
| Total nucleotides |  |  |  | 59 [37 (NS)+22 (S)]/29874 including non-coding nucleotides |

We compared the nucleotide sequences of cold-adapted live attenuated SARS-CoV-2 vaccine strain (SARS-CoV-2/human/Korea/CNUHV03-CA22°C/2020) with those in wild-type SARS-CoV-2: (SARS-CoV-2/human/Korea/CNUHV03/2020), which was used for developing cold-adapted live attenuated vaccine strain in Vero cells.
(S): (synonymous substitution). (NS): (nonsynonymous substitution)

**Table S2.** Changed amino acid sequences in cold-adapted live attenuated SARS-CoV-2 vaccine strain (SARS-CoV-2/human/Korea/CNUHV03-CA22°C/2020)

| Protein name |  |  | Changed amino acid sequences | The number of changed sequence |
| --- | --- | --- | --- | --- |
| ORF1ab polyprotein | ORF1a polyprotein | nsp2 | Deletion : GHV(82~84), M85V, G233E | 5/641 |
|  |  | nsp3 | N1184D, G1585D, S1922N, A2372V, P2725L, A2753V | 6/1945 |
|  |  | nsp4 | S2871G, P2972S, G2985R | 3/500 |
|  |  | nsp6 | F3606L, N3609K | 2/290 |
|  |  | nsp7 | D3926A | 1/83 |
|  | ORF1b | RNA-dependent RNA polymerase | R4608W, S4806N, A5155V | 3/932 |
|  |  | helicase | L5604F, V5849I | 2/601 |
|  |  | 2'-O-ribose methyl transferase | C7038S, P7059S | 2/298 |
| S protein |  |  | C38Y, S86F, G96E, G151S, S968A | 5/1274 |
| ORF7a |  |  | Y67C | 1/122 |
| N protein |  |  | I361K | 1/420 |
| Total amino acids |  |  |  | 31/9755<br>(including non-changed ORFs) |

We compared the amino acid sequences of cold-adapted live attenuated SARS-CoV-2 vaccine strain (SARS-CoV-2/human/Korea/CNUHV03-CA22°C/2020) with those in wild-type SARS-CoV-2 (SARS-CoV-2/human/Korea/CNUHV03/2020), which was used for developing cold-adapted live attenuated vaccine strain in Vero cells.
Single letter abbreviation name of amino acid: Glycine(G), Alanine(A), Valine(V), Cysteine(C), Proline(P), Leucine(L), Isoleucine(I), Methionine(M), Tryptophan(W), Phenylalanine(F), Serine(S), Tyrosine(Y), Asparagine(N), Lysine(K), Arginine(R), Histidine(H), Aspartic Acid(D), Glutamic Acid(E)

**Table S3.**
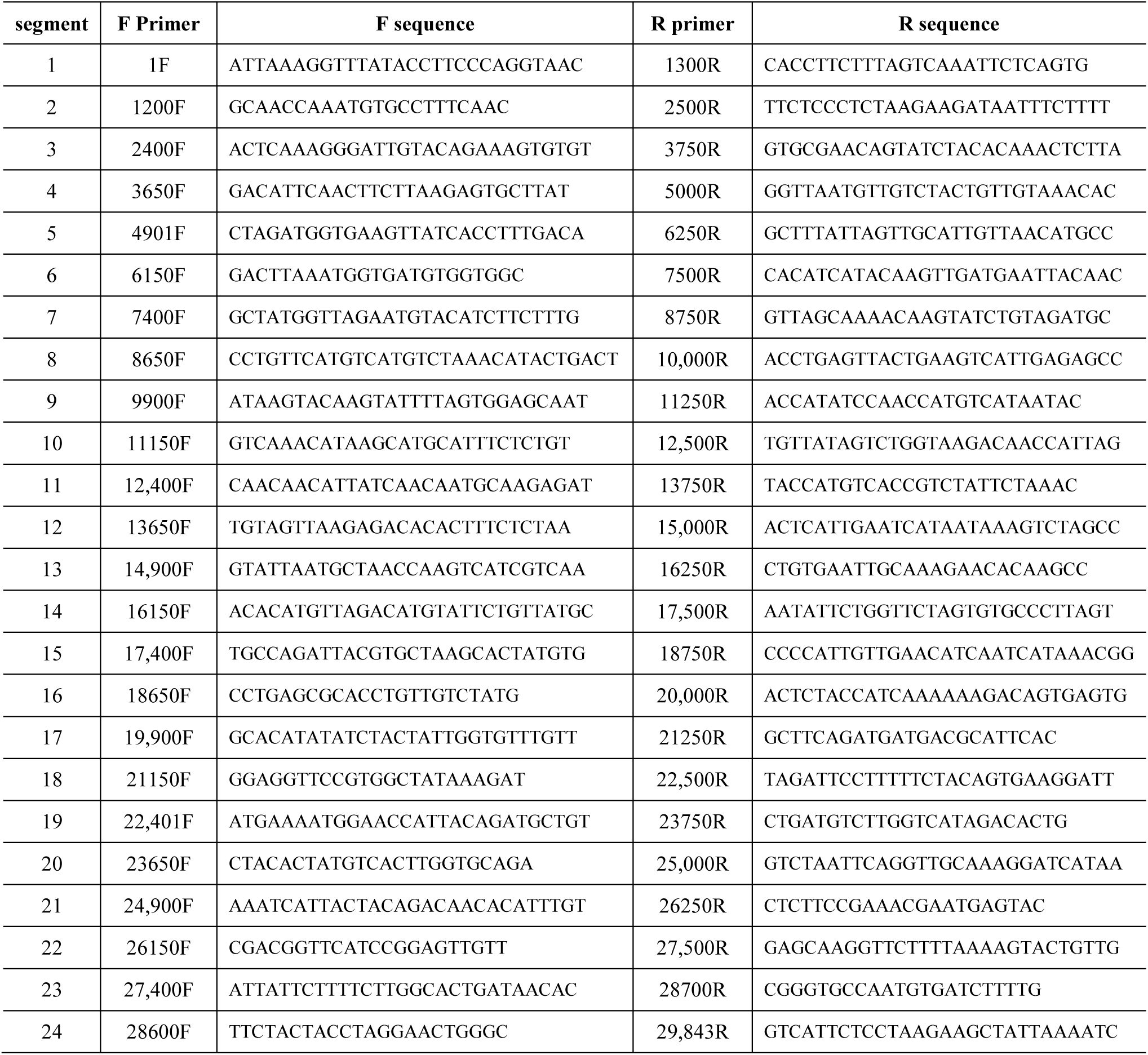
Primers used for PCR amplification of the gene segments in cold-adapted live attenuated SARS-CoV-2 vaccine strain (SARS-CoV-2/human/Korea/CNUHV03-CA22°C/2020)

**Table S4.**
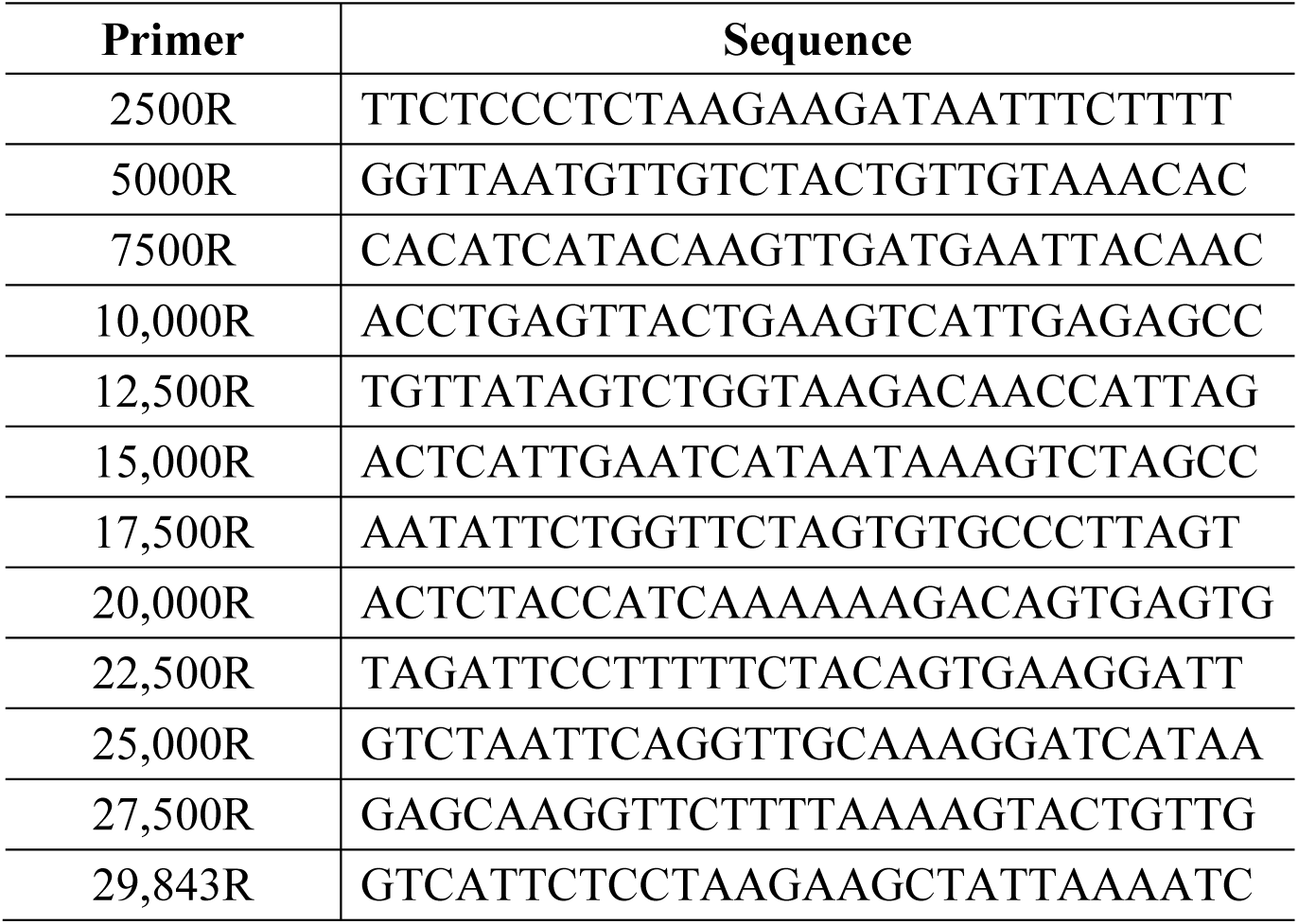
Primers for the synthesis of cDNA for cold-adapted live attenuated SARS-CoV-2 vaccine strain (SARS-CoV-2/human/Korea/CNUHV03-CA22°C/2020)

**Fig. S1.**
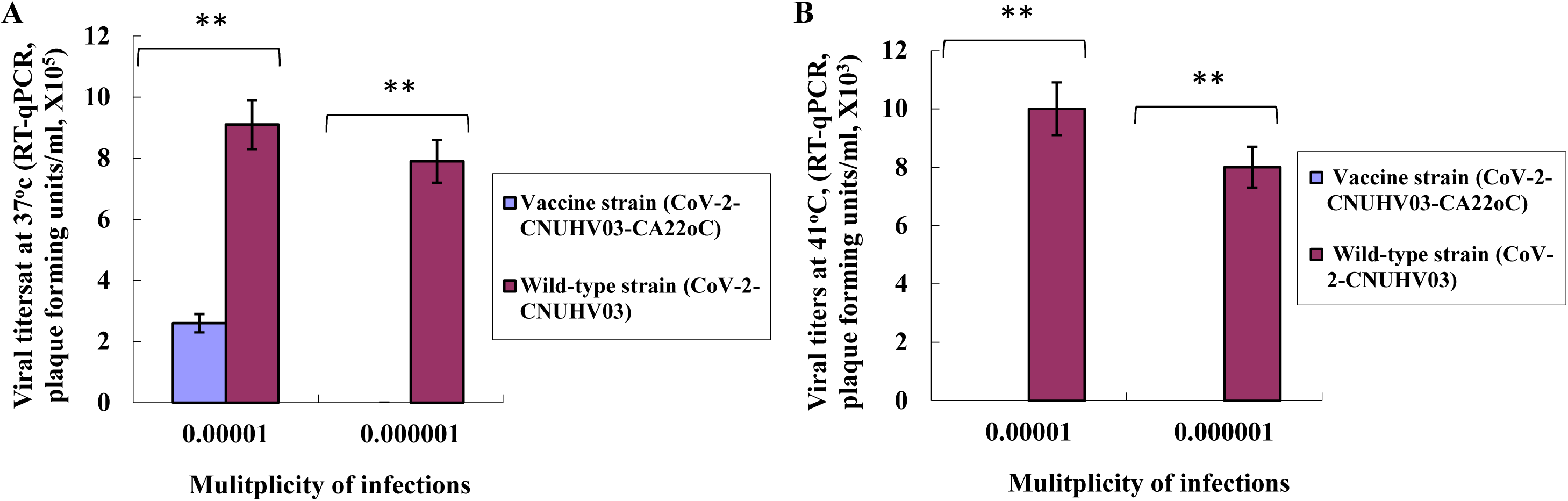
Temperature sensitivity of cold-adapted SARS-CoV-2 vaccine strain. Vero cells were infected with 0.00001 or 0.000001 m.o.i of cold-adapted SARS-CoV-2 vaccine strain (CoV-2-CNUHV03-CA22°C) or wild-type SARS-CoV-2 (CoV-2-CNUHV03) and were incubated at 37°C or 41°C for 3 days. Viral titers in cell supernatants were determined by RT-qPCR using SARS-CoV-2 N primers and probe. Viral detection limit is 10 pfu. **A**, viral titers at 37°C; **B**, viral titers at 41°C. **\*p<0.05, **p<0.001**

**Fig. S2.**
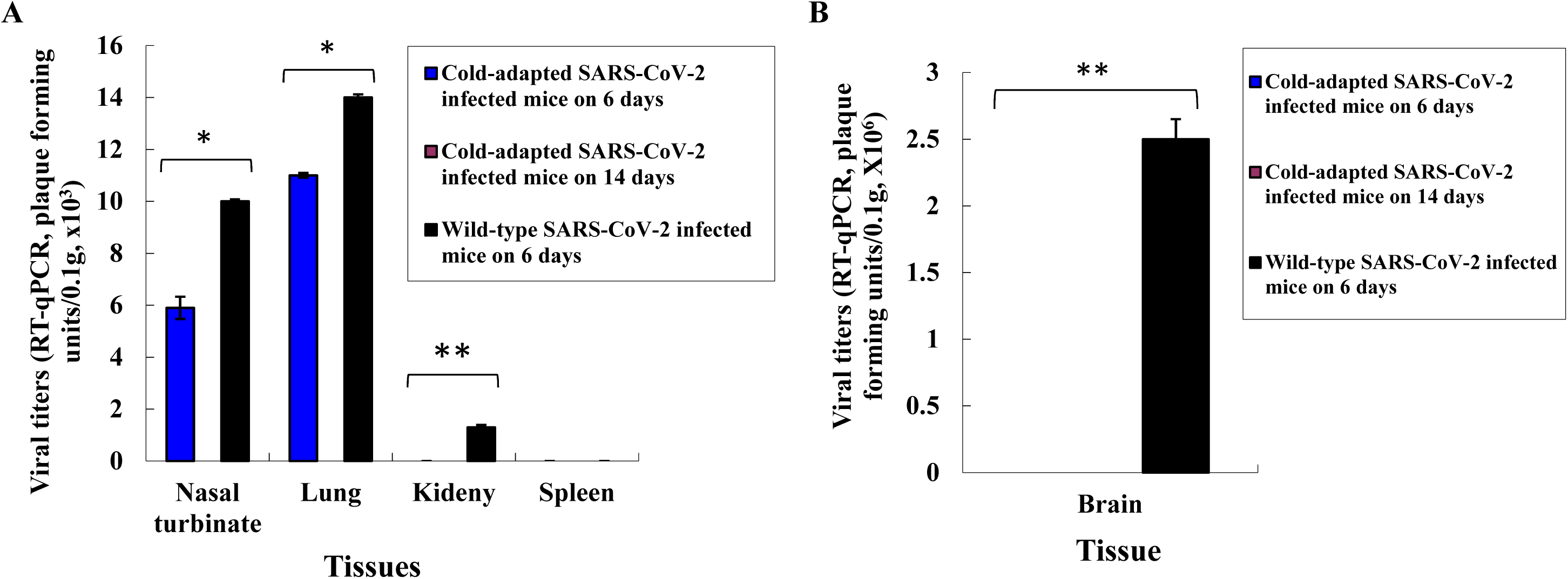
Viral titers in tissues of hACE2 transgenic mice infected with cold adapted SARS-CoV-2. Viral titers in tissues of Fig. 1c were quantified RT-qPCR. Viral titers are the mean of 3 tissues ± standard deviations. Viral detection limit is 10 pfu. **A**, viral titers in nasal turbinate, lung, kidney, and spleen, **B**, viral titers in brain. **\*p<0.05, **p<0.001**

**Fig. S3.**
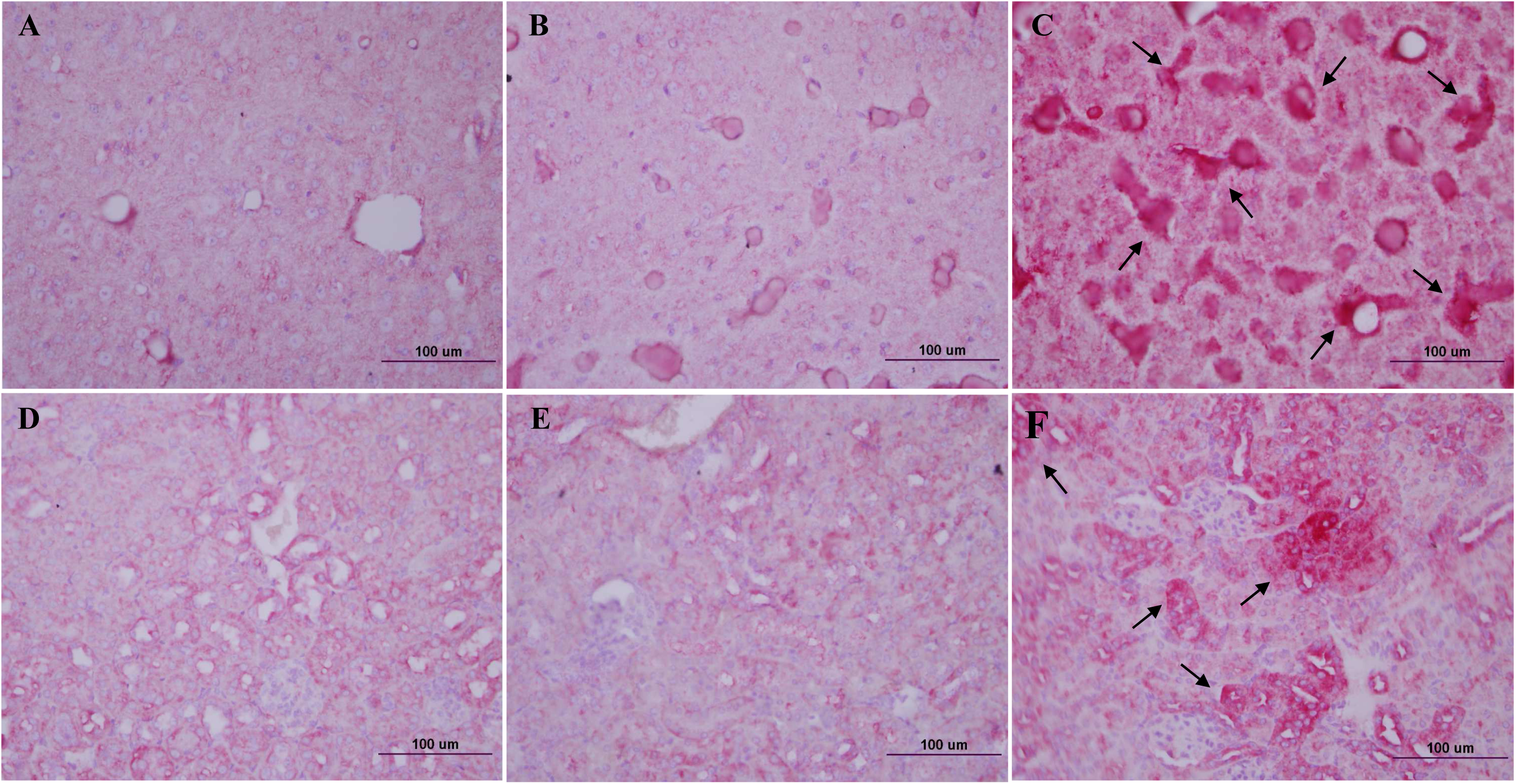
Antigen staining in brain and kidney of hACE2 transgenic mice infected with cold adapted SARS-CoV-2. Brain and kidney tissues of Fig. 1C (on day 6 p.i.) were stained with SARS-CoV-2 NP antibody (X400). **A**, brain tissue of PBS-mock mouse; **B**, brain tissue of mouse i.n. infected with cold adapted SARS-CoV-2 (CoV-2-CNUHV03-CA22°C) (2X10^4^ pfu); **C,** brain tissue of mouse i.n. infected with wild-type SARS-CoV-2 (CoV-2-CNUHV03) (2X10^4^ pfu); **D**, kidney tissue of PBS-mock mouse; **E**, kidney tissue of mouse i.n. infected with cold adapted SARS-CoV-2 (CoV-2-CNUHV03-CA22°C) (2X10^4^ pfu); **F,** kidney tissue of mouse i.n. infected with wild-type SARS-CoV-2 (CoV-2-CNUHV03) (2X10^4^ pfu). Arrow: positive antigen staining.

**Fig. S4.**
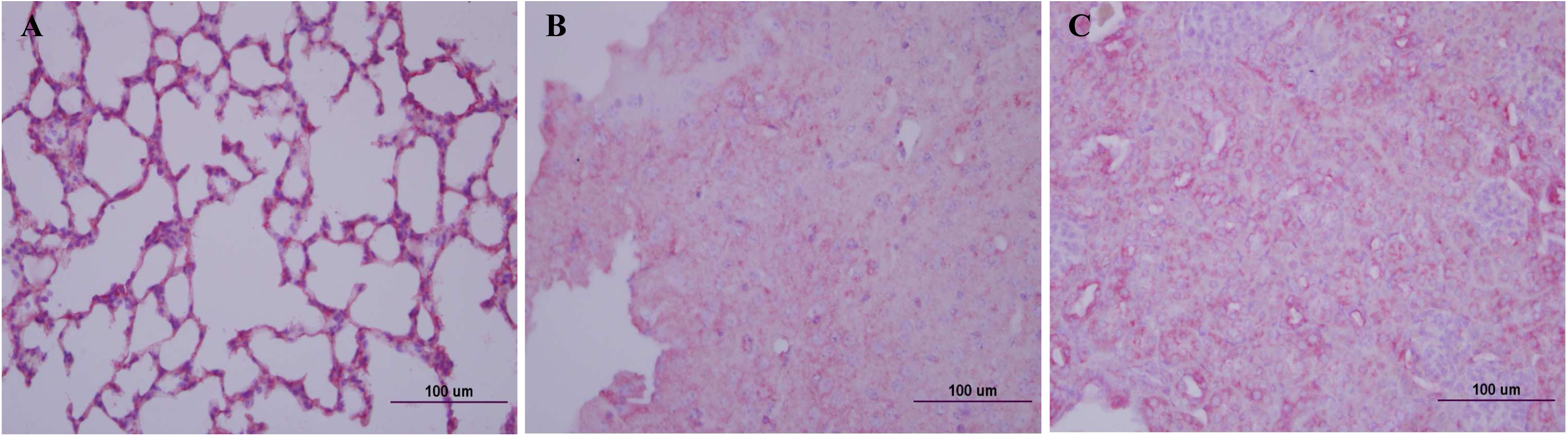
Antigen staining in lung, brain and kidney of hACE2 transgenic mice infected with cold adapted SARS-CoV-2 on day 14 p.i. Lung, brain and kidney tissues of Fig. 1 (on day 14 p.i.) were stained with SARS-CoV-2 NP antibody (X400). **A**, lung tissue of mouse i.n. infected with cold adapted SARS-CoV-2 (CoV-2-CNUHV03-CA22°C) (2X10^4^ pfu); **B**, brain tissue of mouse i.n. infected with cold adapted SARS-CoV-2 (CoV-2-CNUHV03-CA22°C) (2X10^4^ pfu); **C,** kidney tissue of mouse i.n. infected with cold adapted SARS-CoV-2 (CoV-2-CNUHV03-CA22°C) (2X10^4^ pfu).

**Fig. S5.**
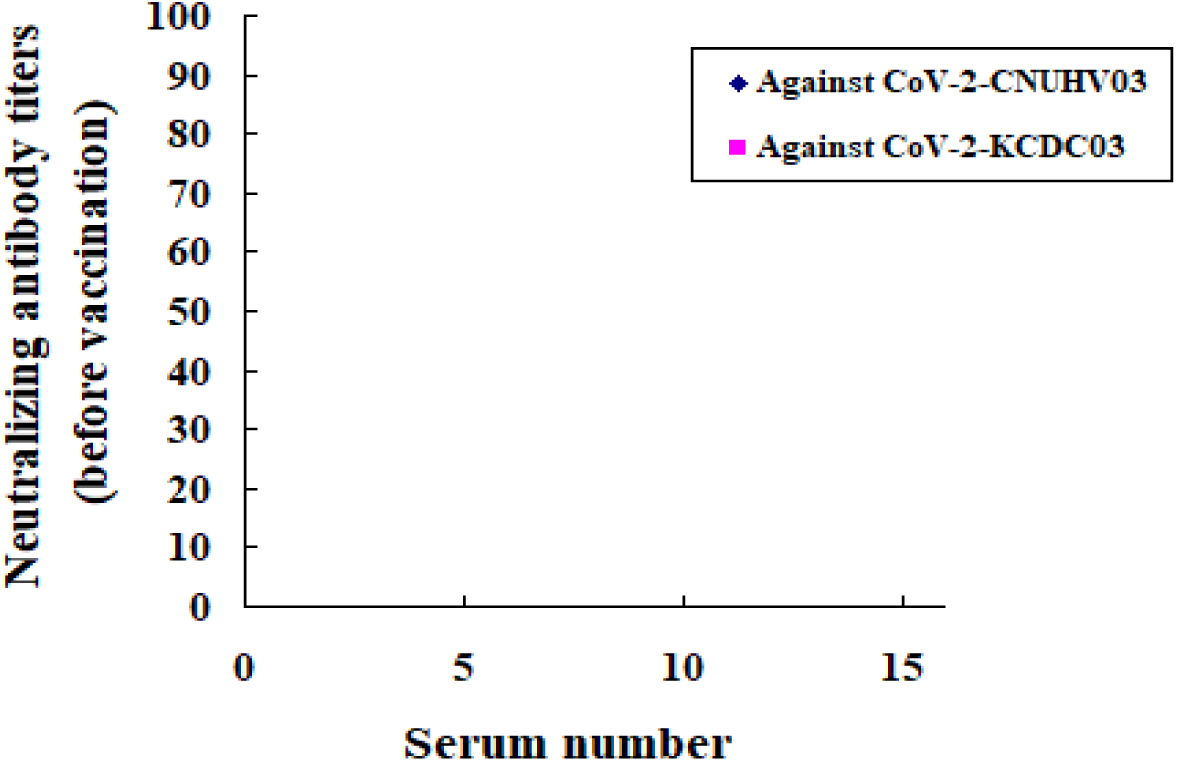
Neutralizing antibody titers in sera collected from hACE2 transgenic mice before immunization. Sera were collected from K18-ACE2 mice (n=16) before immunization, and their neutralizing antibody titers were determined against wild-type SARS-CoV-2 viruses, CoV-2-CNUHV03 and CoV-2-KCDC03 in Vero cells. Detection limit of neutralizing antibody is 10.

**Fig S6.**
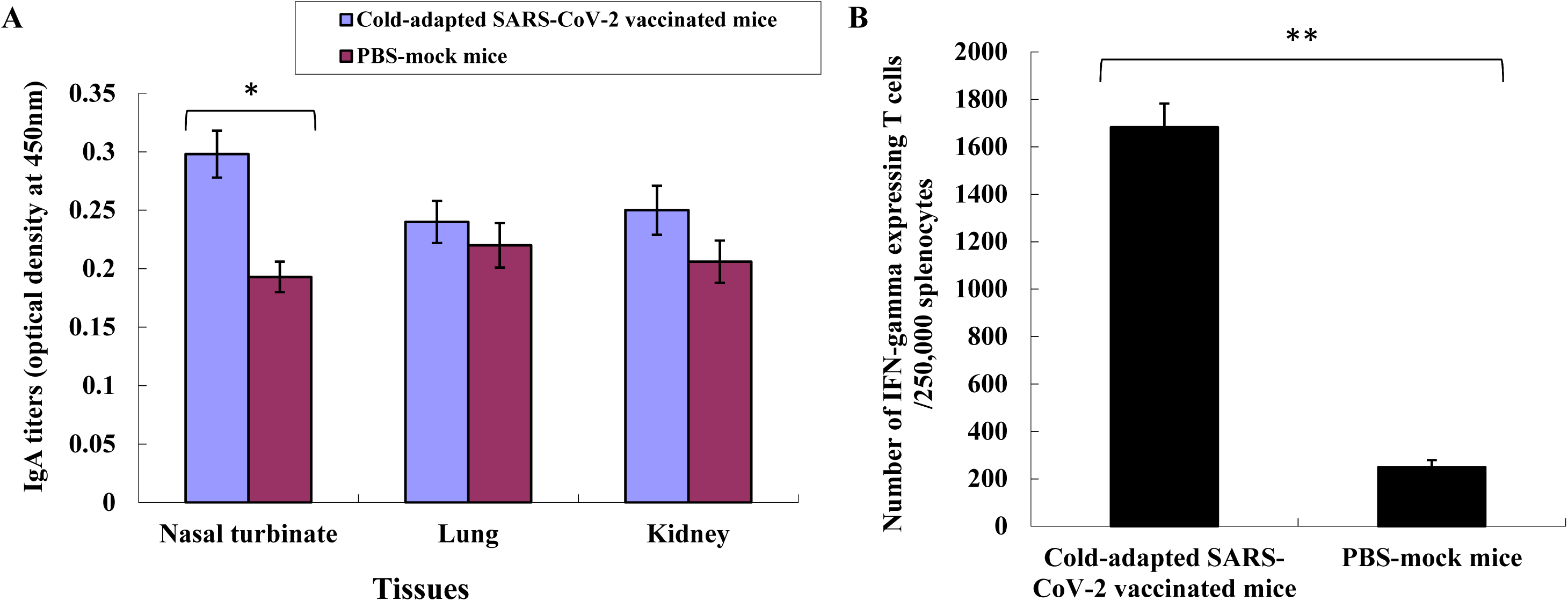
IgA antibody titers and the number of T lymphocytes expressing IFN-γ in the immunized hACE2 transgenic mice. The immunized K18-ACE2 mice (n=3) with 2X10^4^ pfu of CoV-2-CNUHV03-CA22°C were euthanized to collect tissues (nasal turbinate, lungs, kidneys, spleens) on 19 days after immunization. Nasal turbinate, lungs, and kidneys were homogenized in PBS and were used for detection of IgA antibody by ELISA using the purified and inactivated SARS-CoV-2 antigen (SARS-CoV-2/human/Korea/CNUHV03/2020). Lymphocytes were collected from spleens and used to determine the number of T lymphocytes expressing IFN-γ by ELISPOT assay. Data are the mean of 3 tissues ± standard deviations. **A,** IgA antibody titers; **B,** number of T lymphocytes expressing IFN-γ. **\*p<0.05, **p<0.001**

**Fig. S7.**
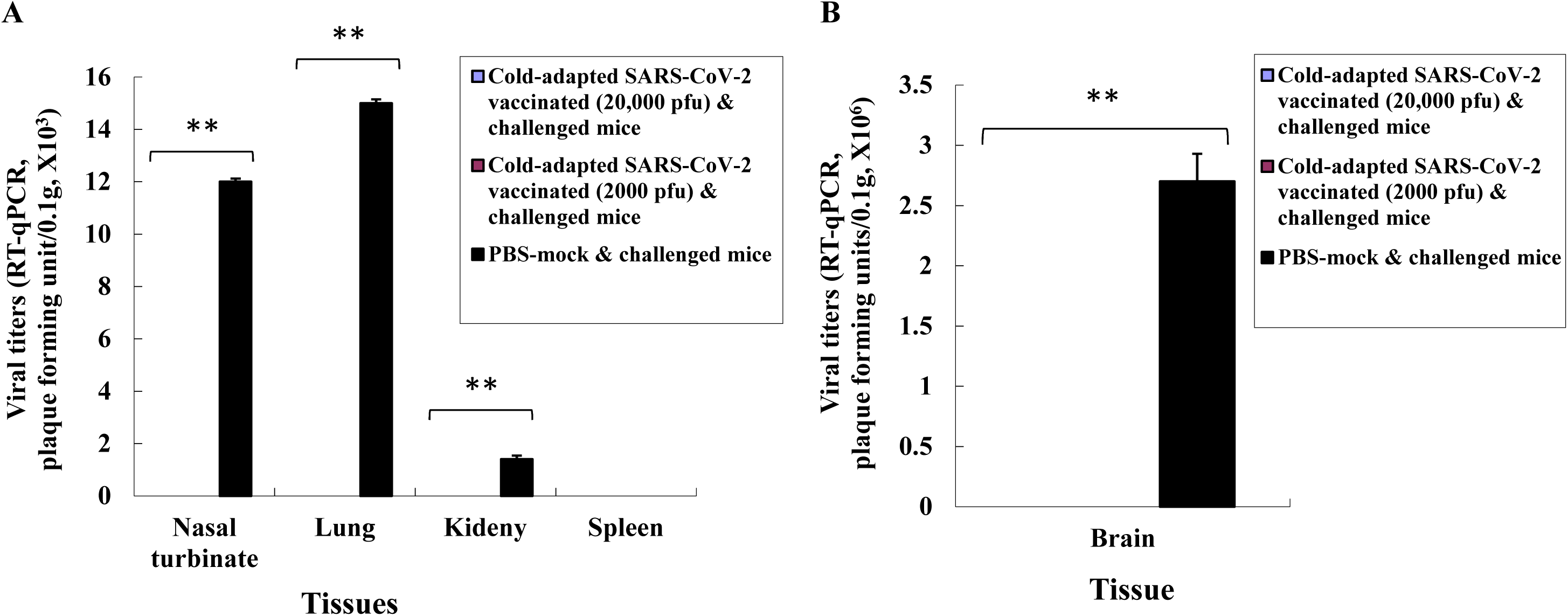
Viral titers in tissues of the immunized and challenged hACE2 transgenic mice. Viral titers in tissues of Fig.3E were quantified RT-qPCR. Viral titers are the mean of 3 tissues ± standard deviations. Viral detection limit is 10 pfu. **A**, viral titers in nasal turbinate, lung, kidney, and spleen; **B**, Viral titer in brain. **\*p<0.05, **p<0.001**

**Fig. S8.**
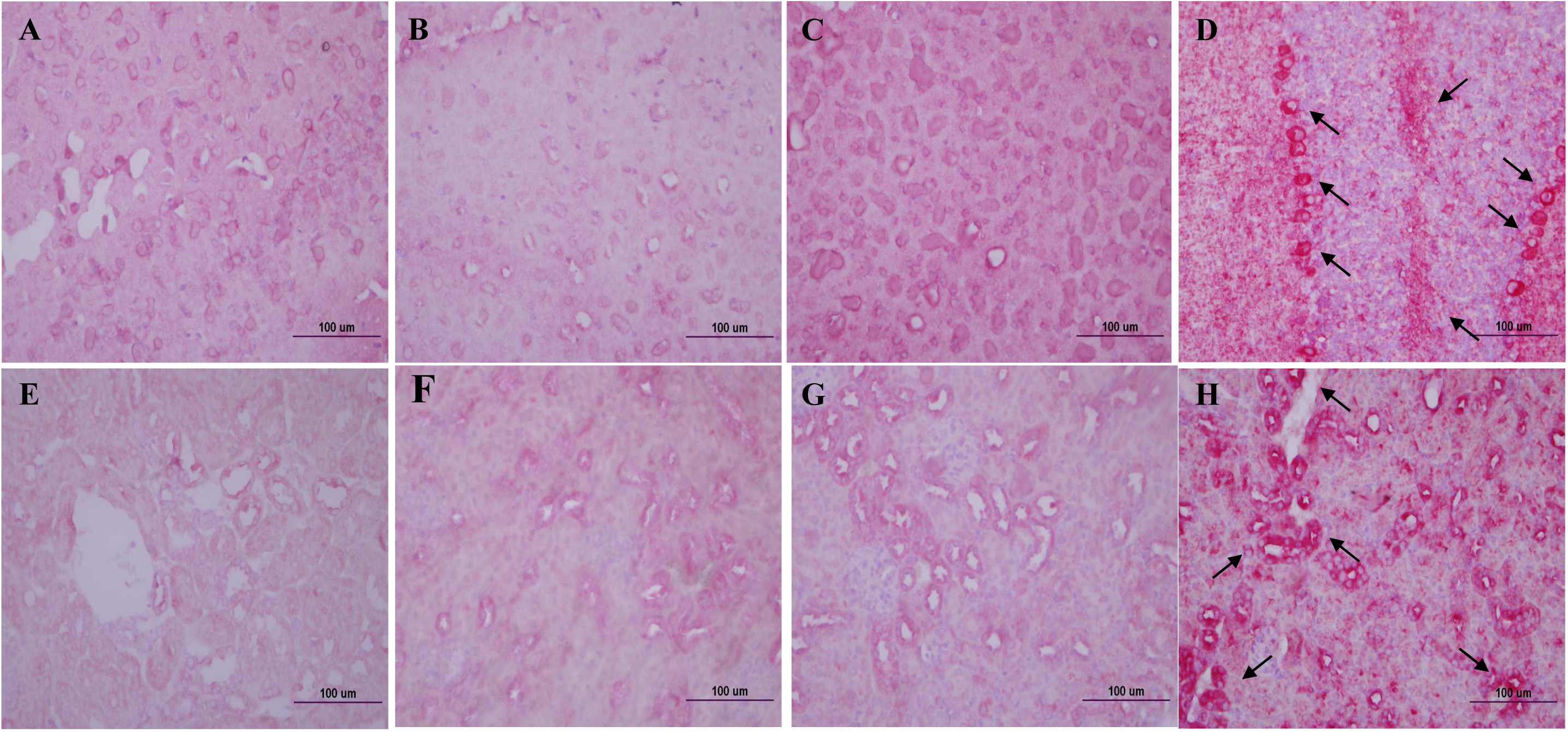
Antigen staining in brain and kidney of the immunized and challenged hACE2 transgenic mice. Brain and kidney tissues of challenged K18-ACE2 mice (Fig. 3E) were stained with SARS-CoV-2 NP antibody (X400). **A**, brain tissue of PBS-mock mouse; **B**, brain tissue of vaccinated mouse with CoV-2-CNUHV03-CA22°C (2X10^3^ pfu) and challenged with CoV-2-KCDC03 (2X10^4^ pfu) ; **C,** brain tissue of vaccinated mouse with CoV-2-CNUHV03-CA22°C (2X10^4^ pfu) and challenged with CoV-2-KCDC03 (2X10^4^ pfu); **D**, brain tissue of PBS-mock vaccinated mouse challenged with CoV-2-KCDC03 (2X10^4^ pfu); **E**, kidney tissue of PBS-mock mouse; **F,** kidney tissue of vaccinated mouse with CoV-2-CNUHV03-CA22°C (2X10^3^ pfu) and challenged with CoV-2-KCDC03 (2X10^4^ pfu); **G,** kidney tissue of vaccinated mouse with CoV-2-CNUHV03-CA22°C (2X10^4^ pfu) and challenged with CoV-2-KCDC03(2X10^4^ pfu); H, kidney tissue of PBS-mock vaccinated mouse challenged with CoV-2-KCDC03 (2X10^4^ pfu). Arrow: positive antigen staining.

**Fig. S9.**
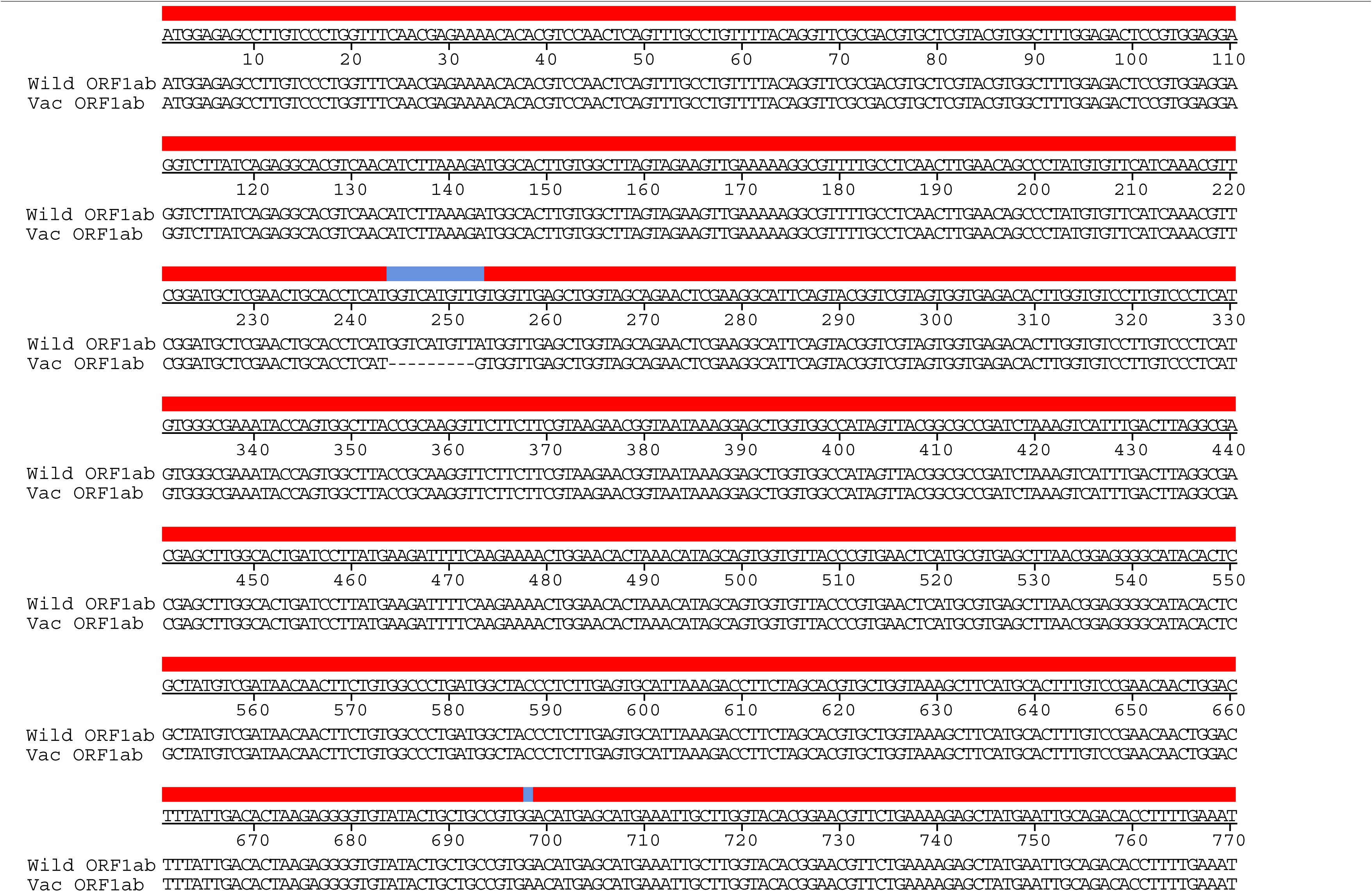
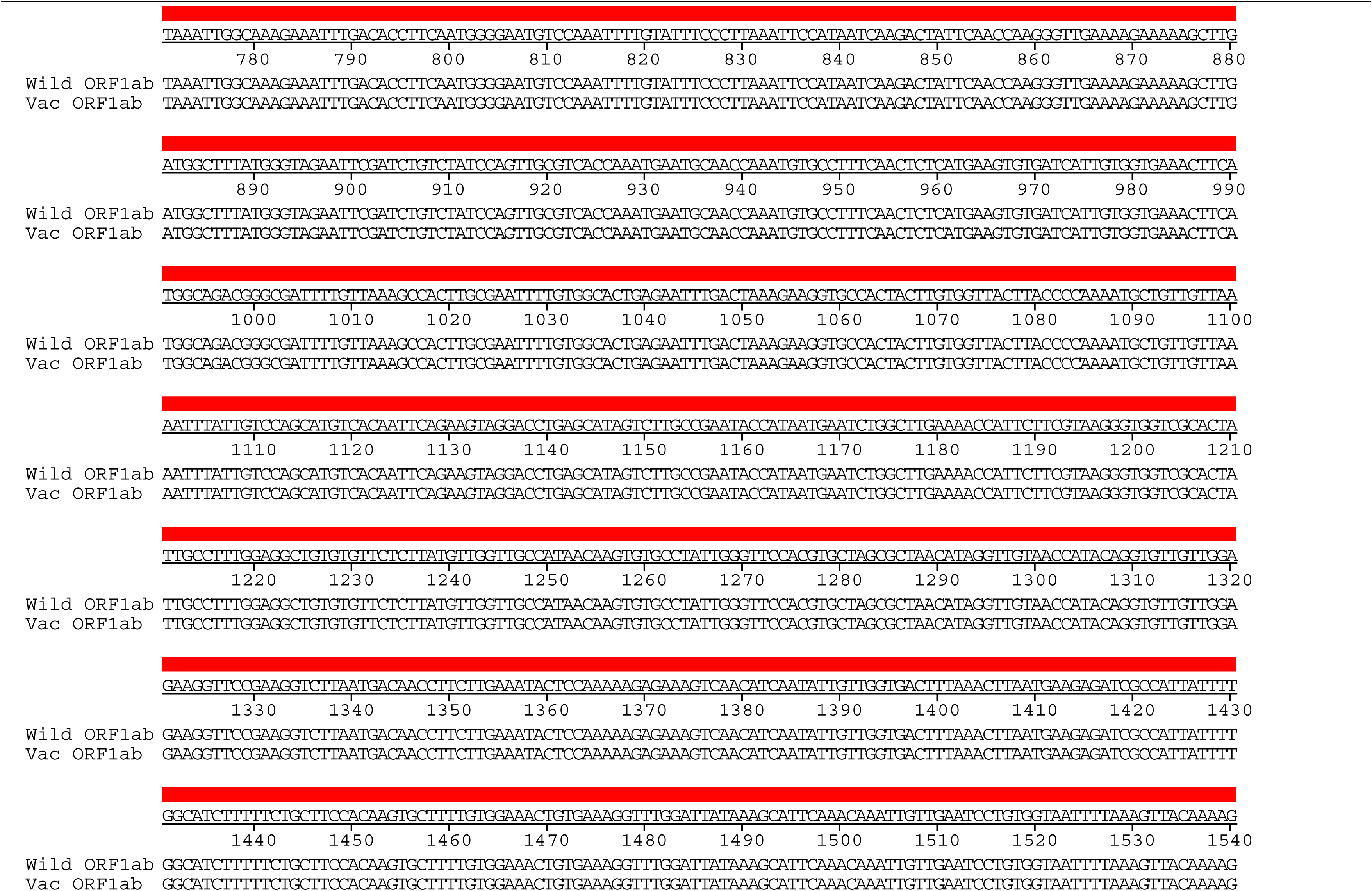
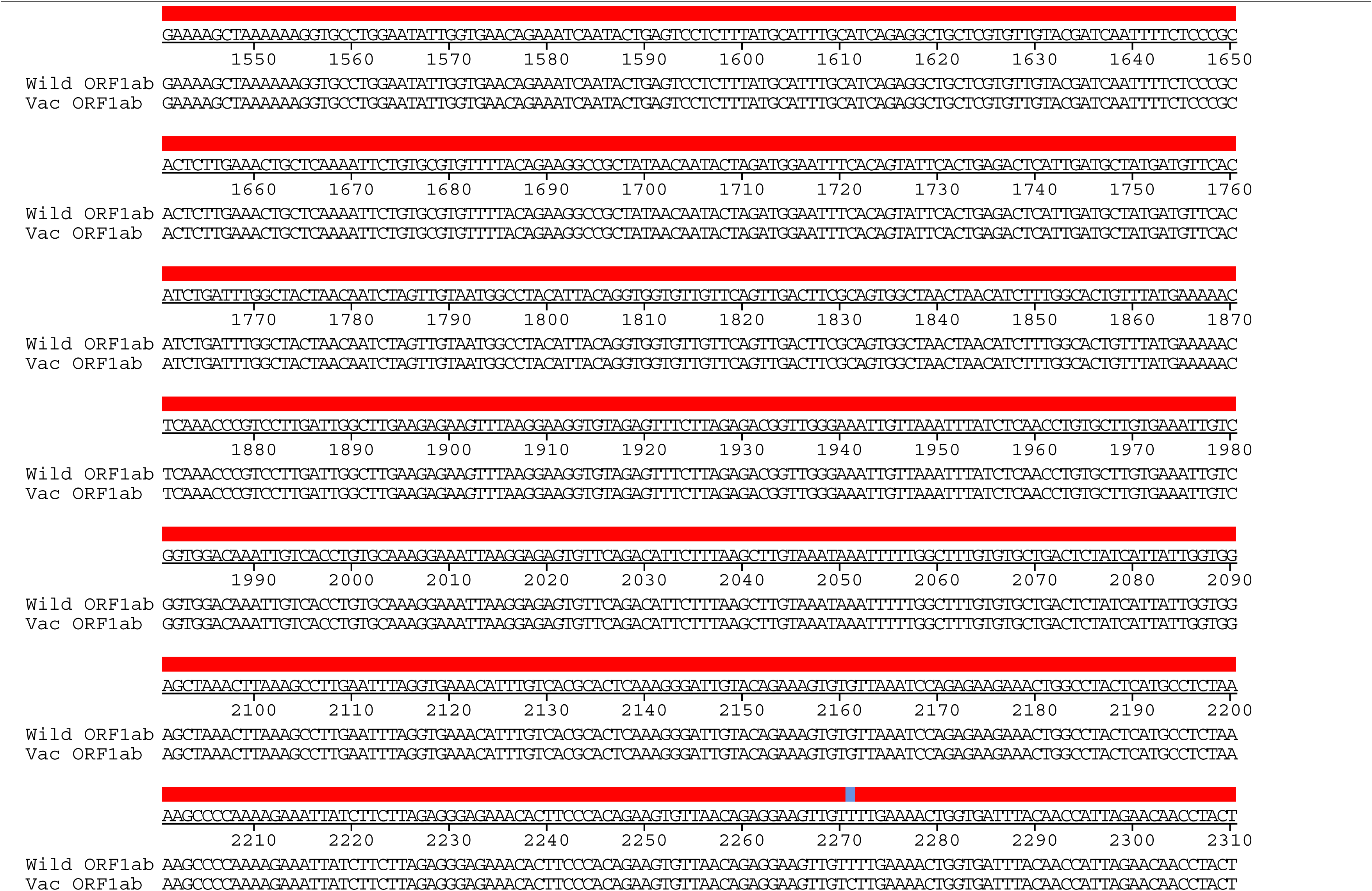
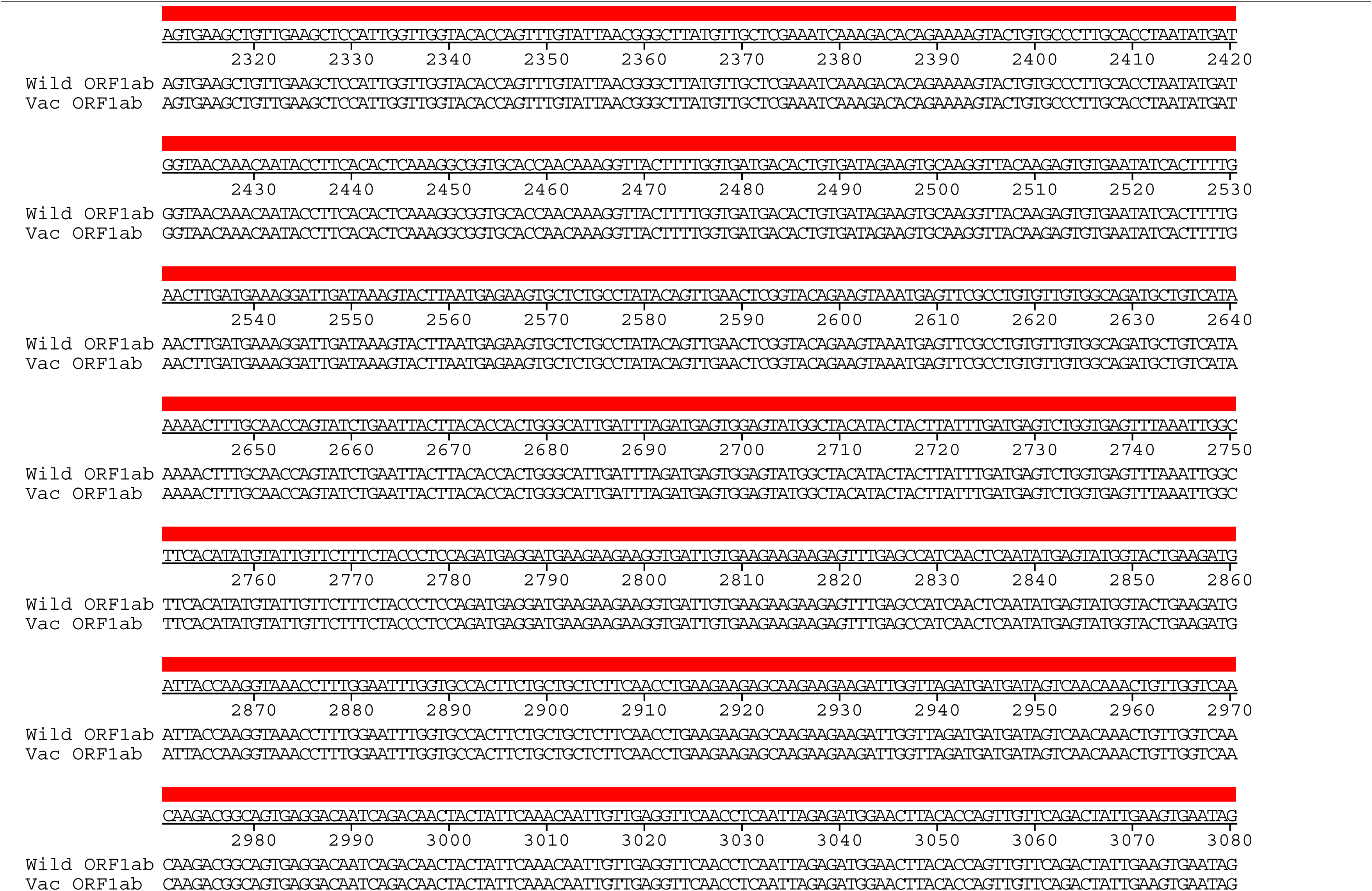
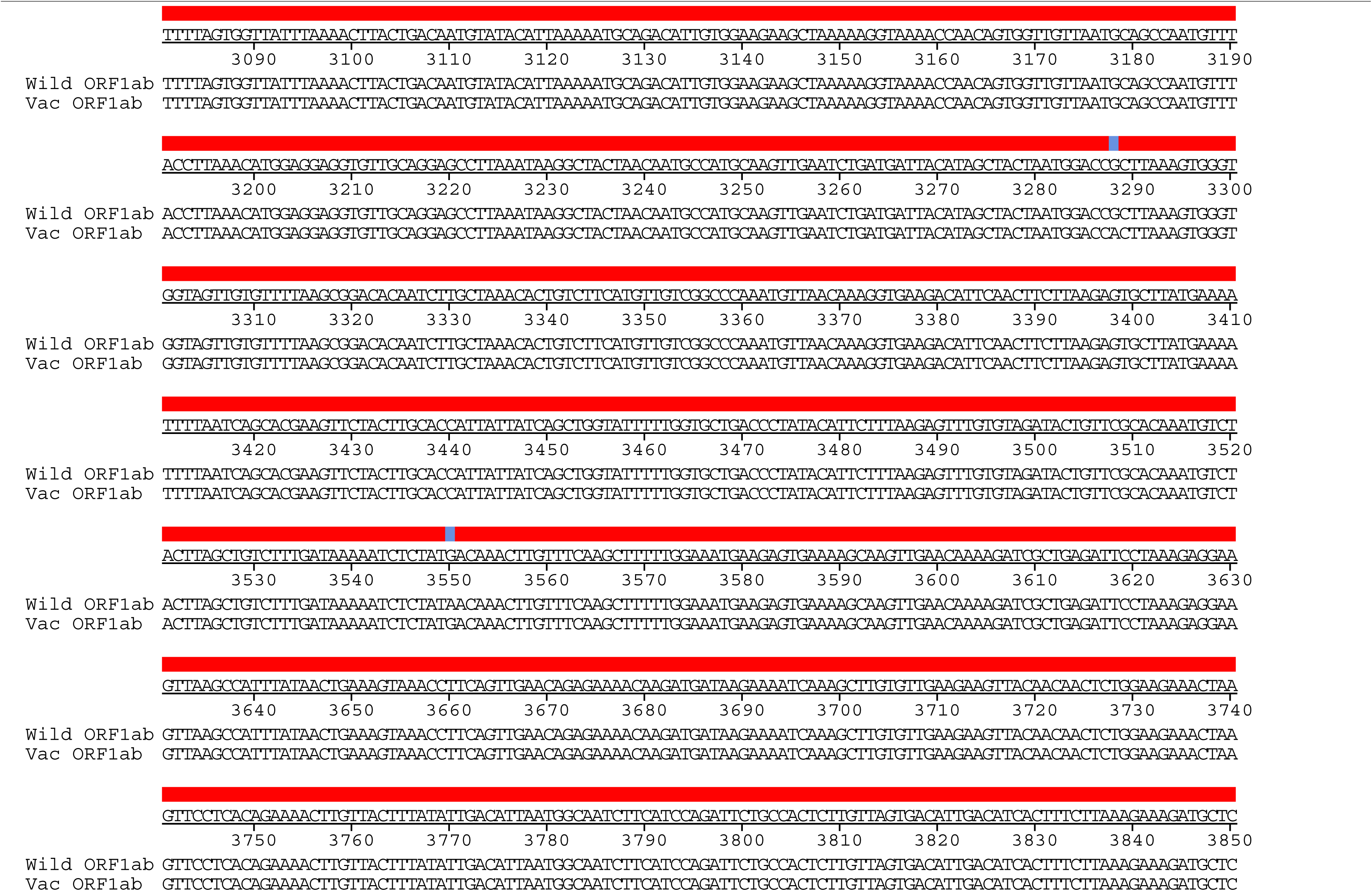
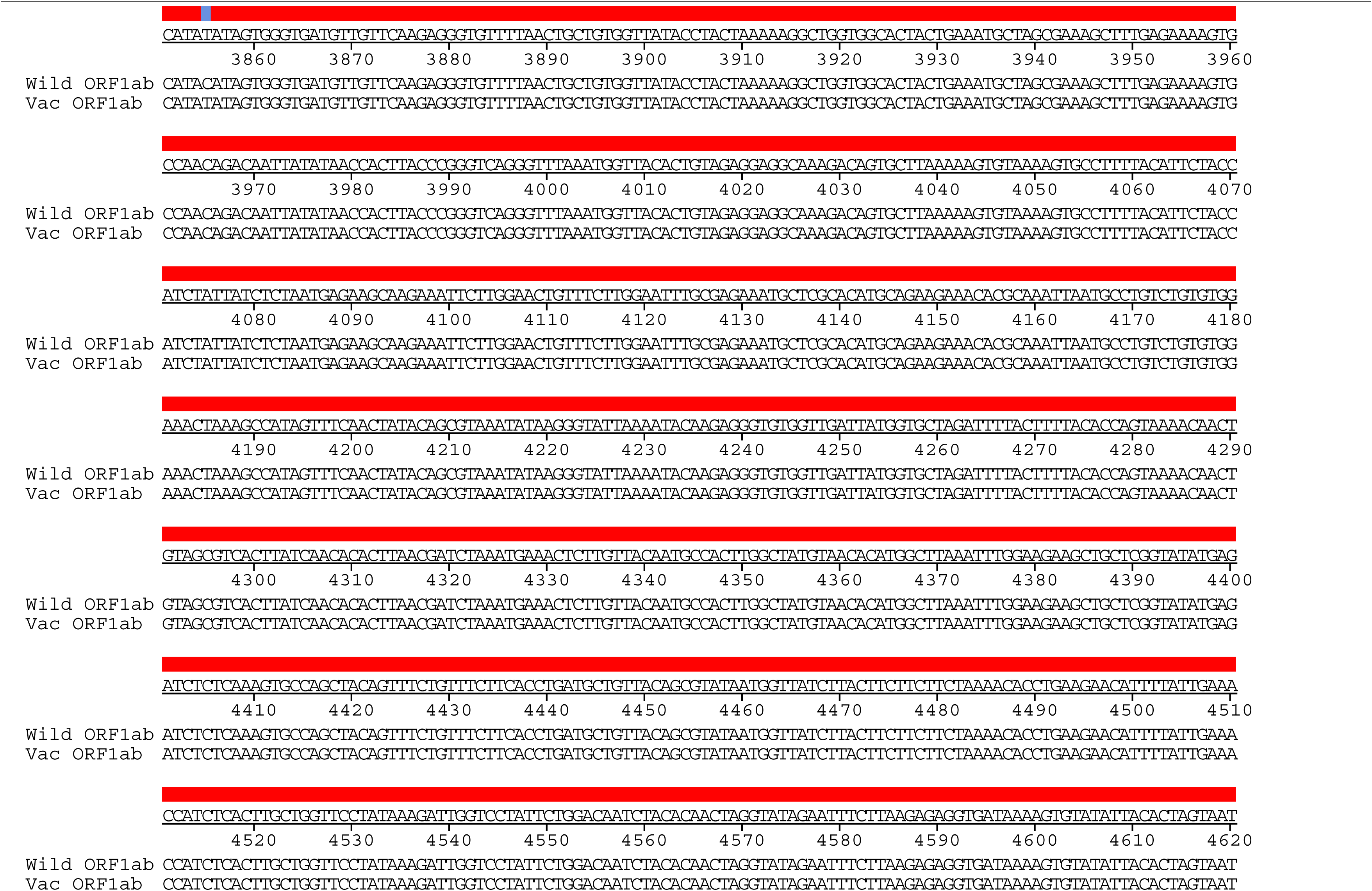
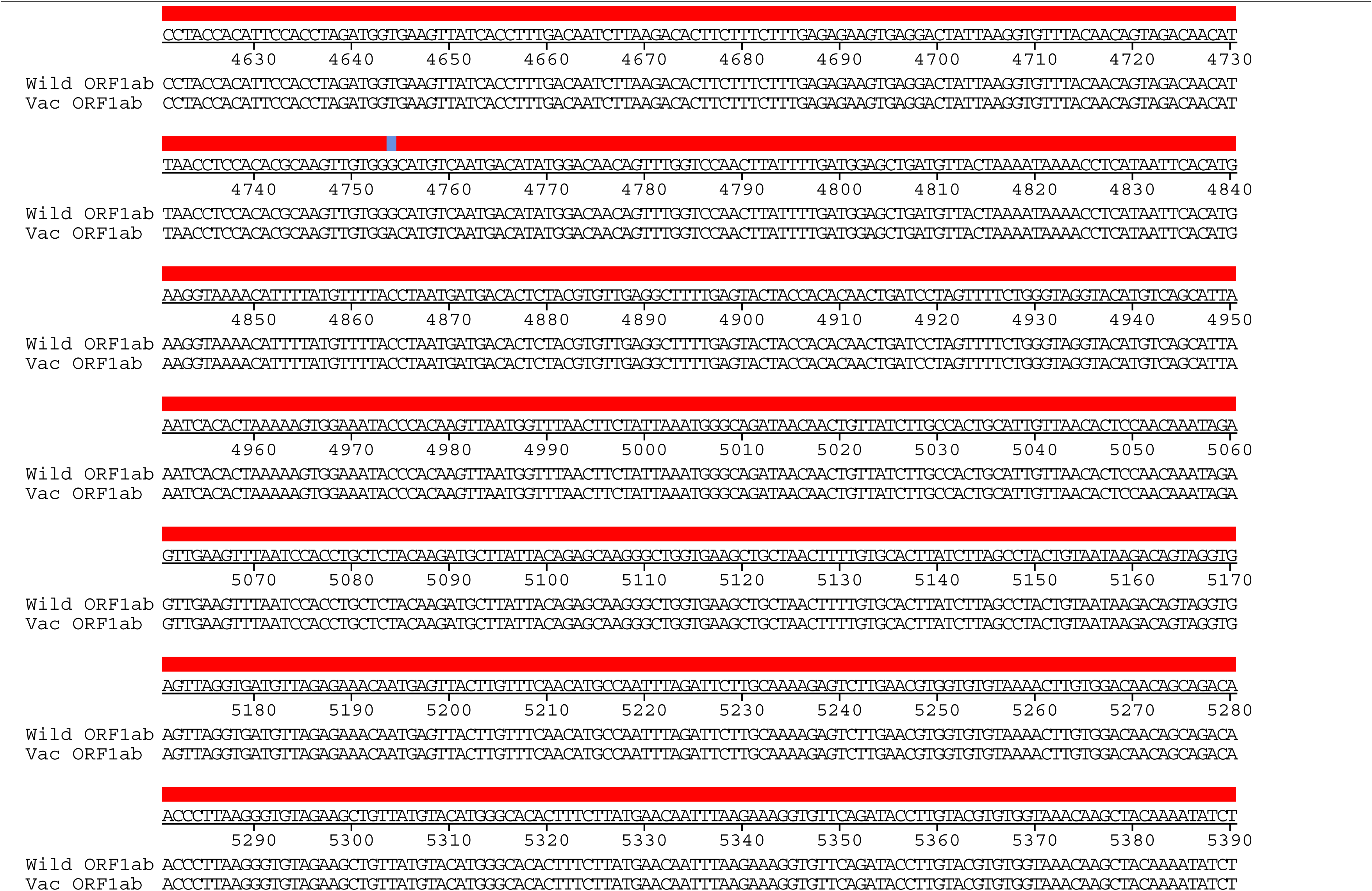
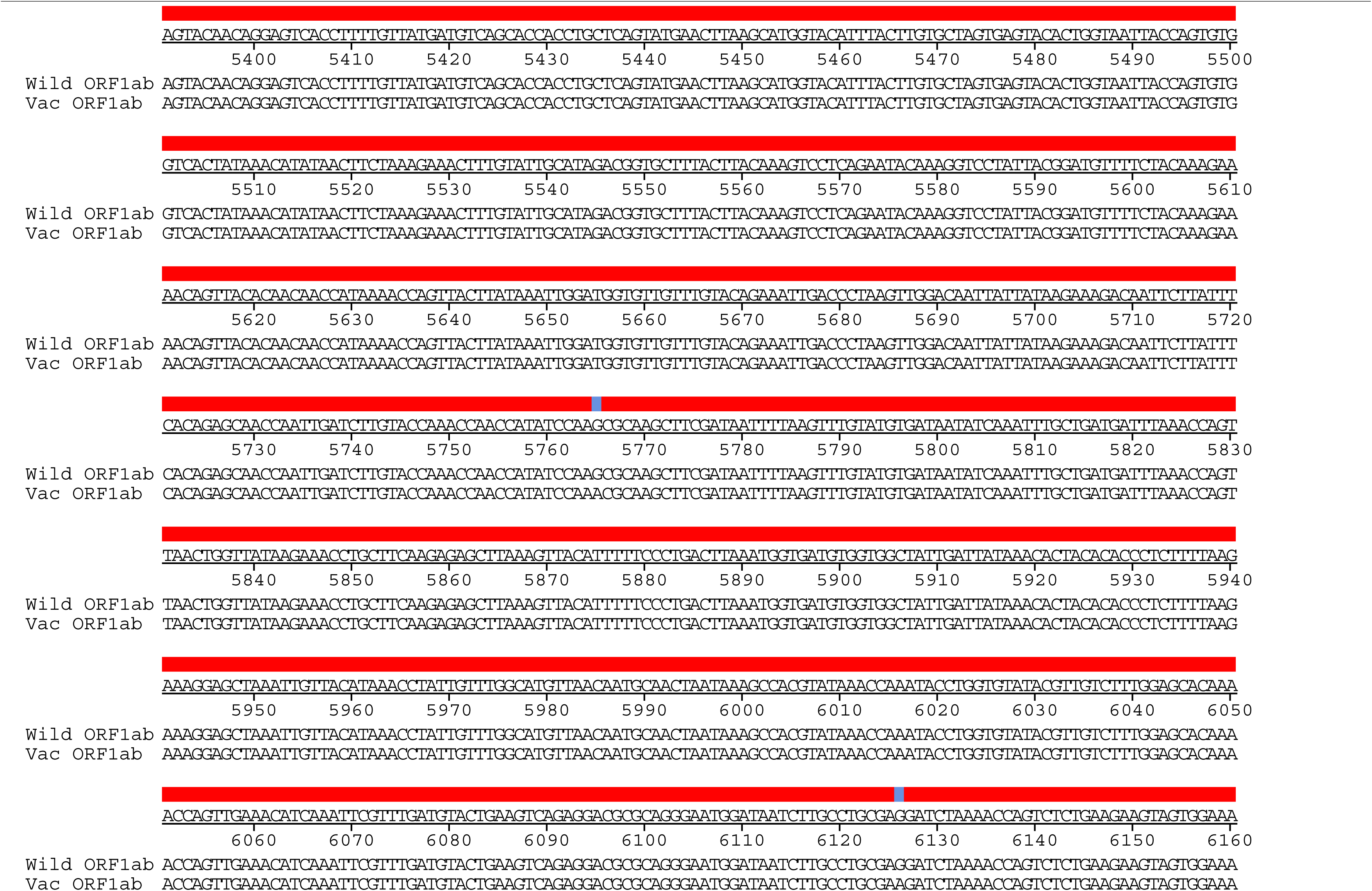
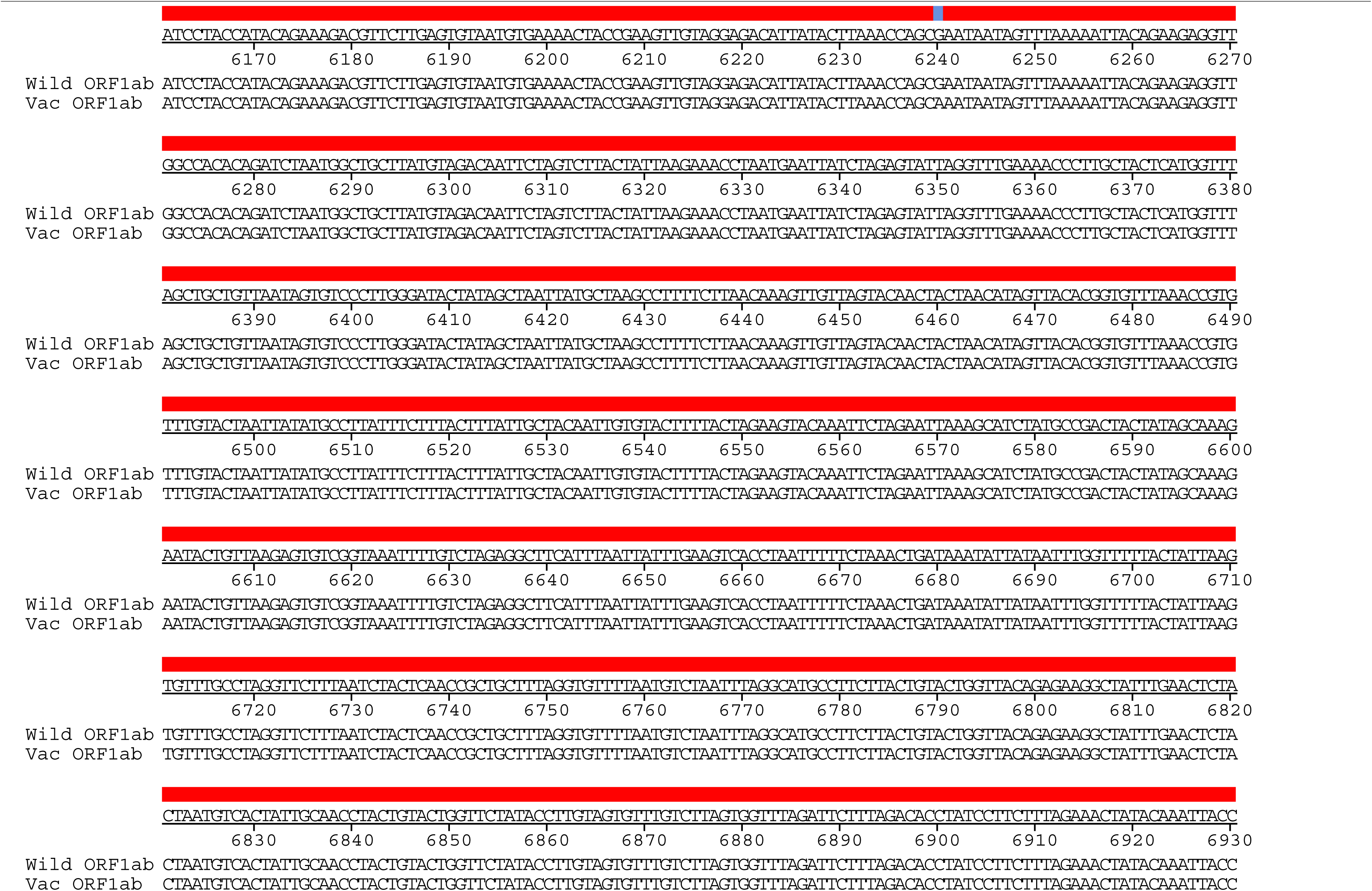
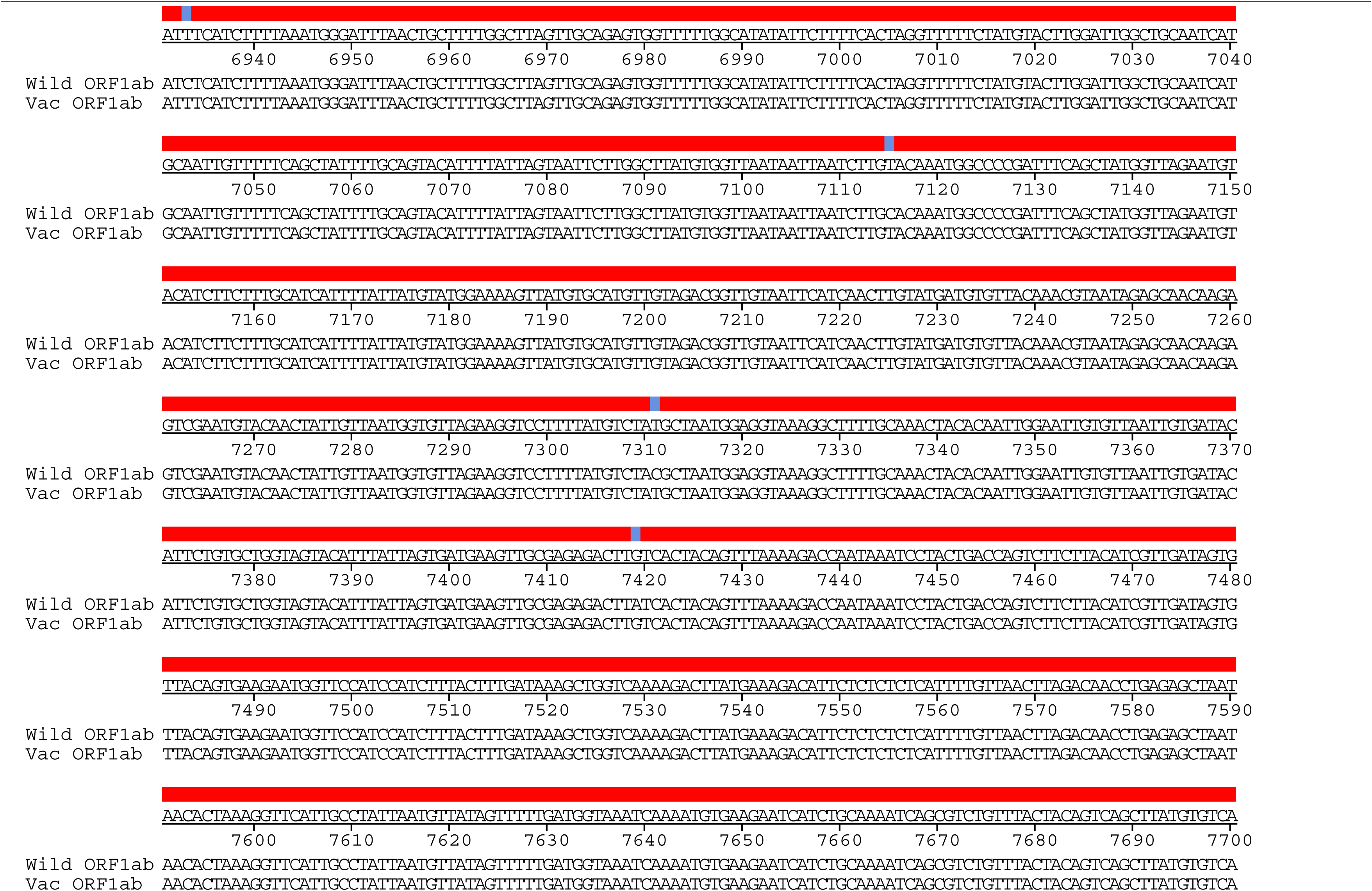
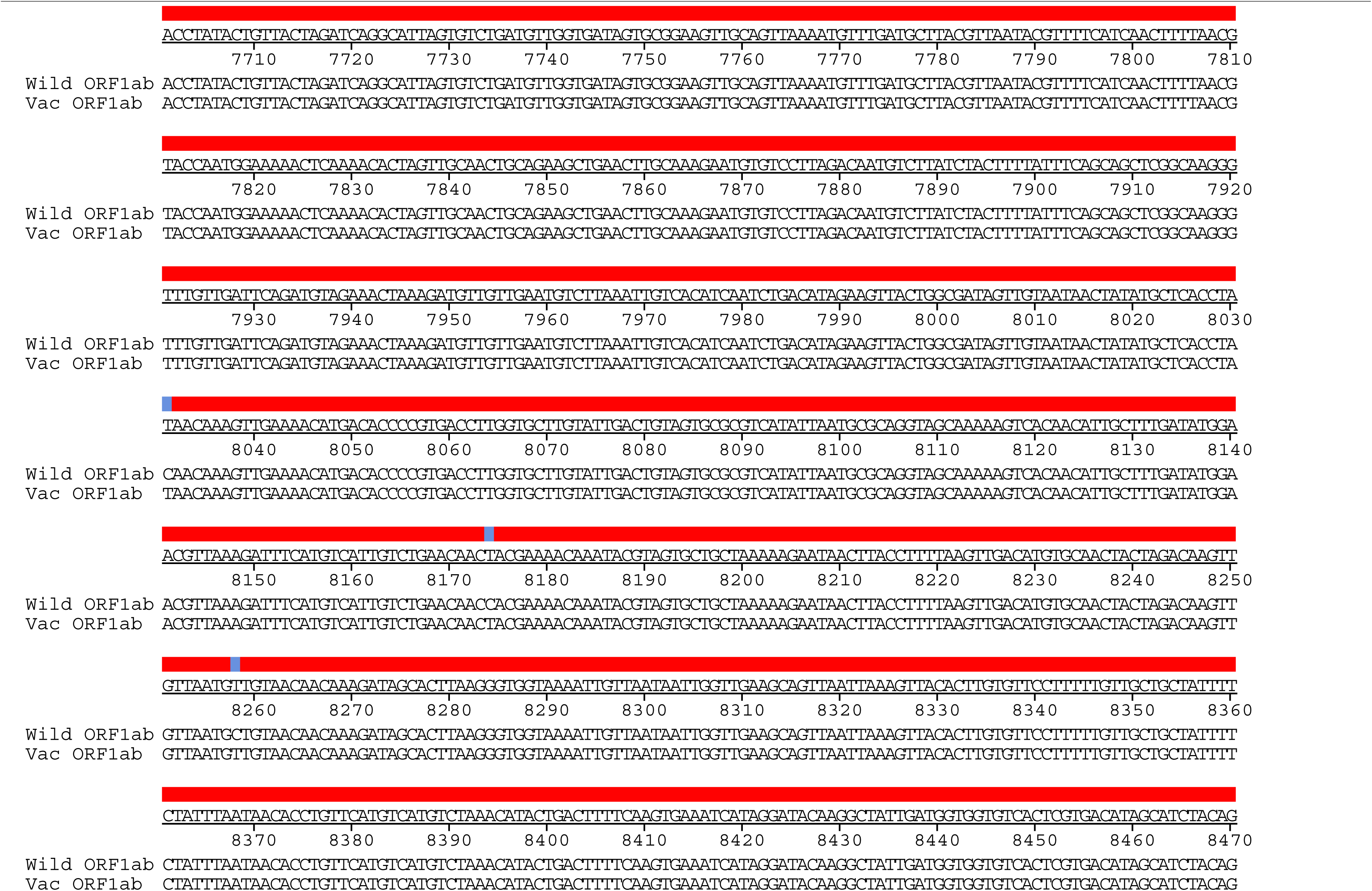
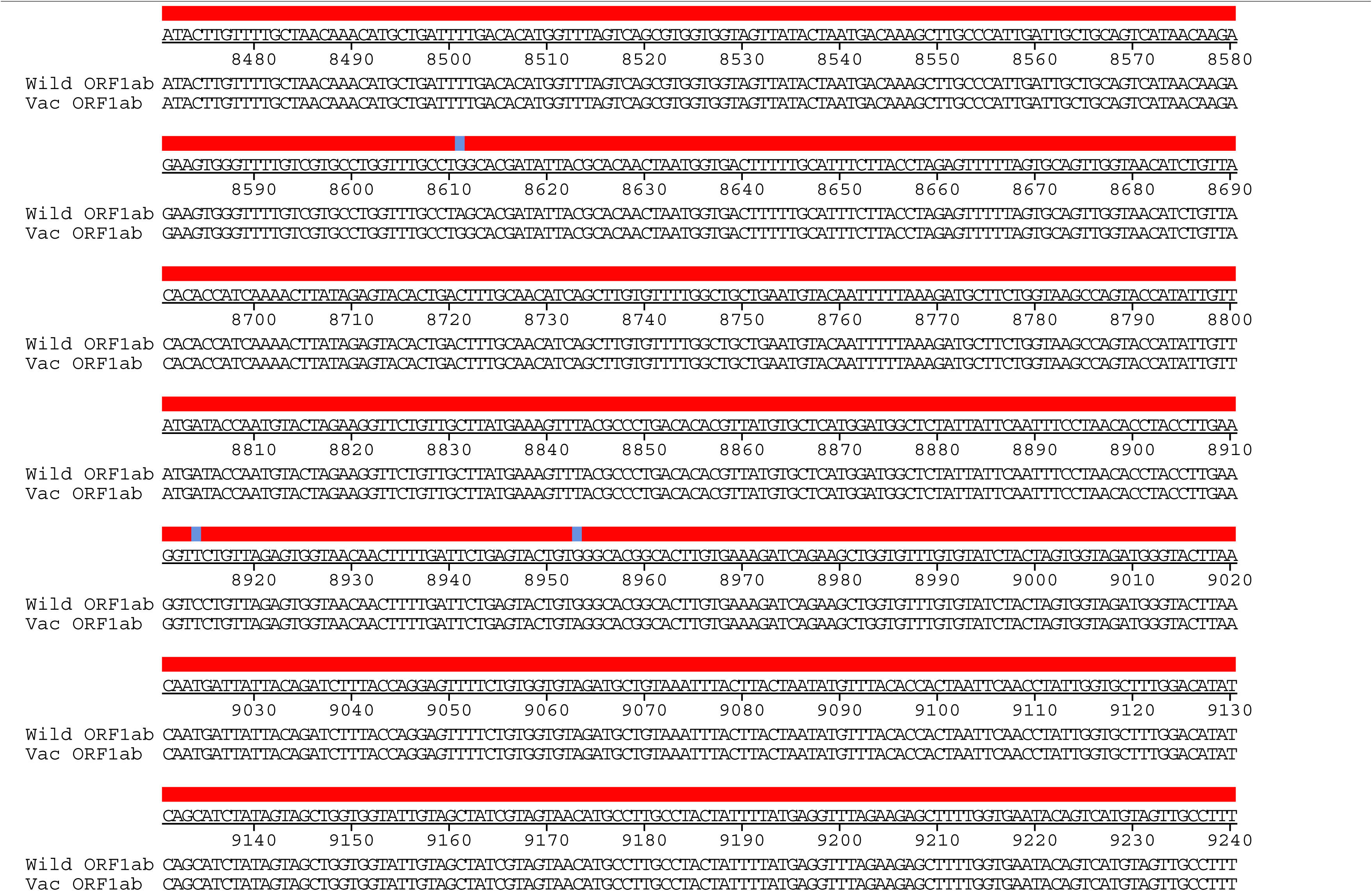
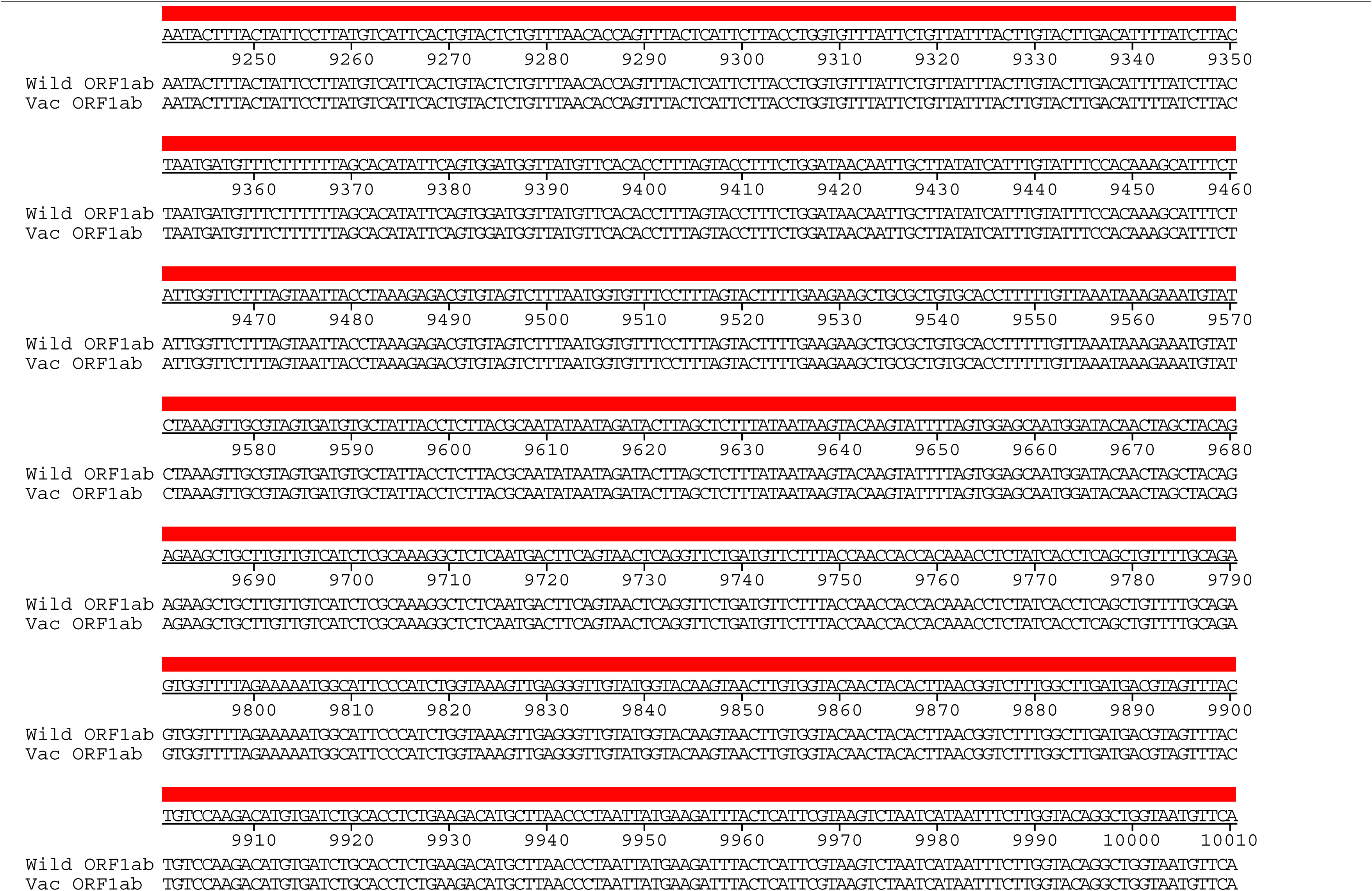
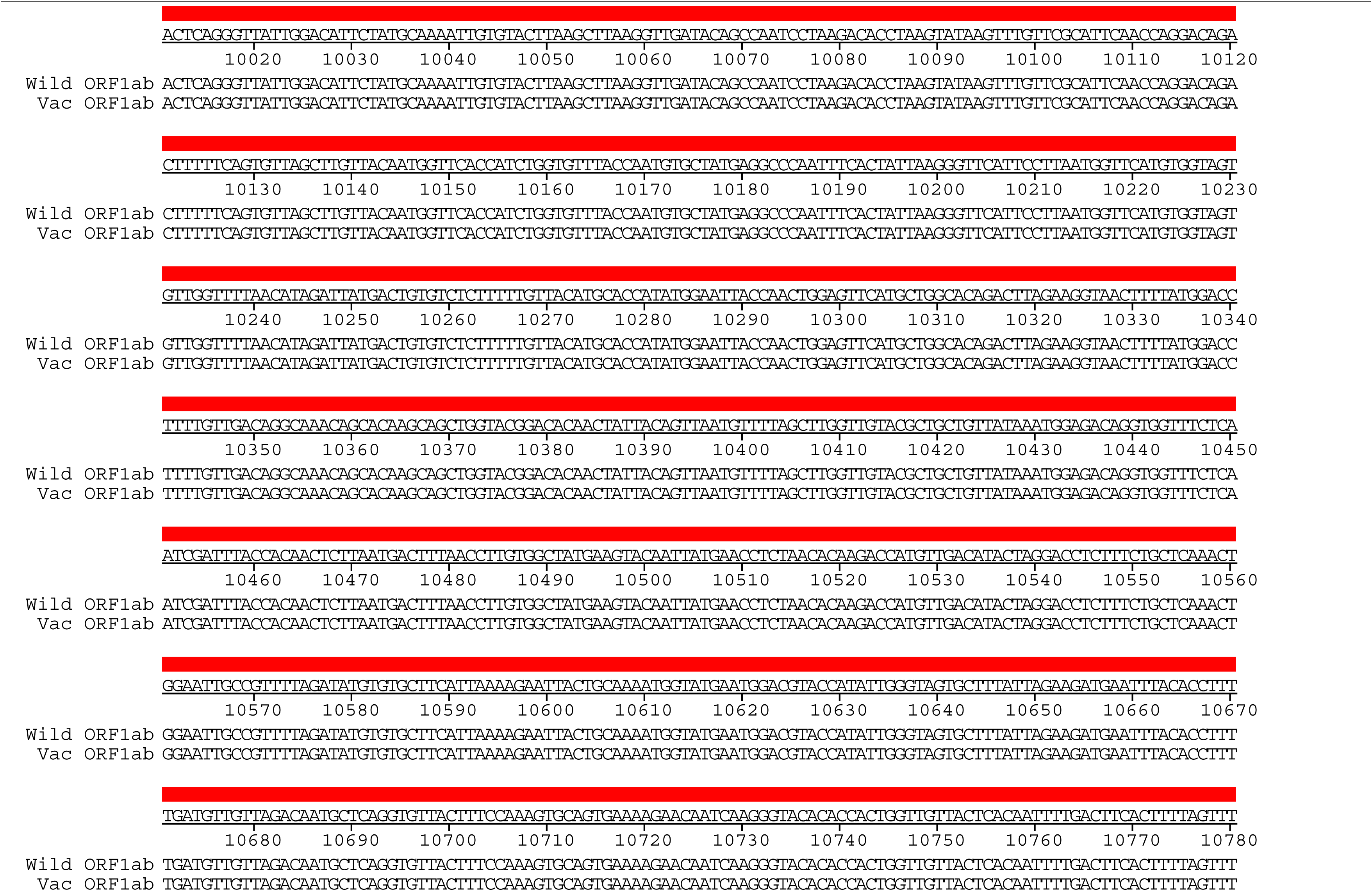
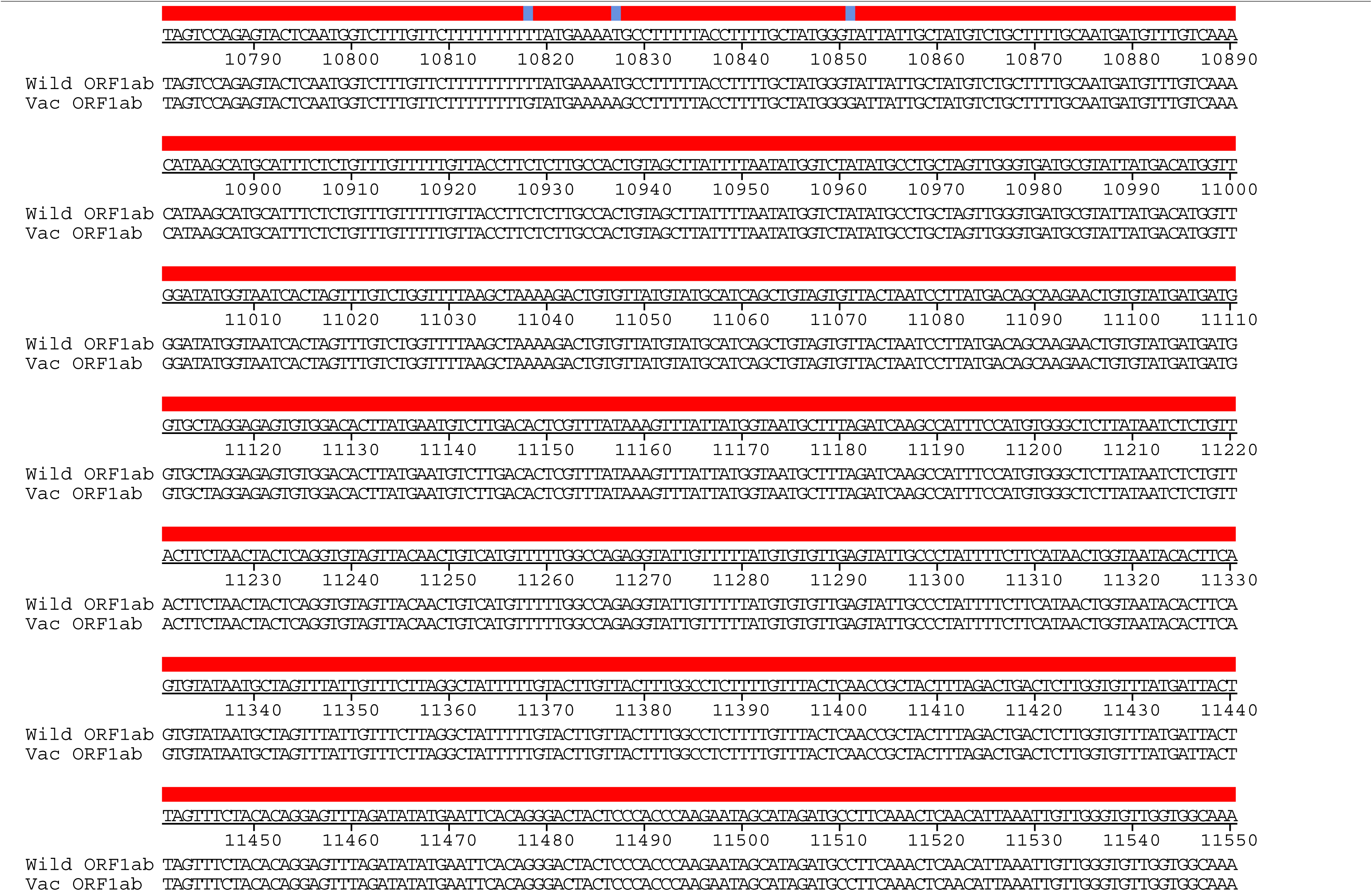
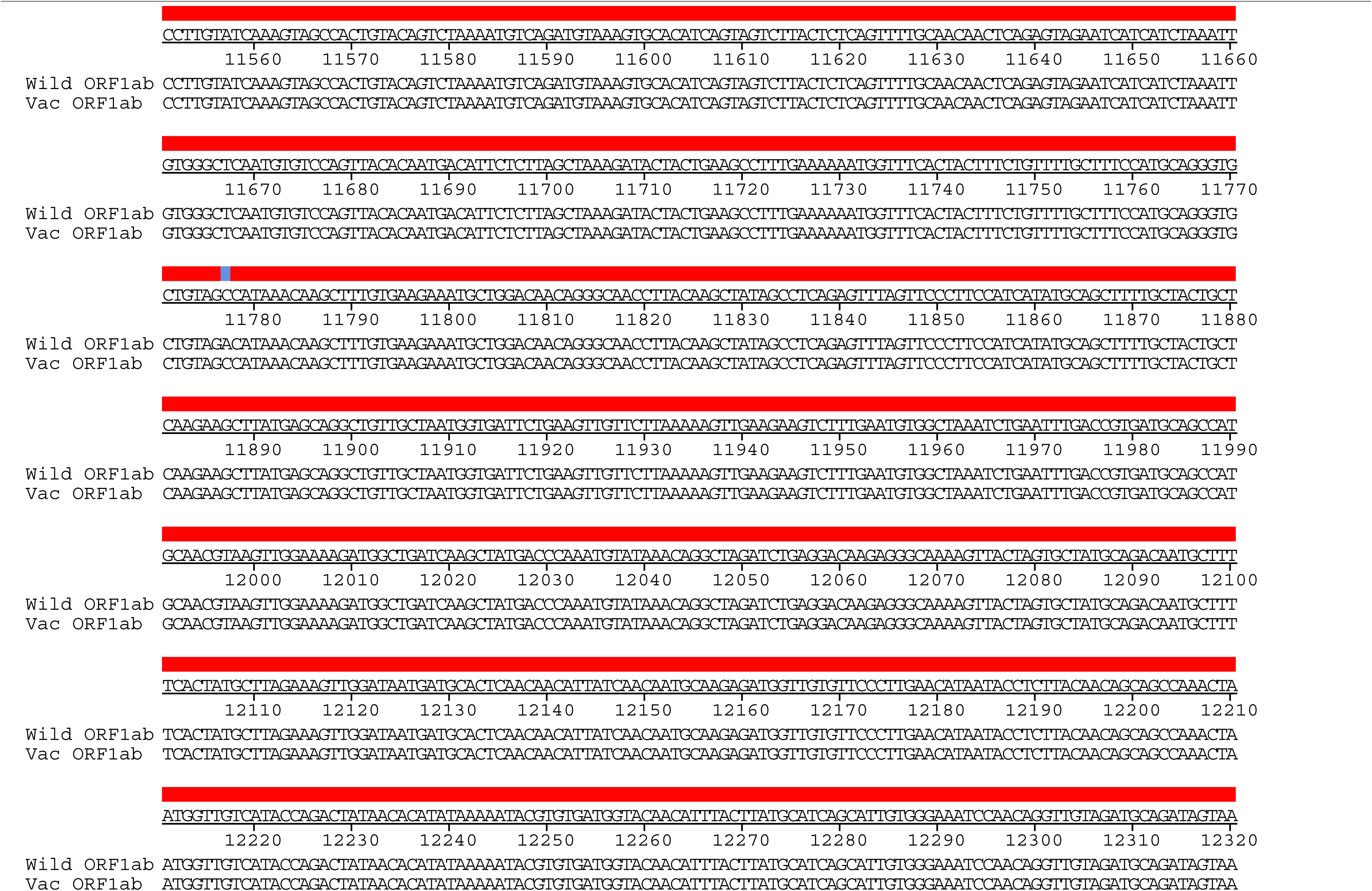
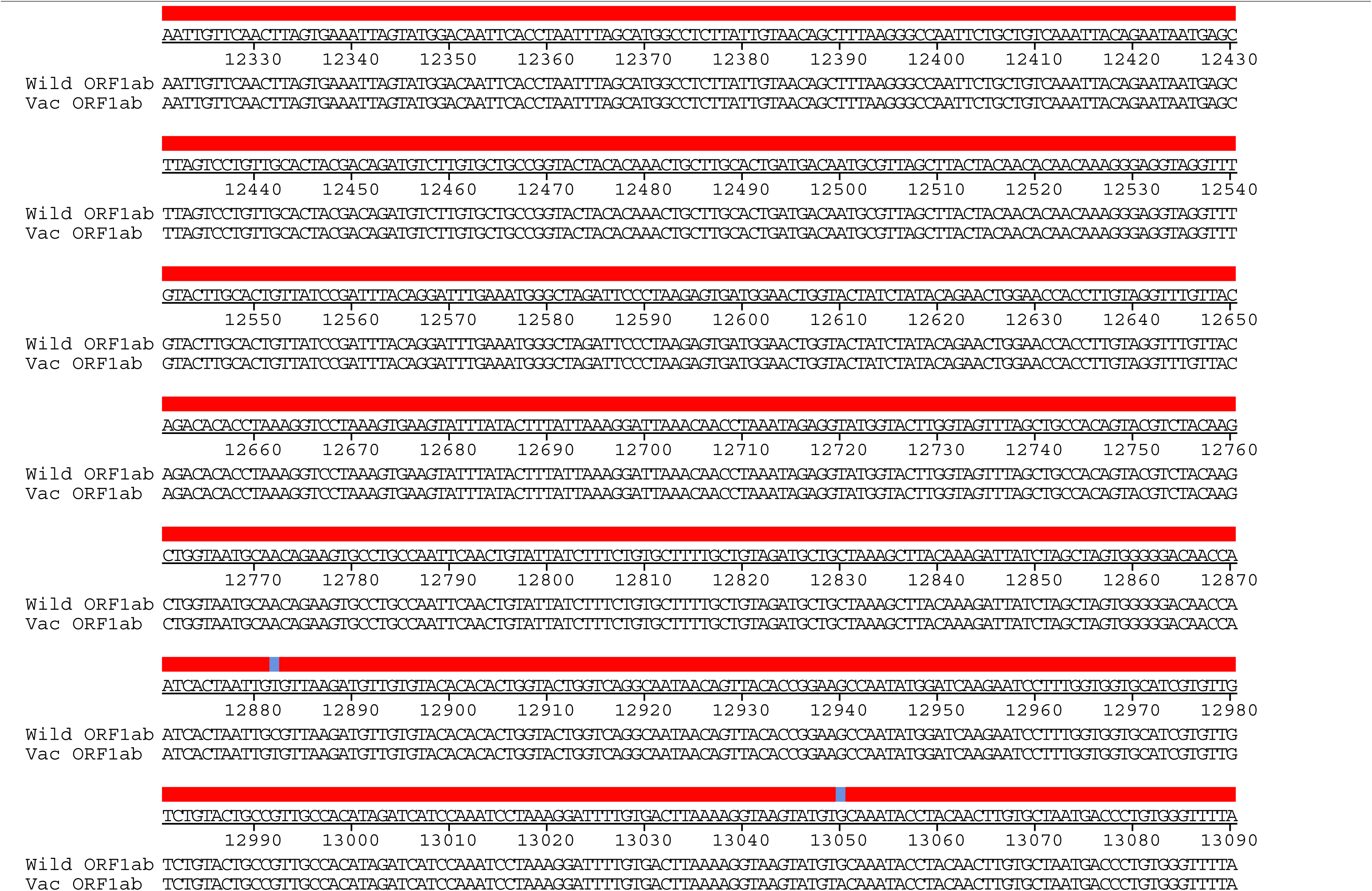
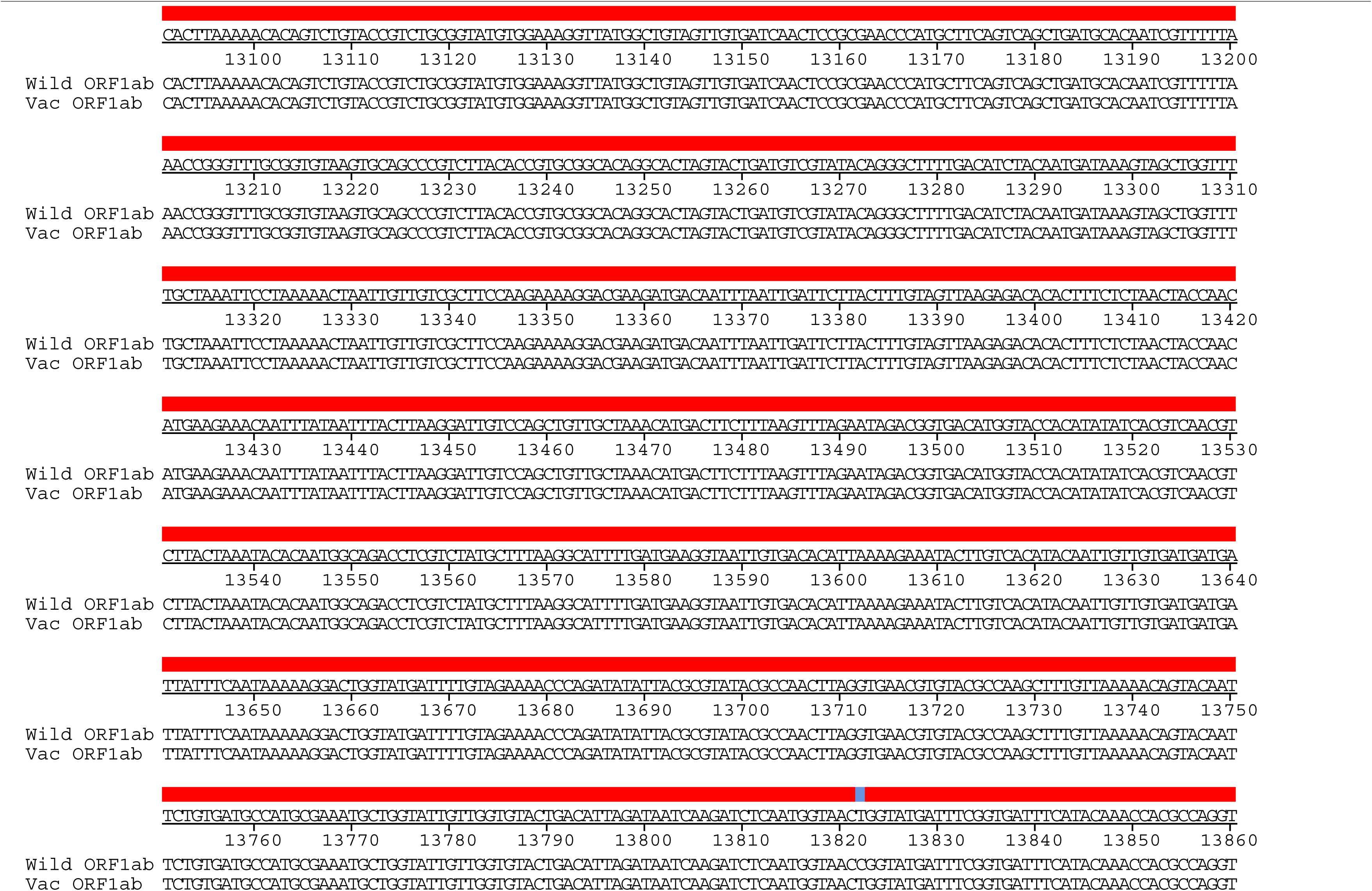
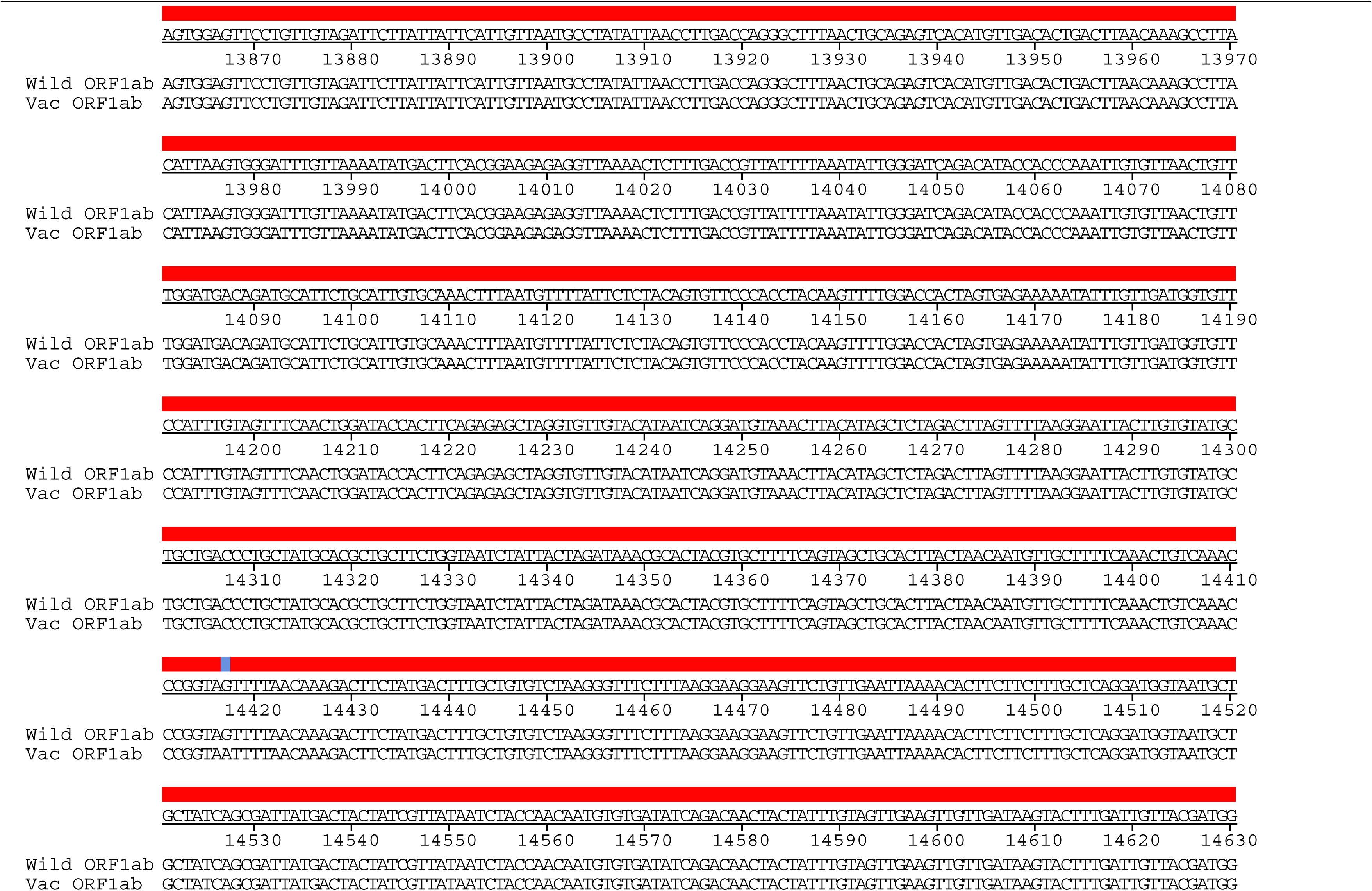
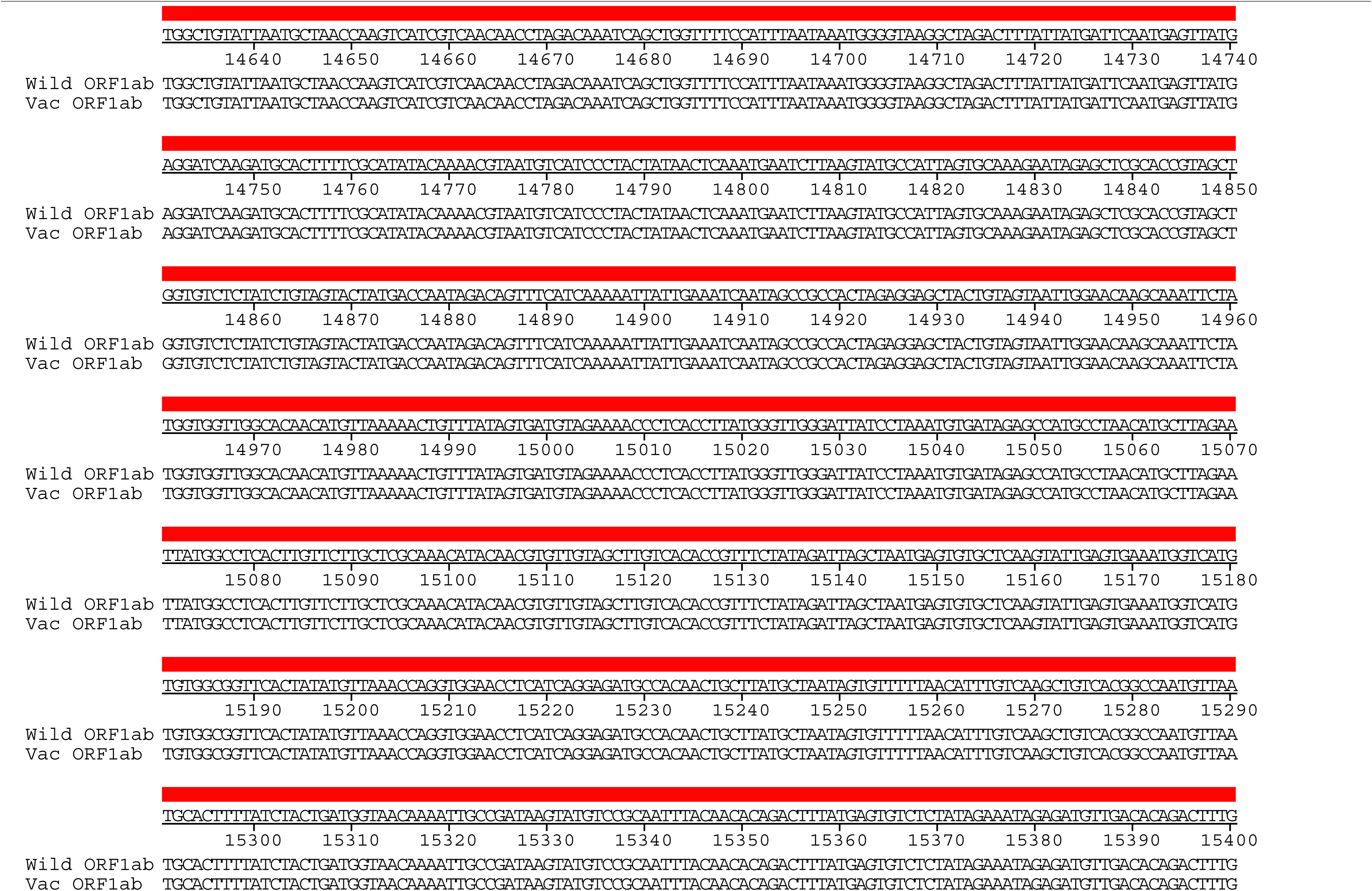
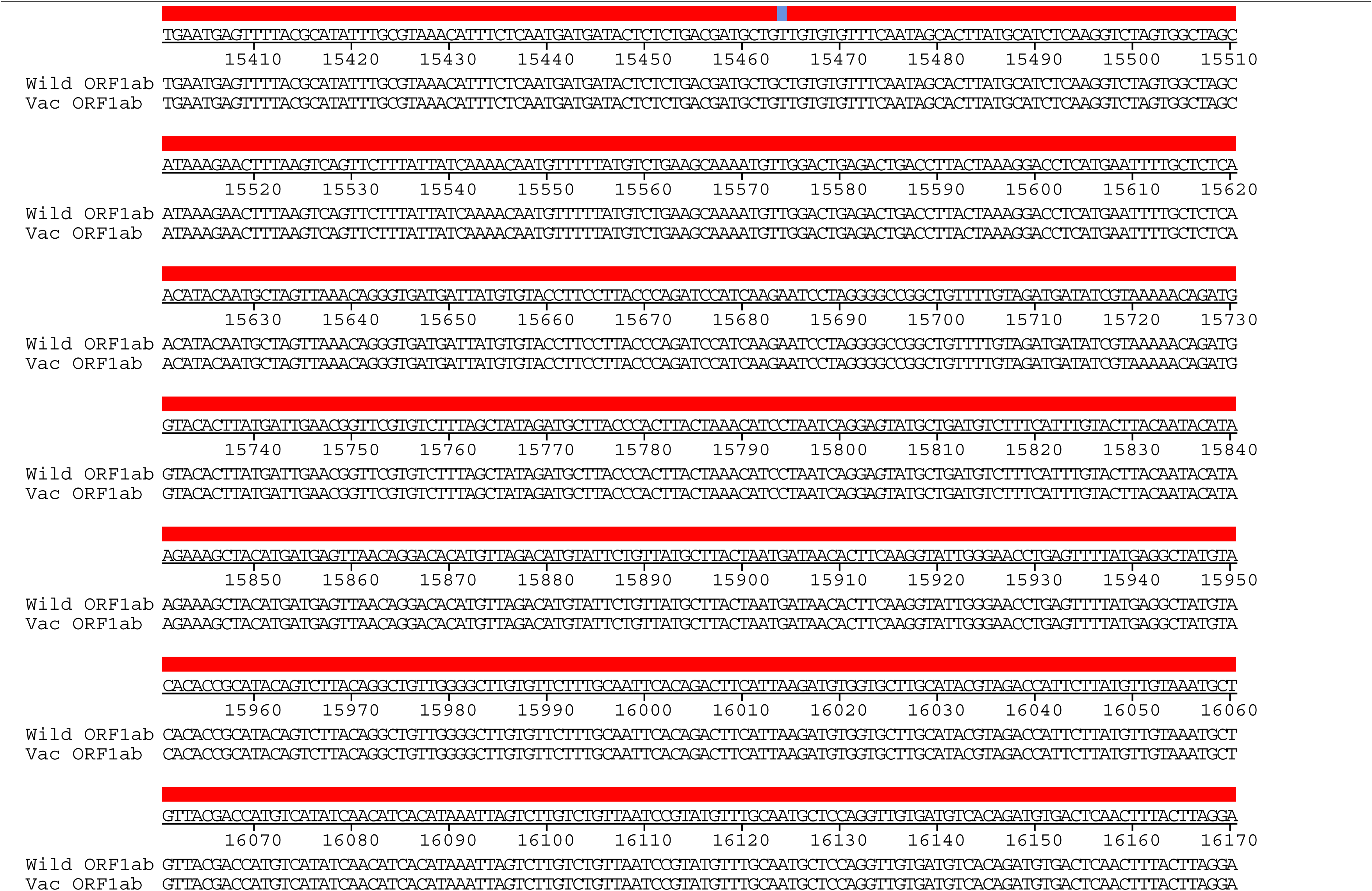
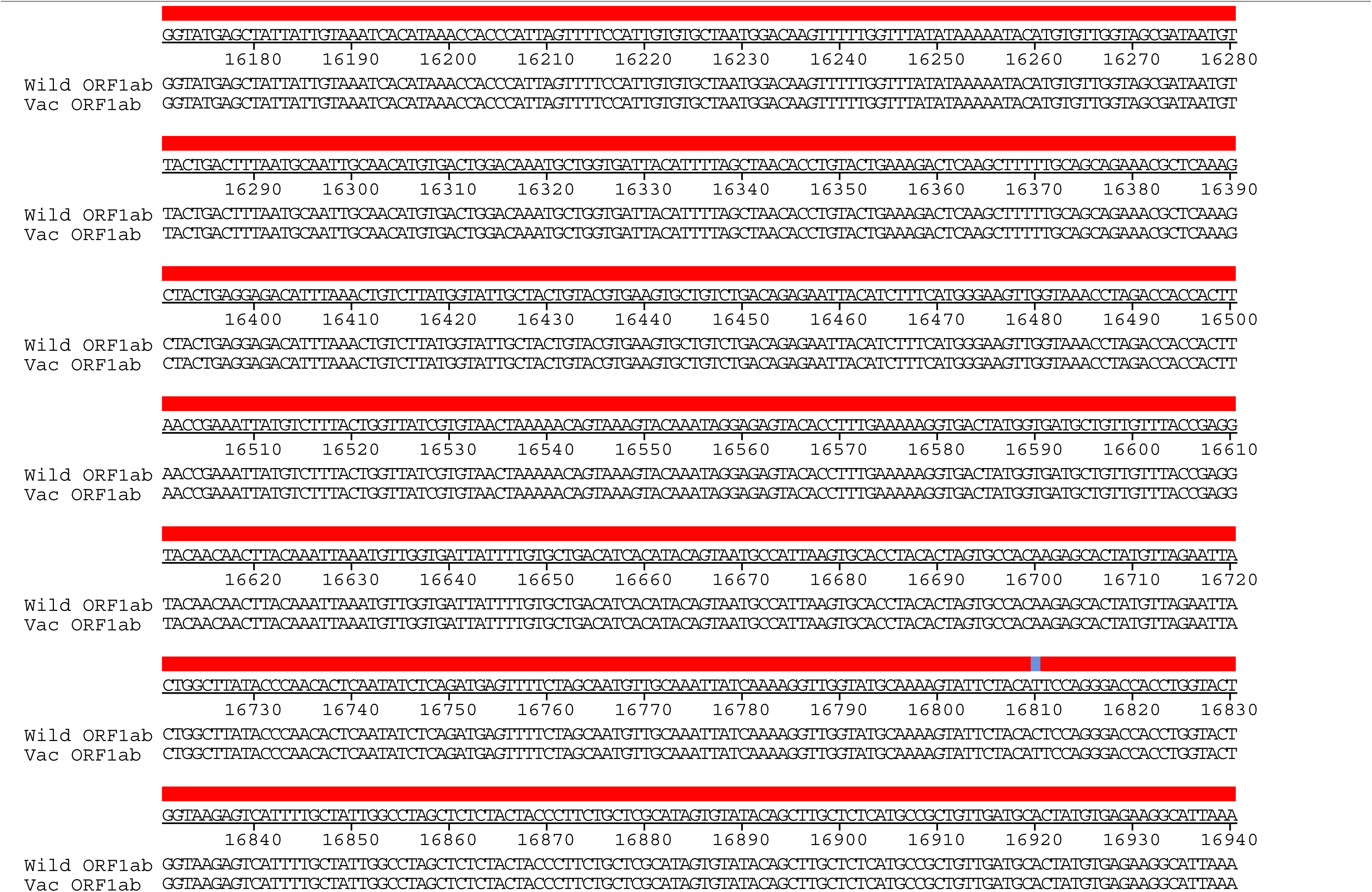
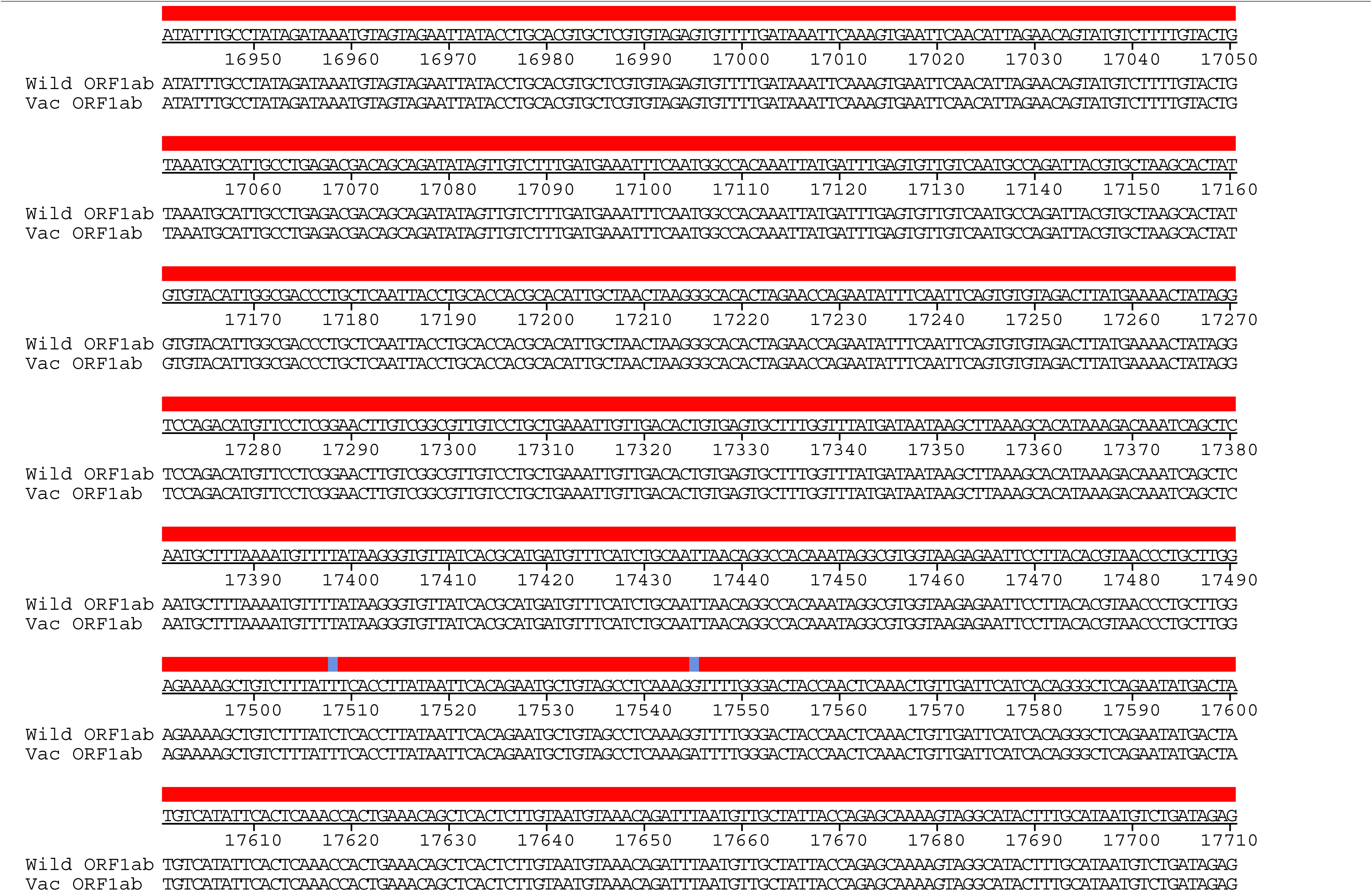
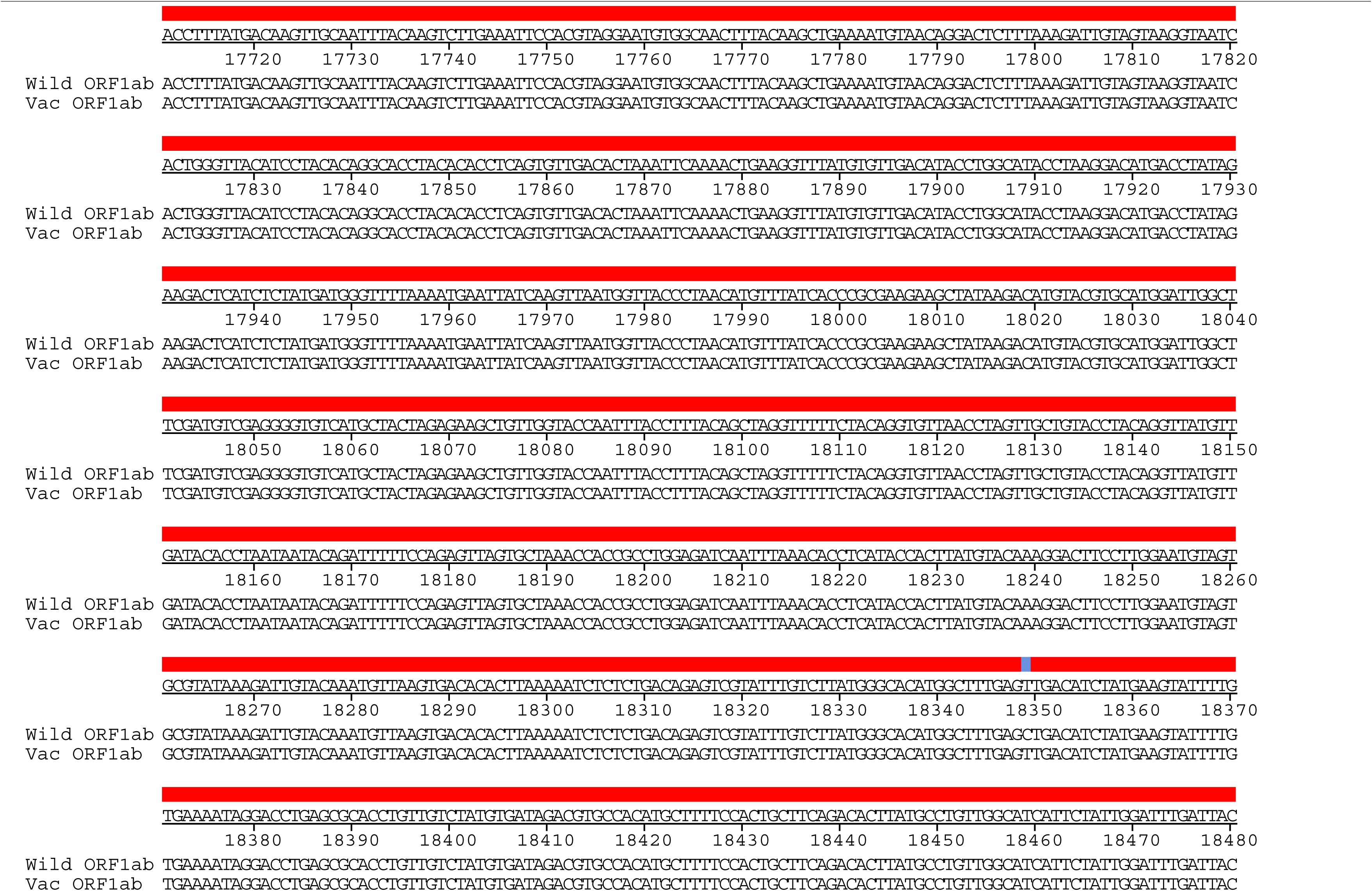
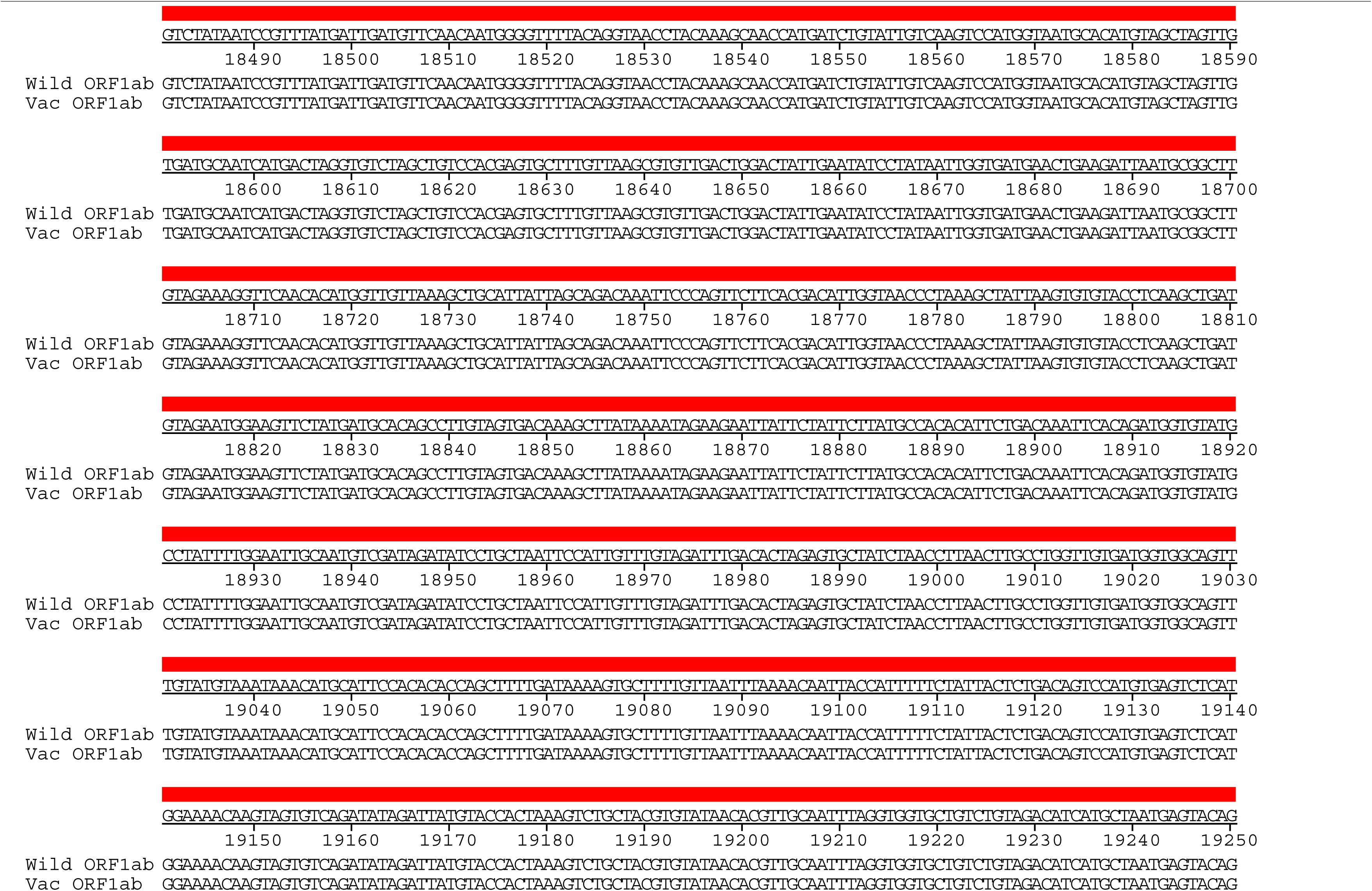
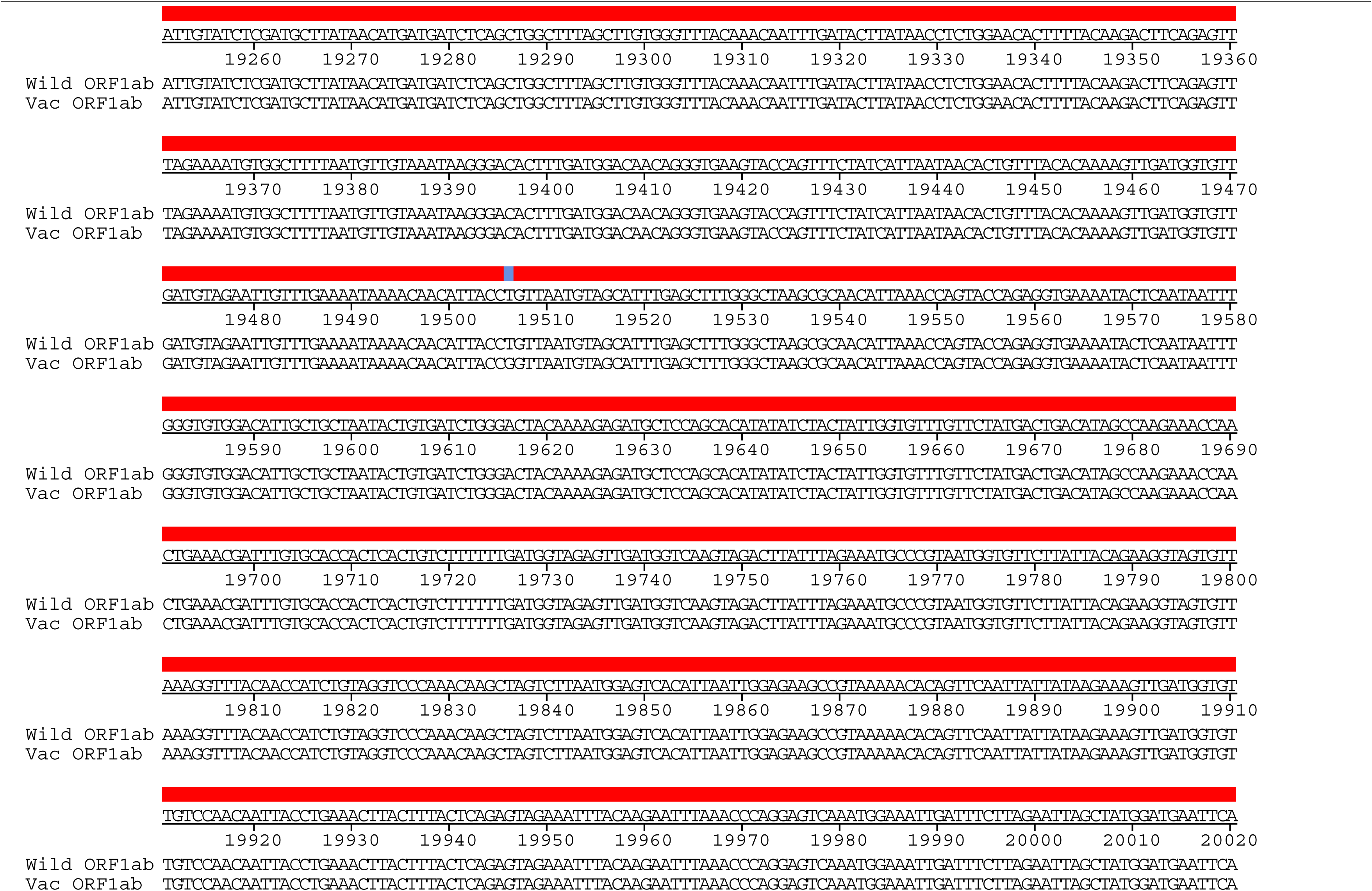
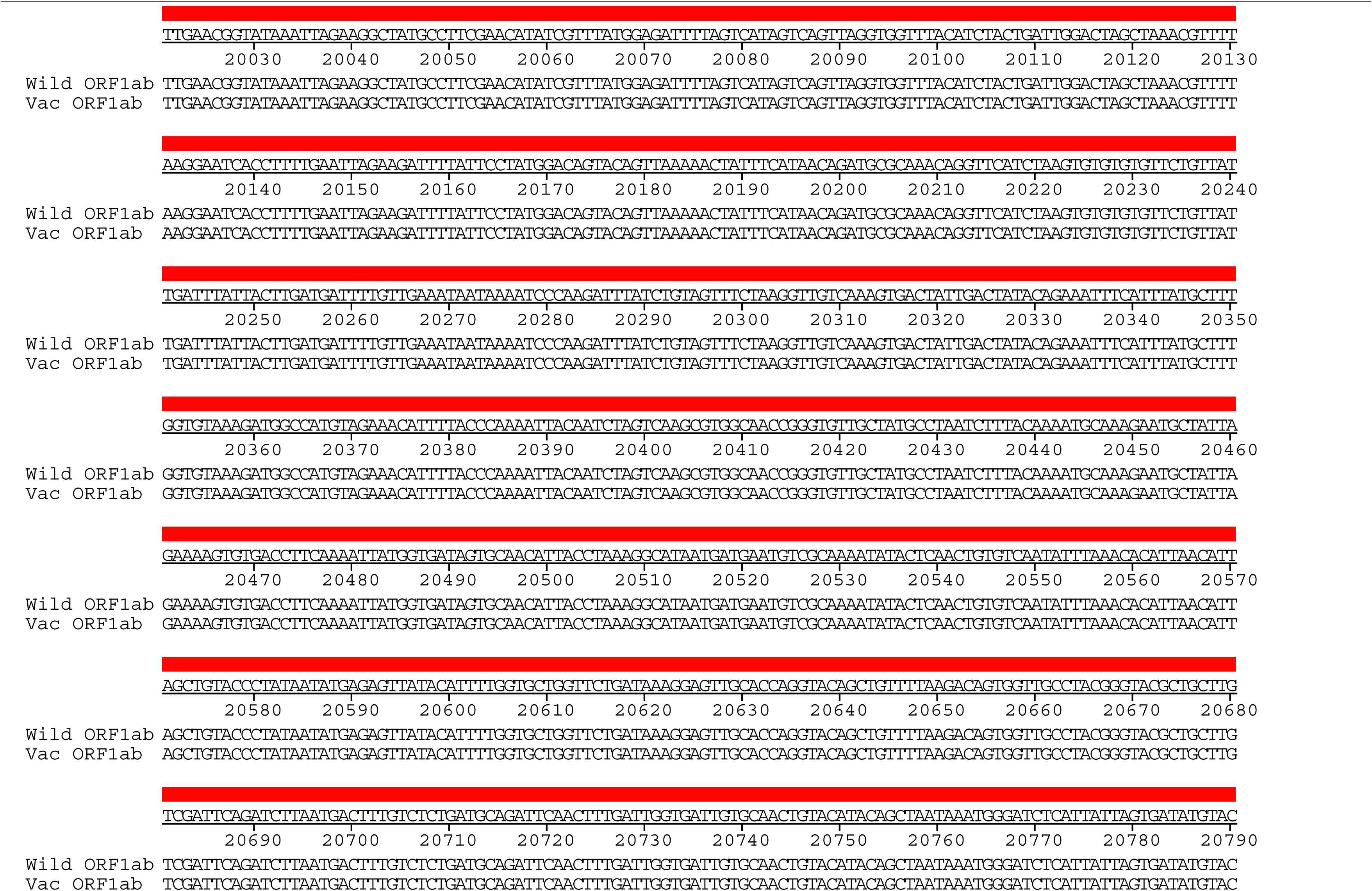
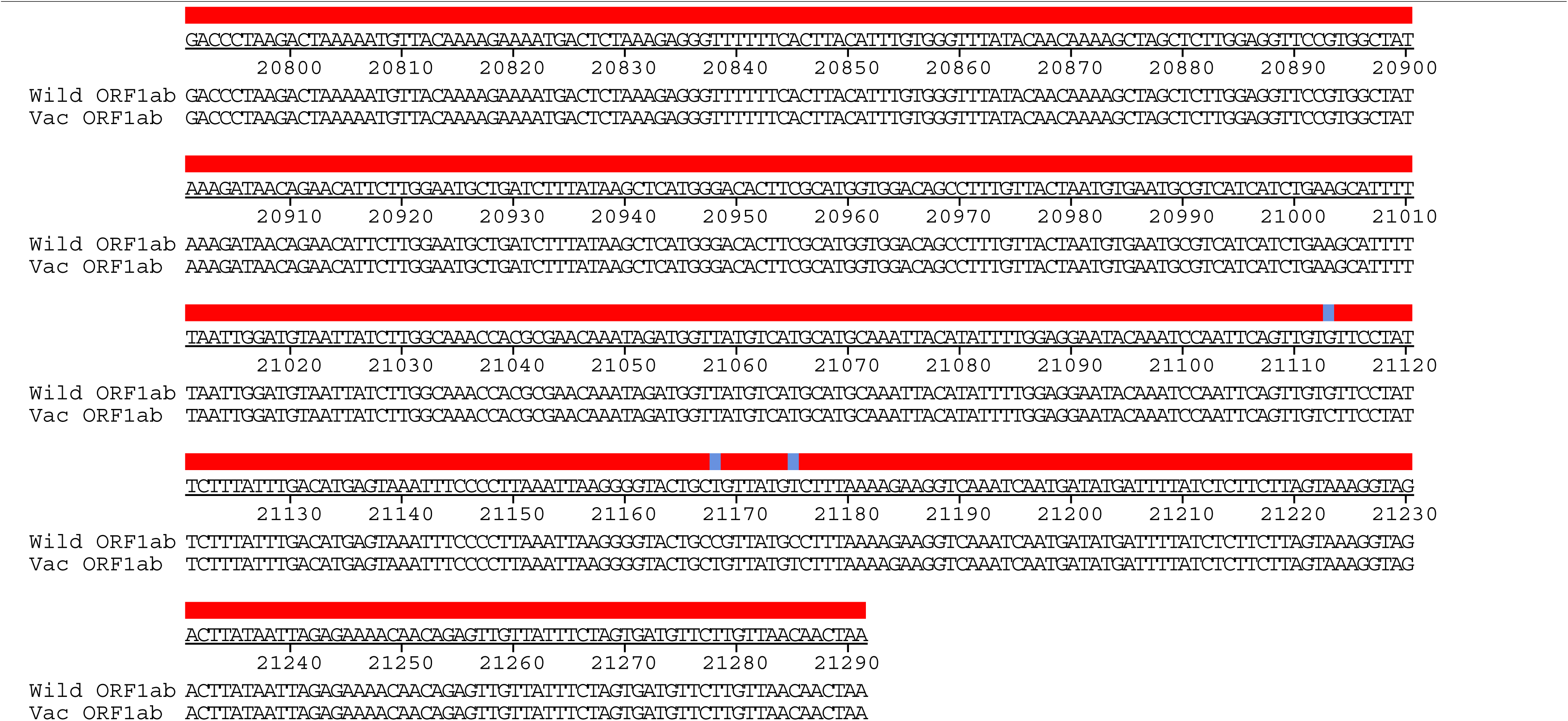
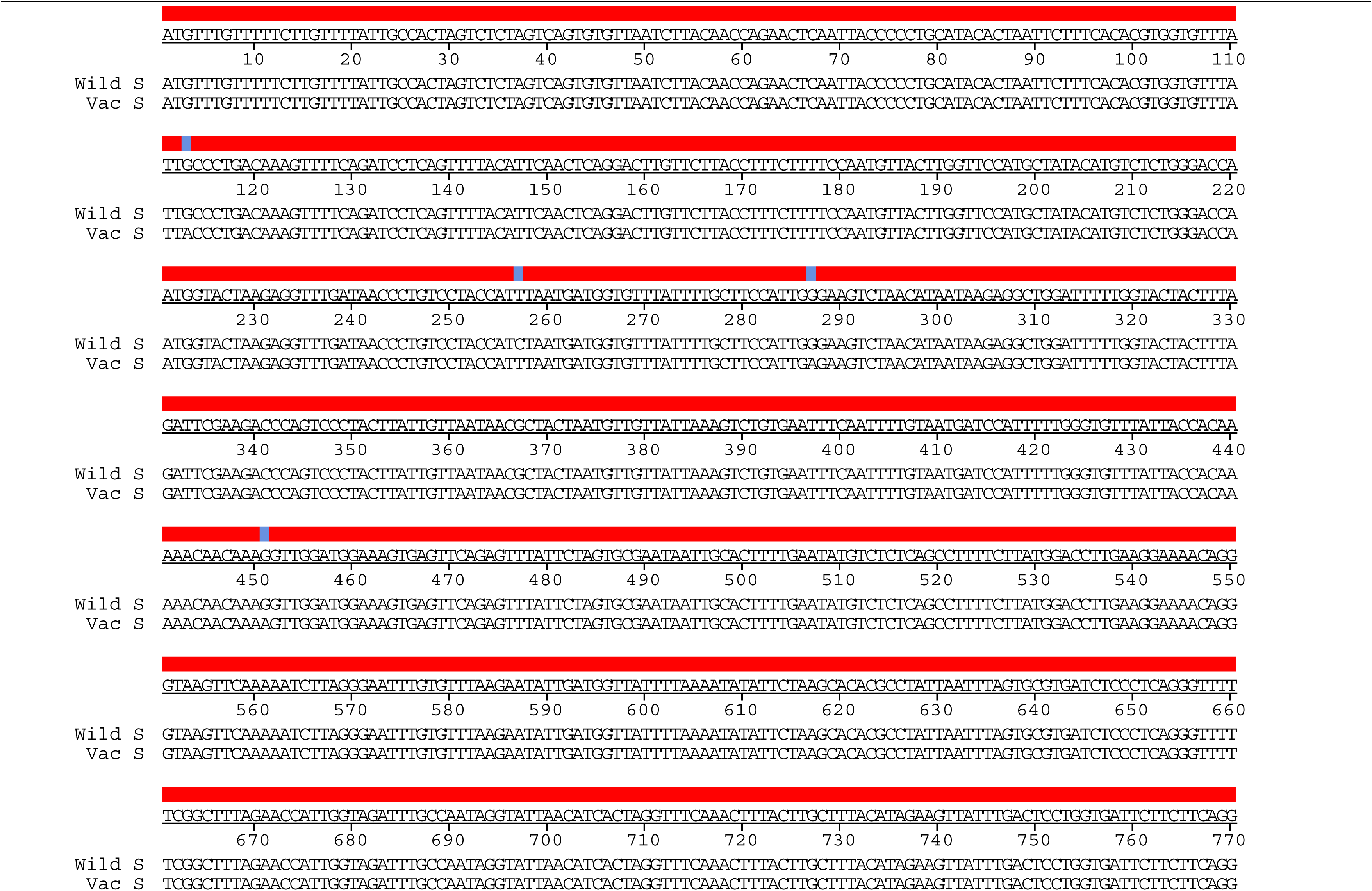
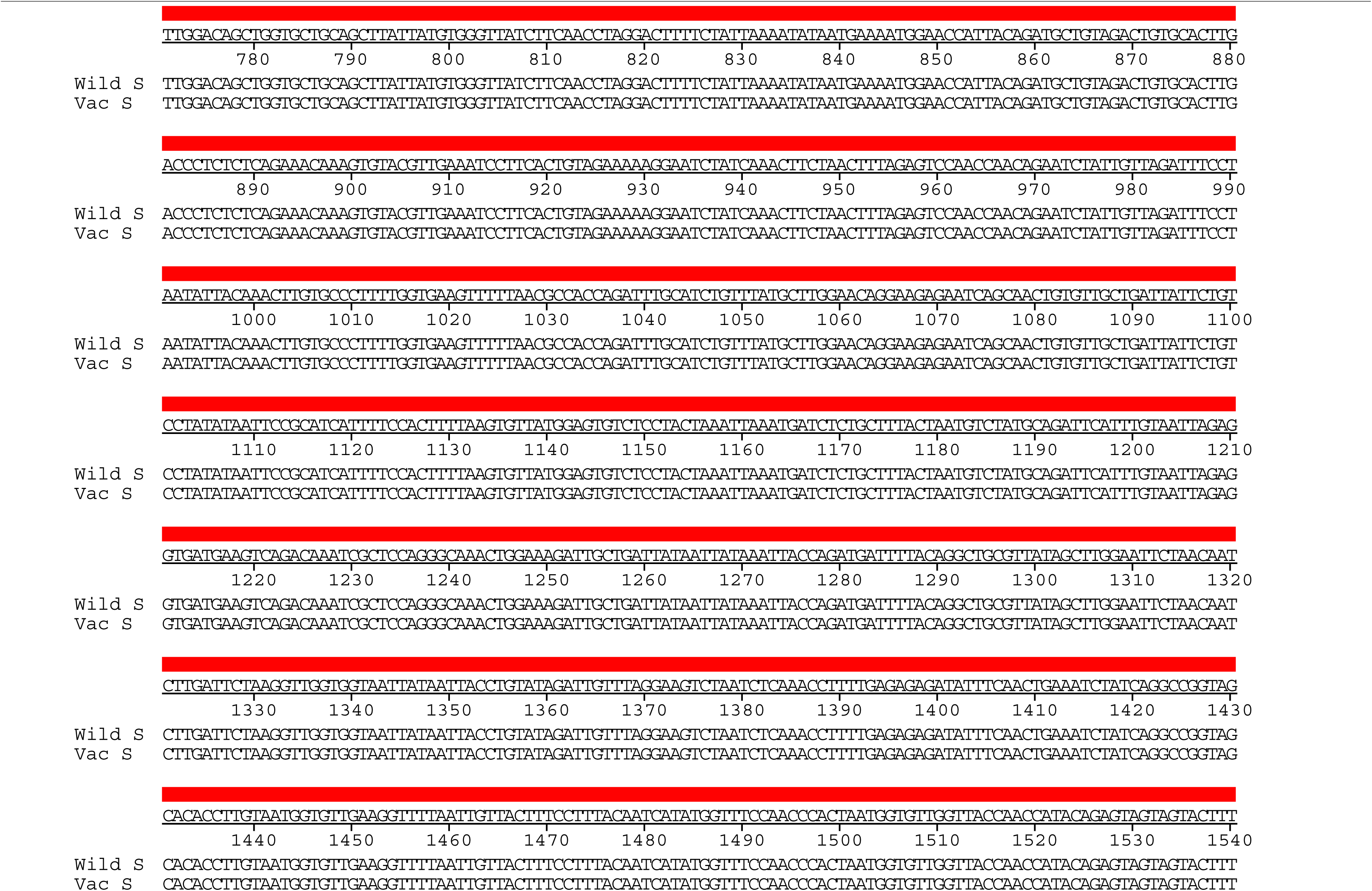
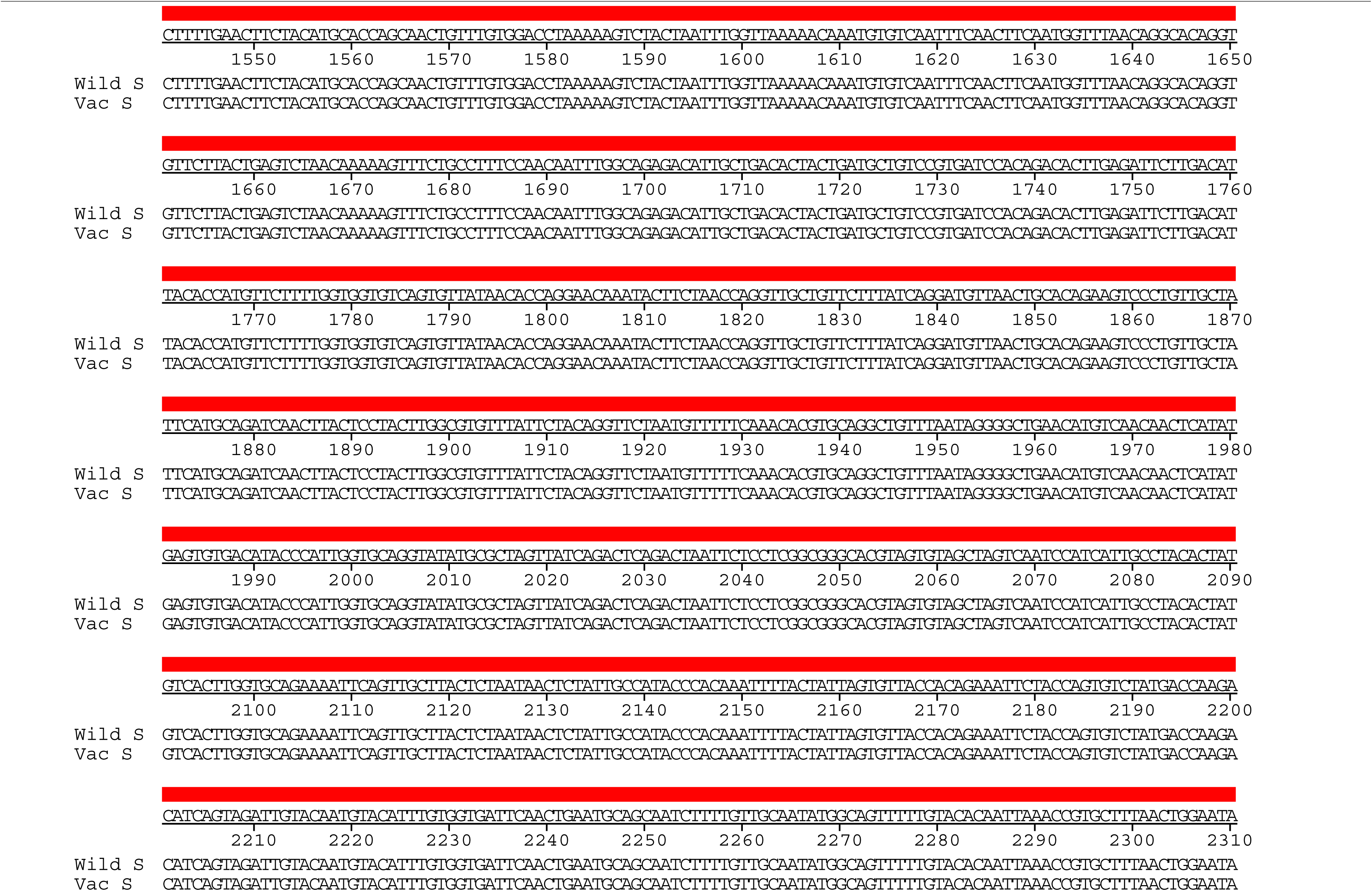
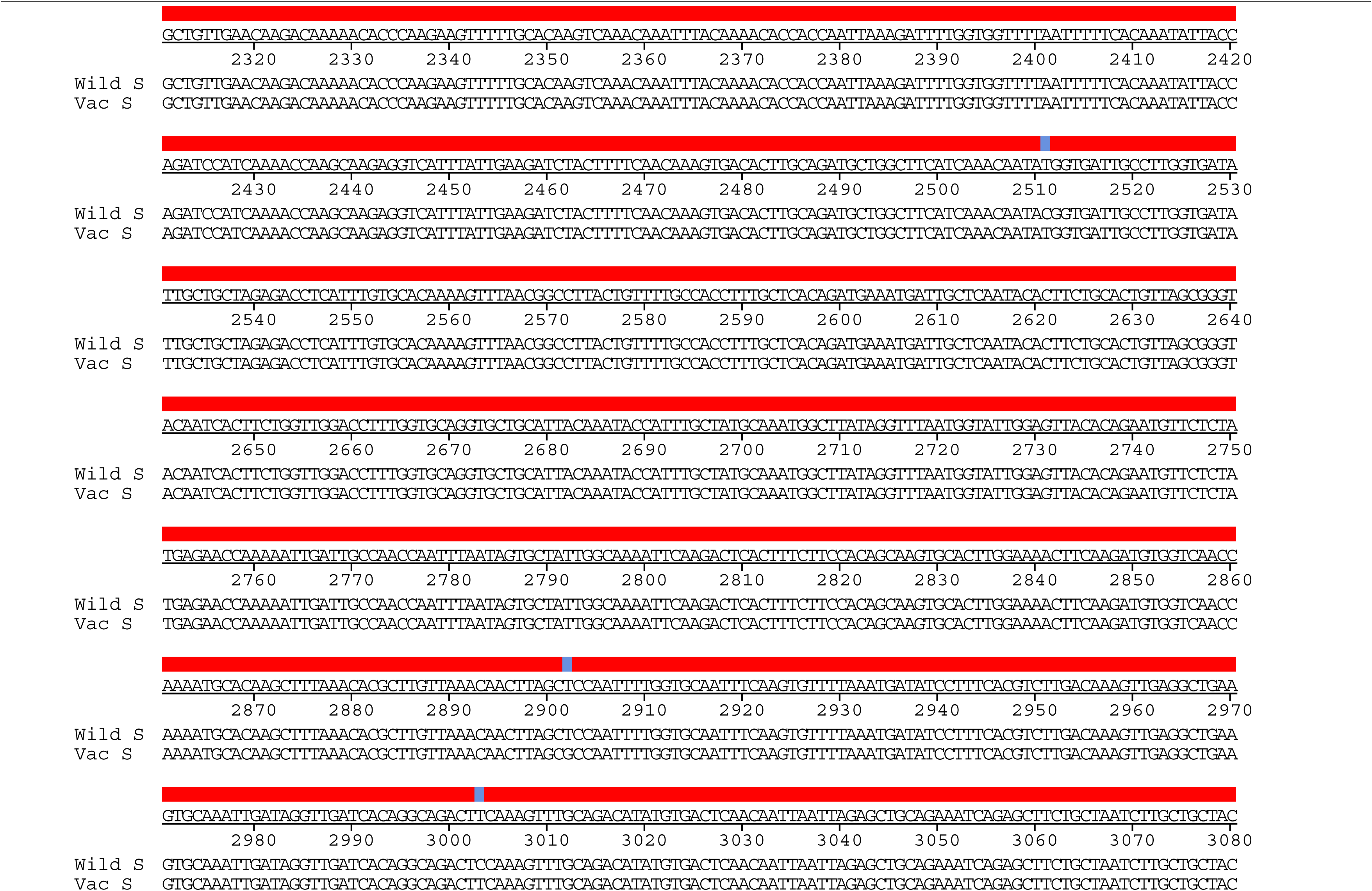
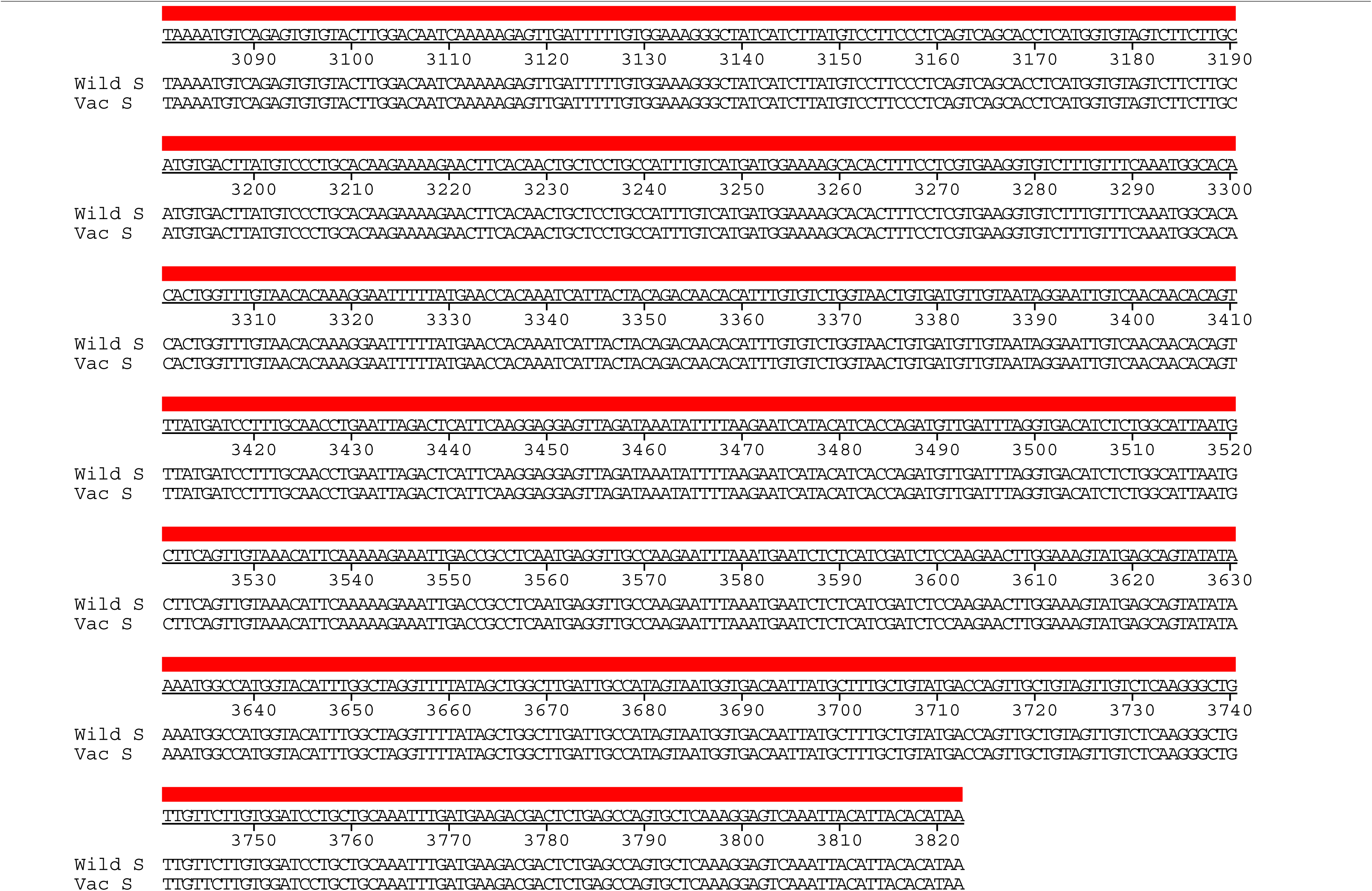
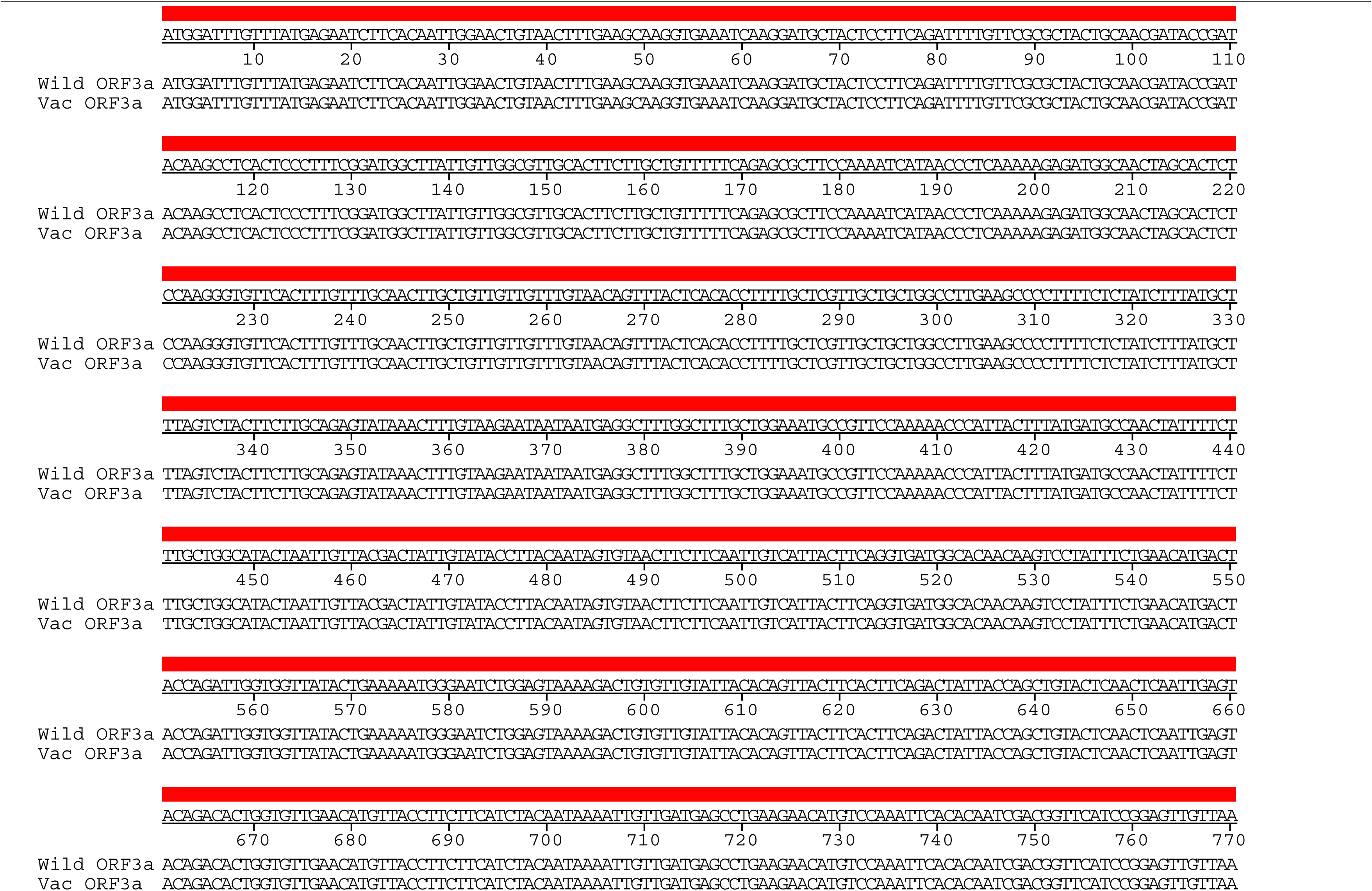
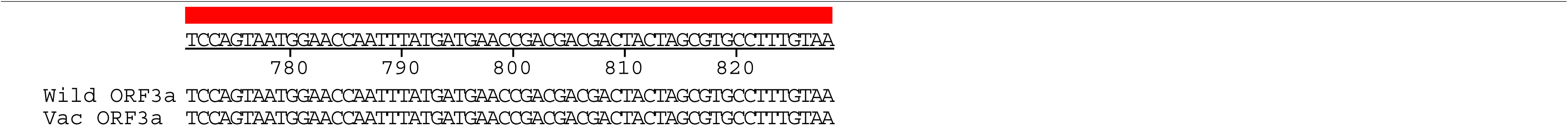
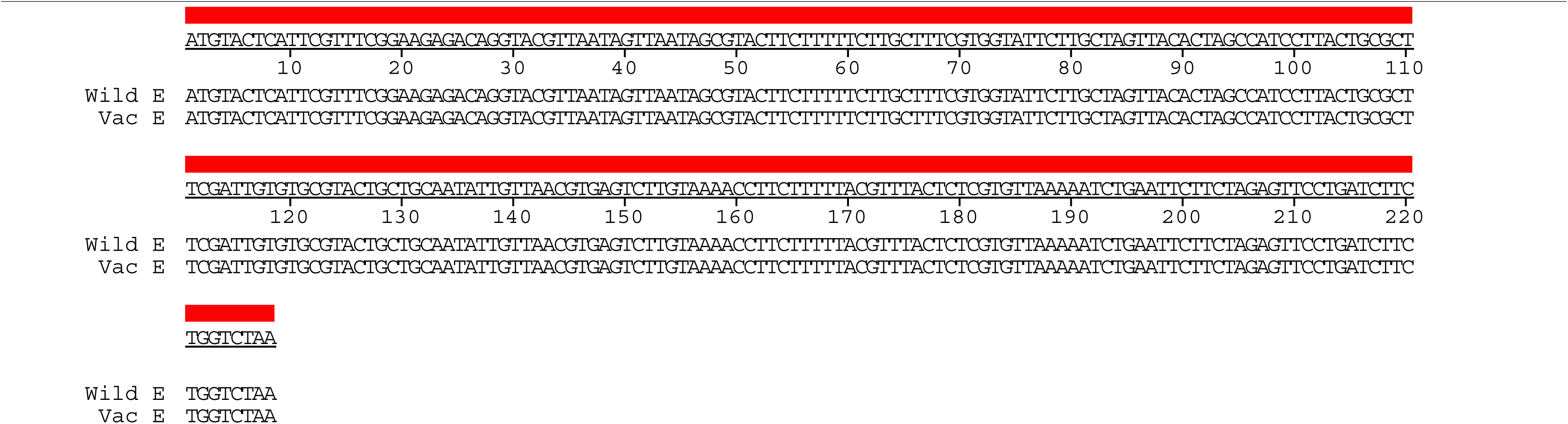
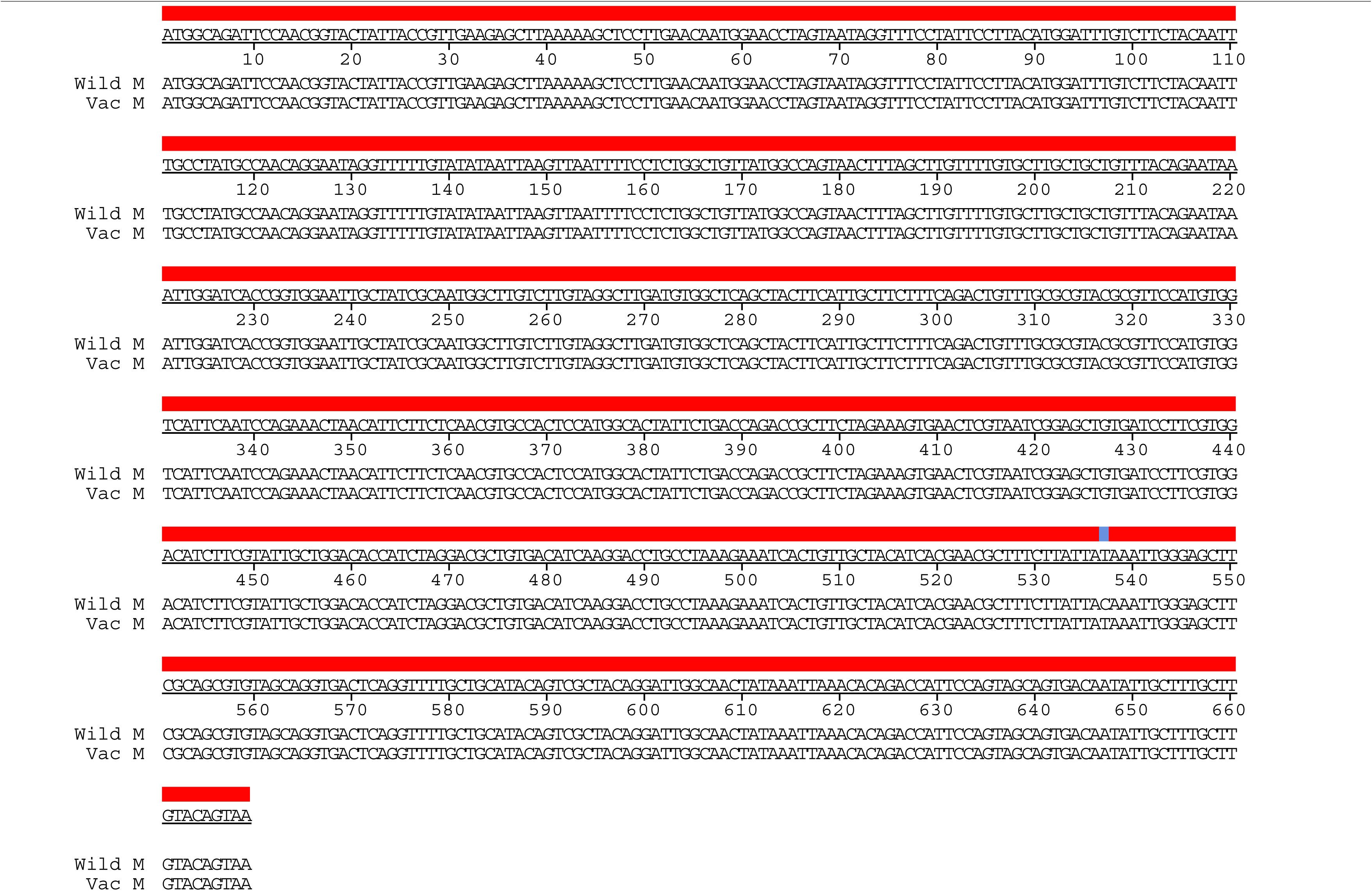
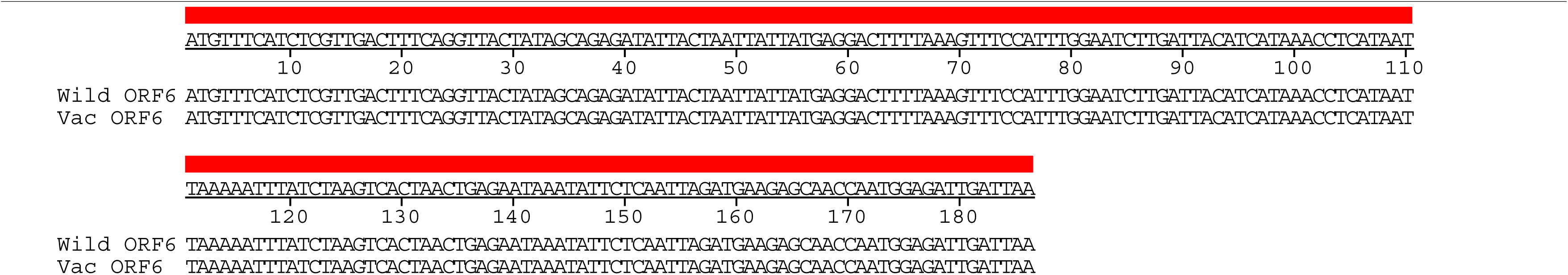
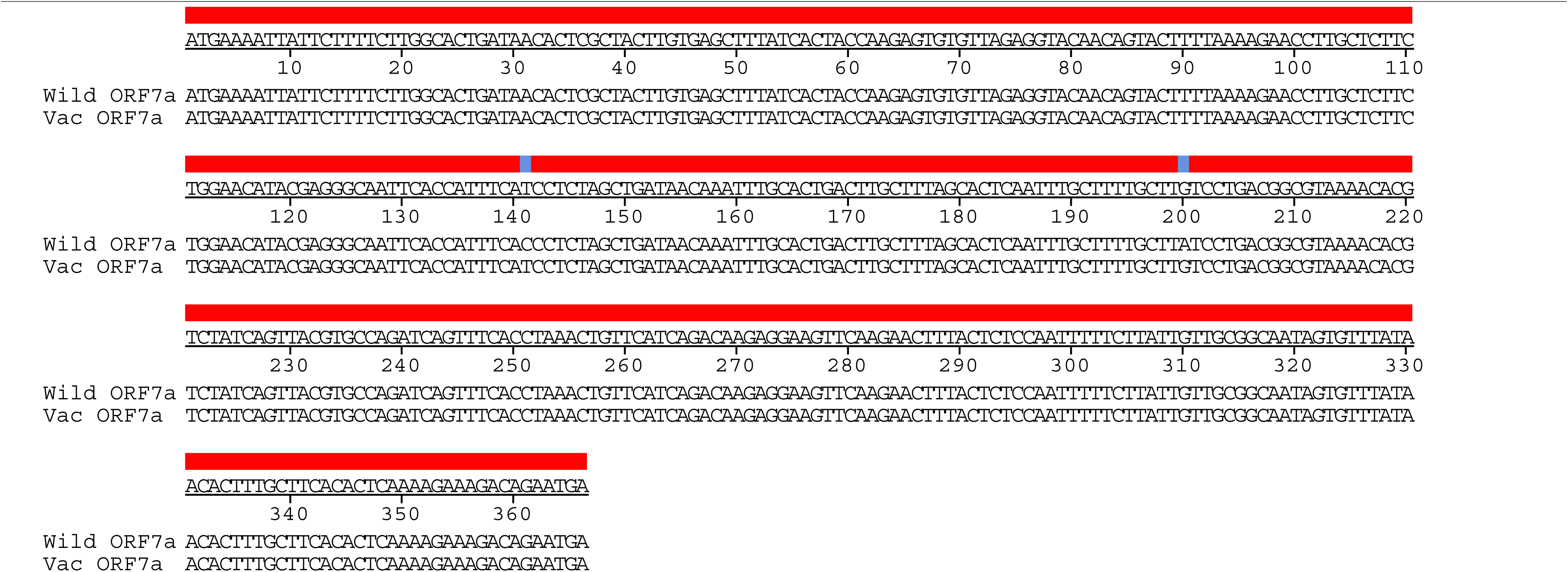
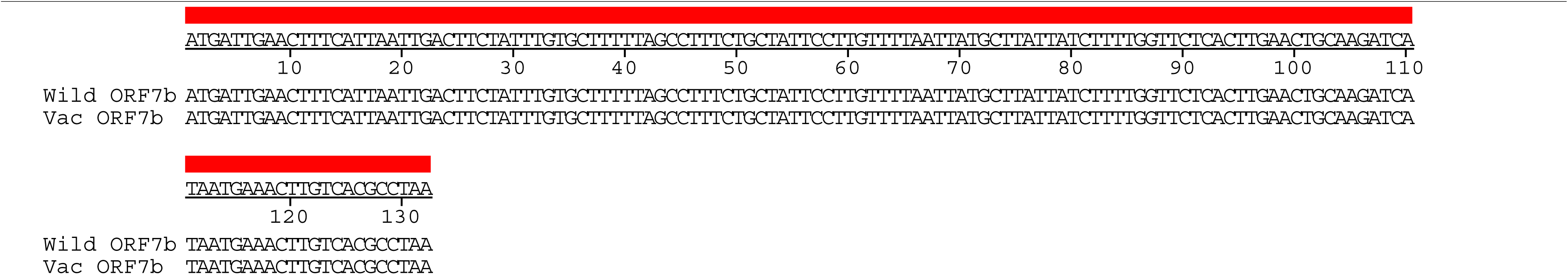
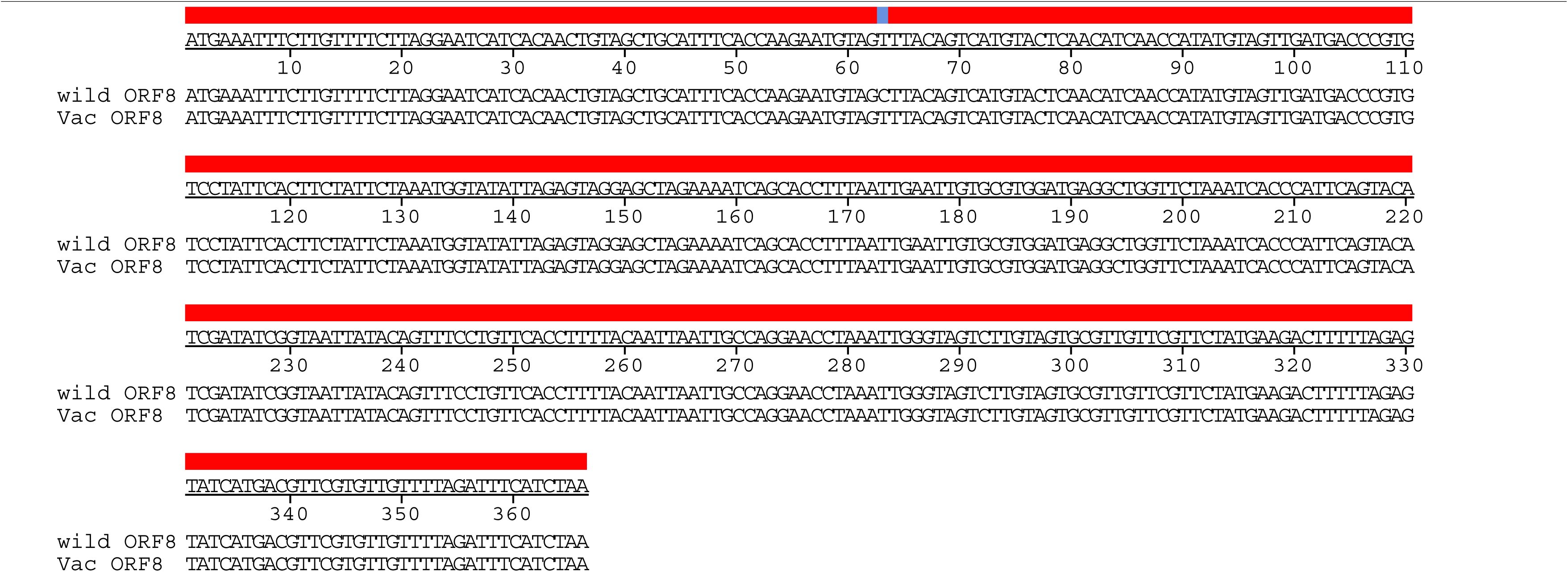
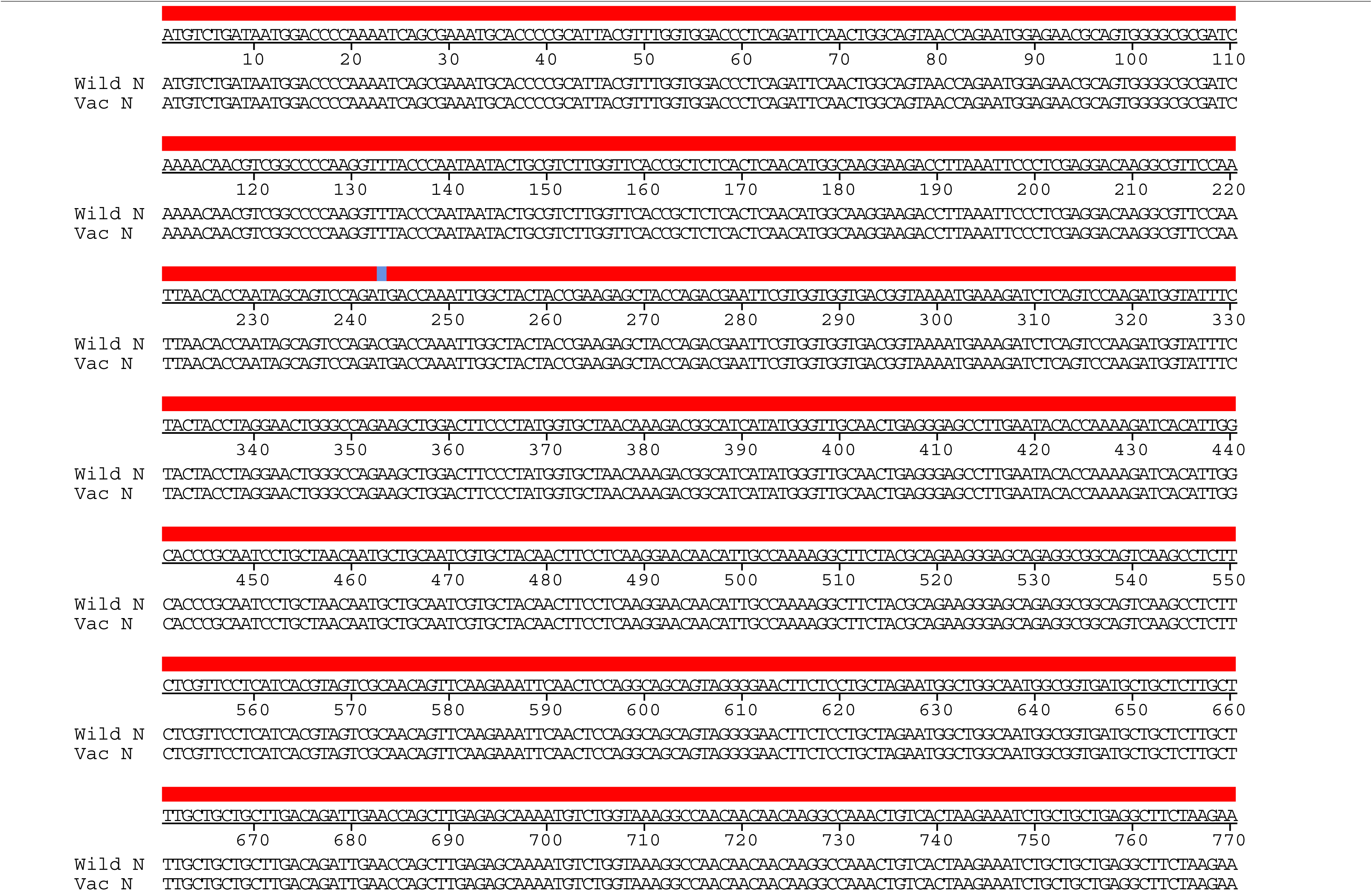
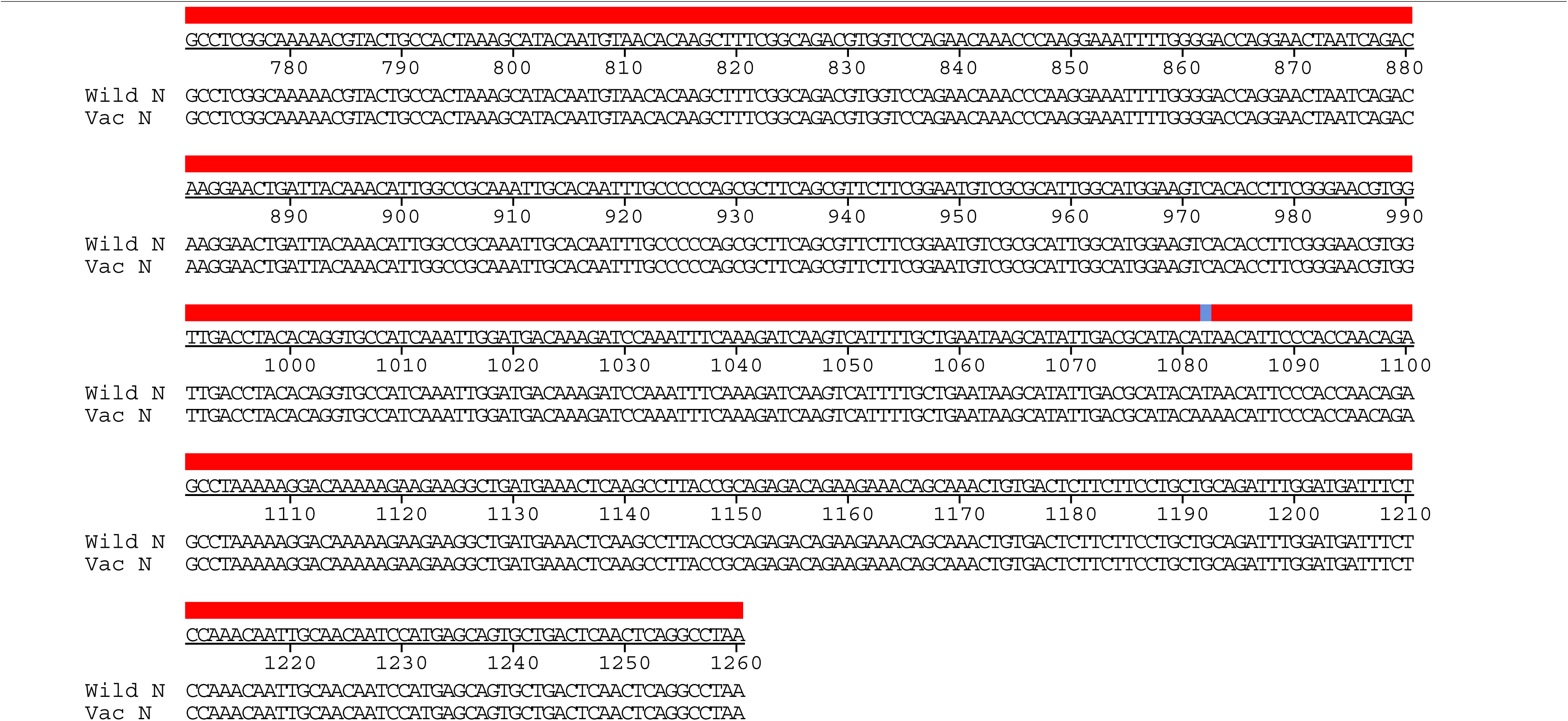
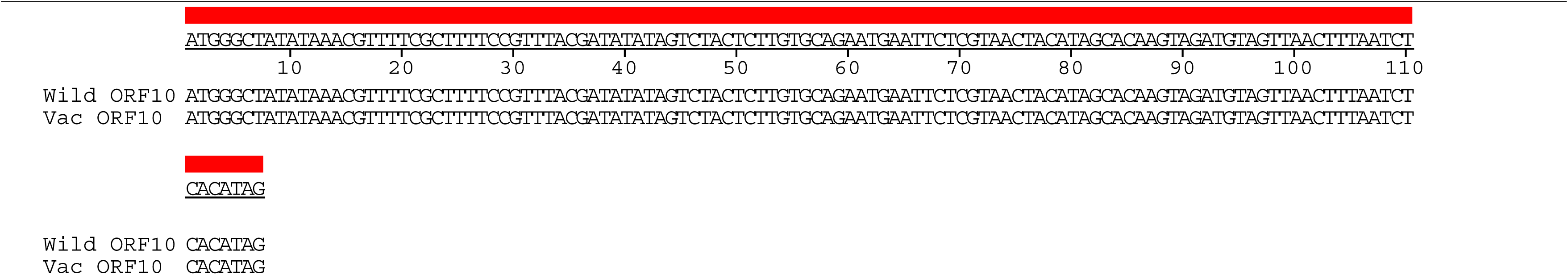
Nucleotide alignment of cold-adapted live attenuated SARS-CoV-2 vaccine strain. Full nucleotide sequences of cold-adapted live attenuated SARS-CoV-2 vaccine strain (CoV-2-CNUHV 03-CA22℃) were aligned with that of wild-type SARS-CoV-2 (CoV-2-CNUHV03) using DNASTAR Lasergene.

**Fig. S10.**
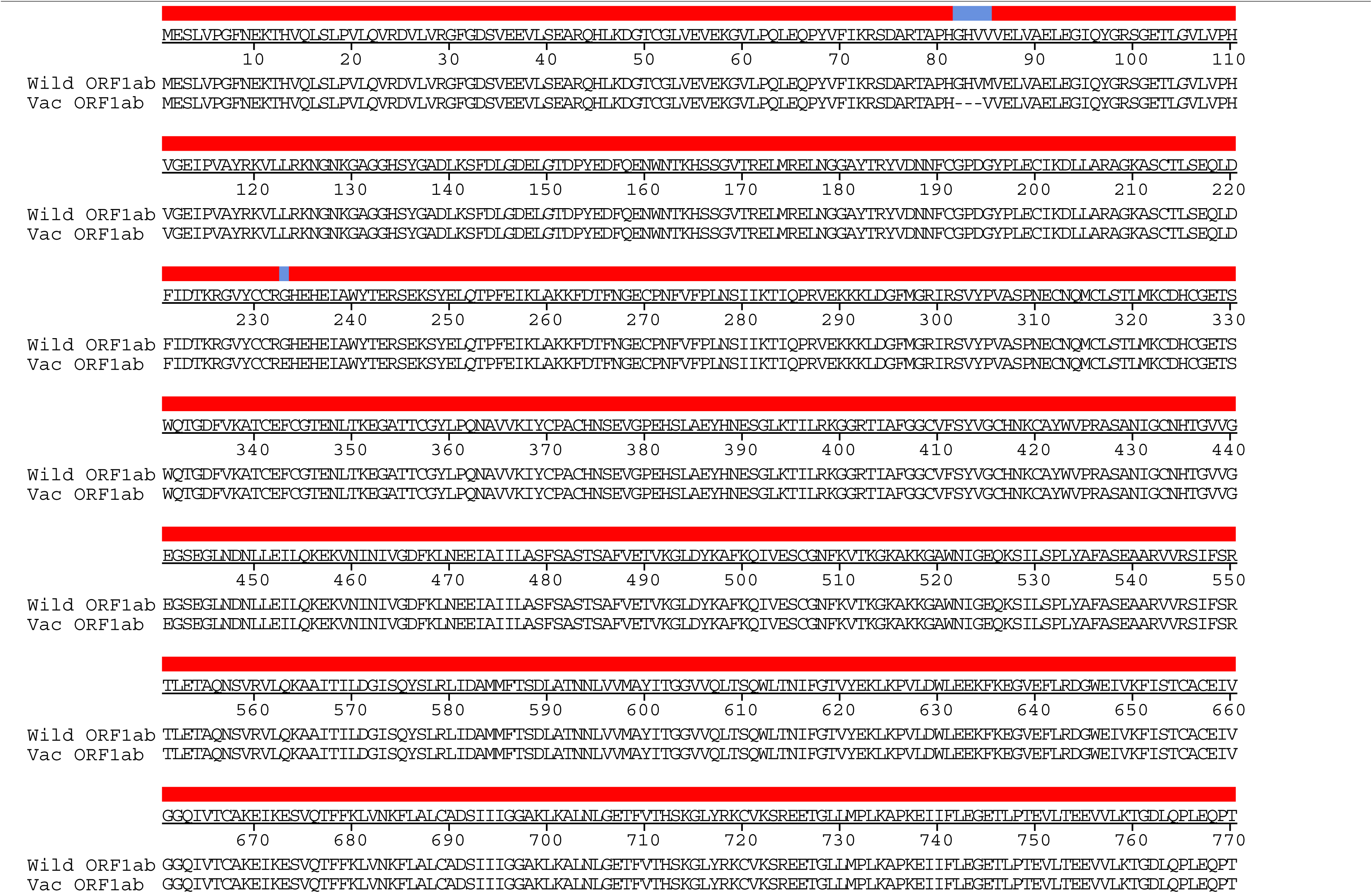
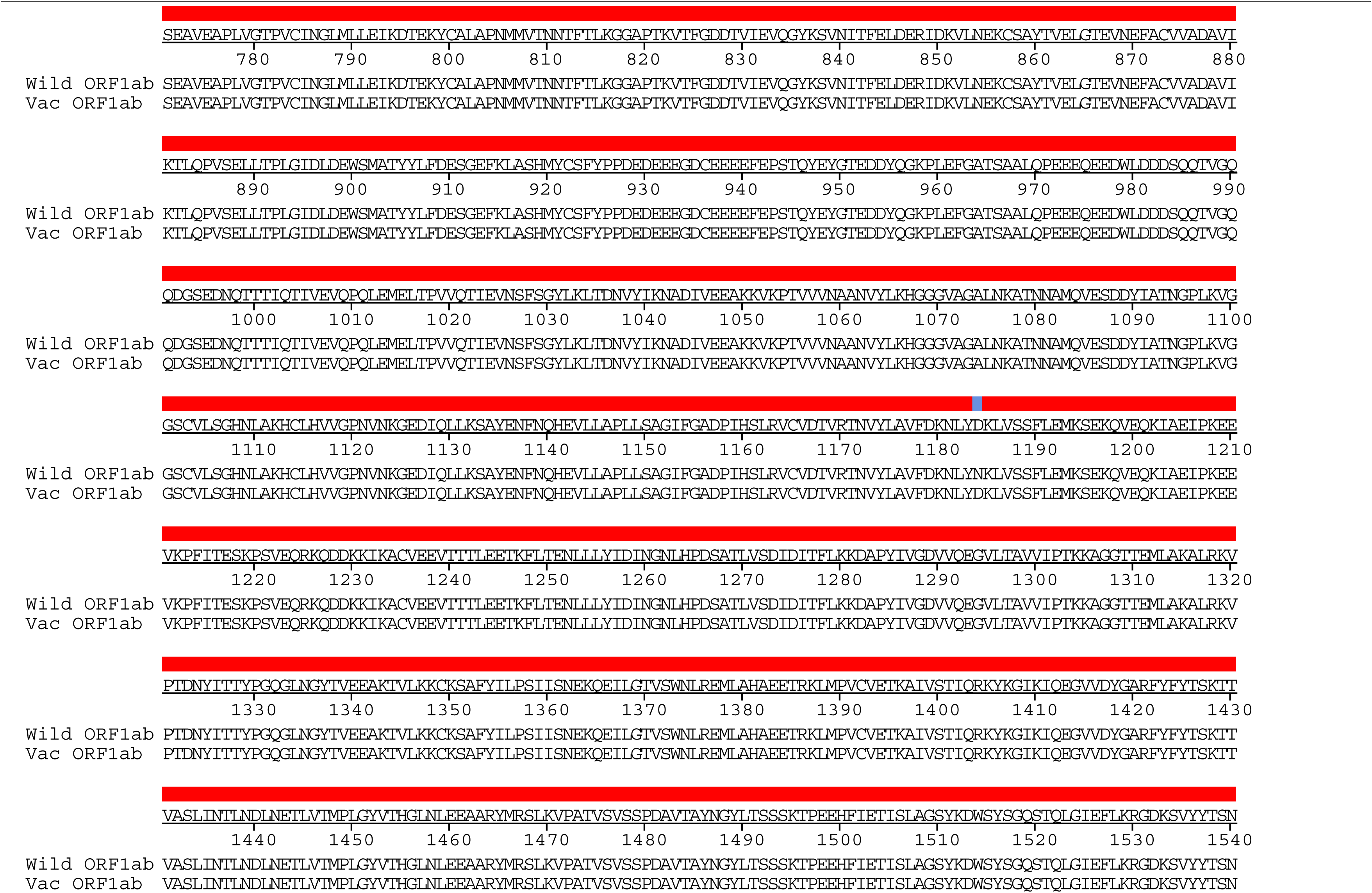
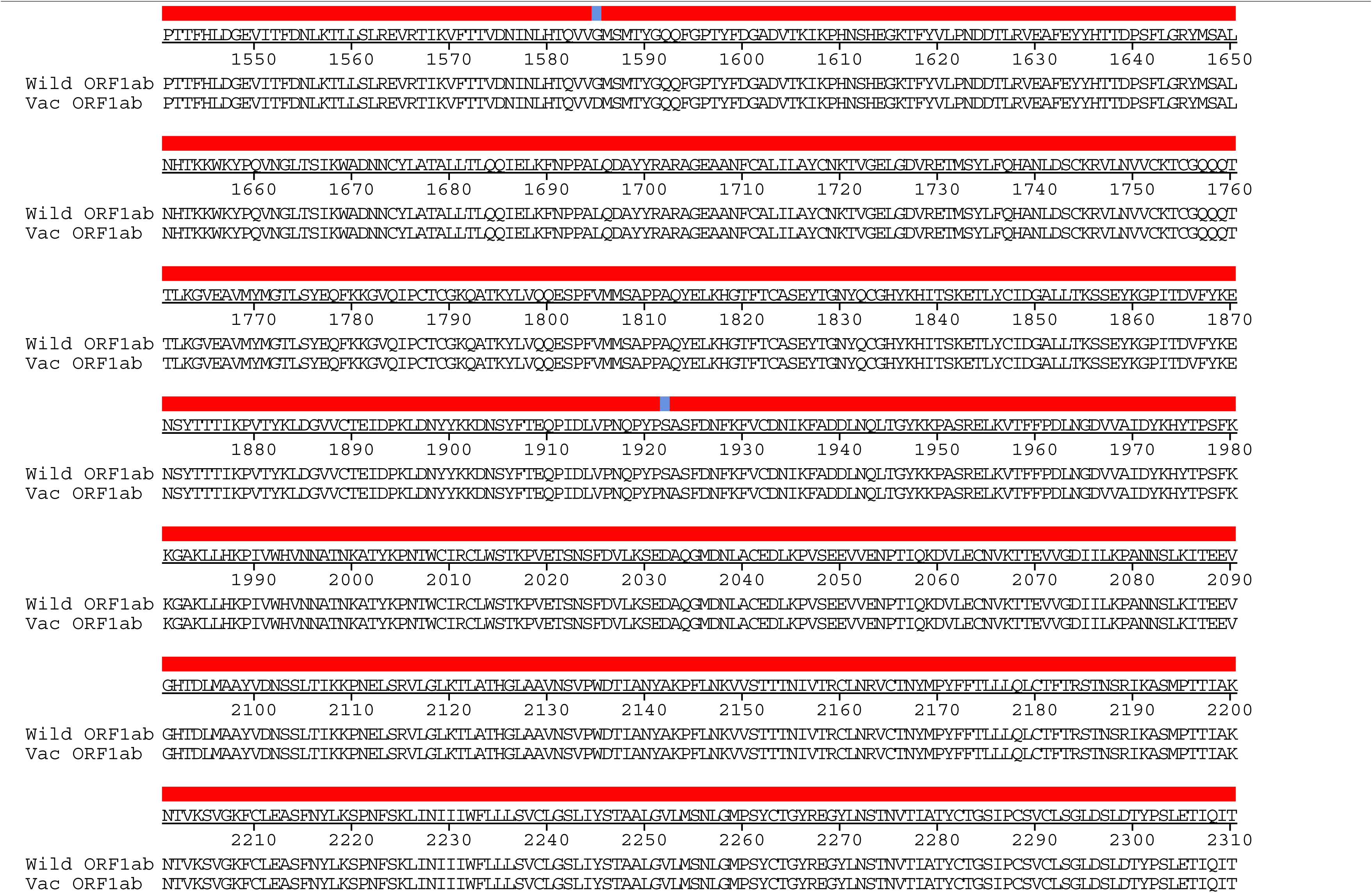
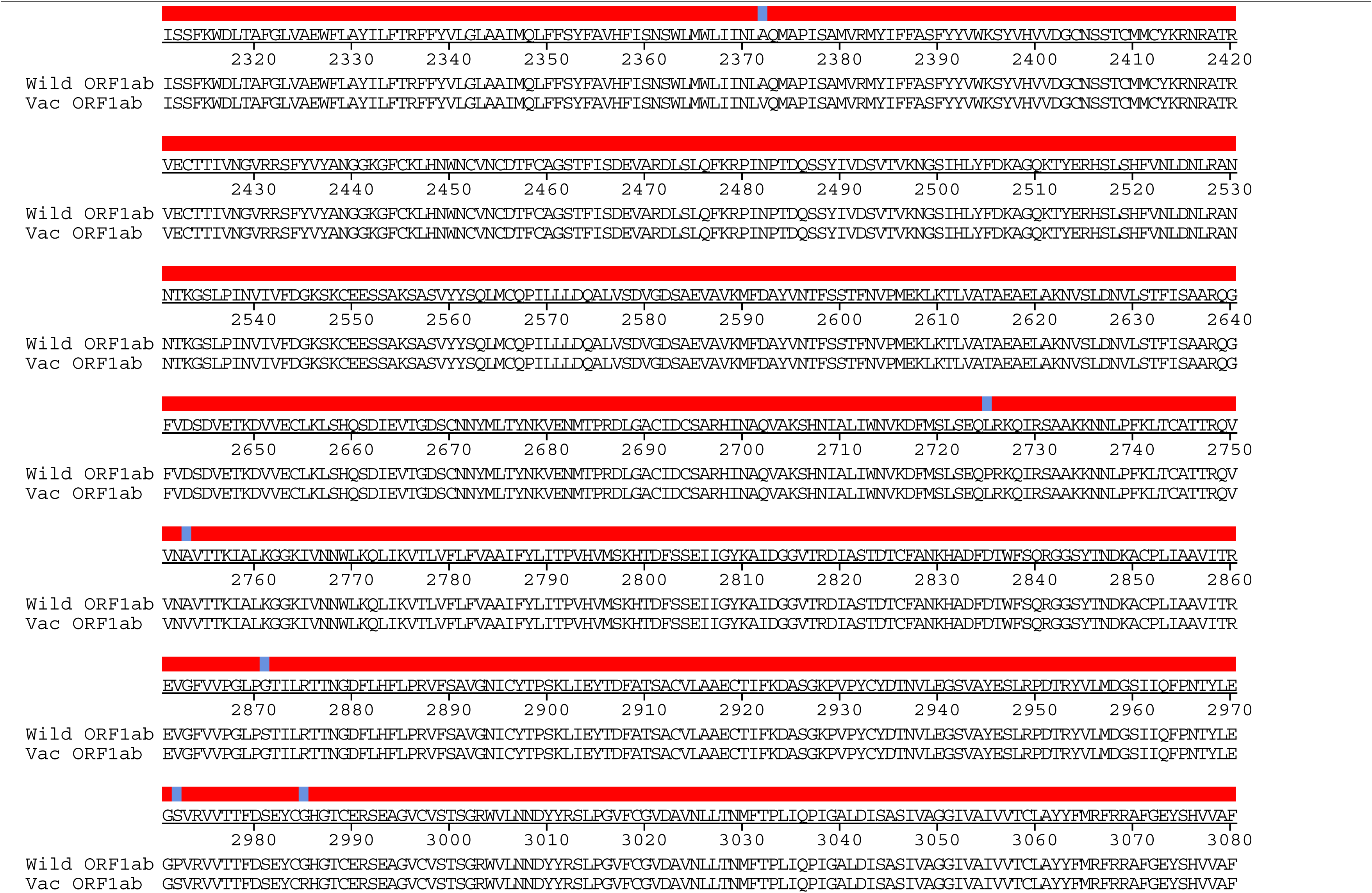
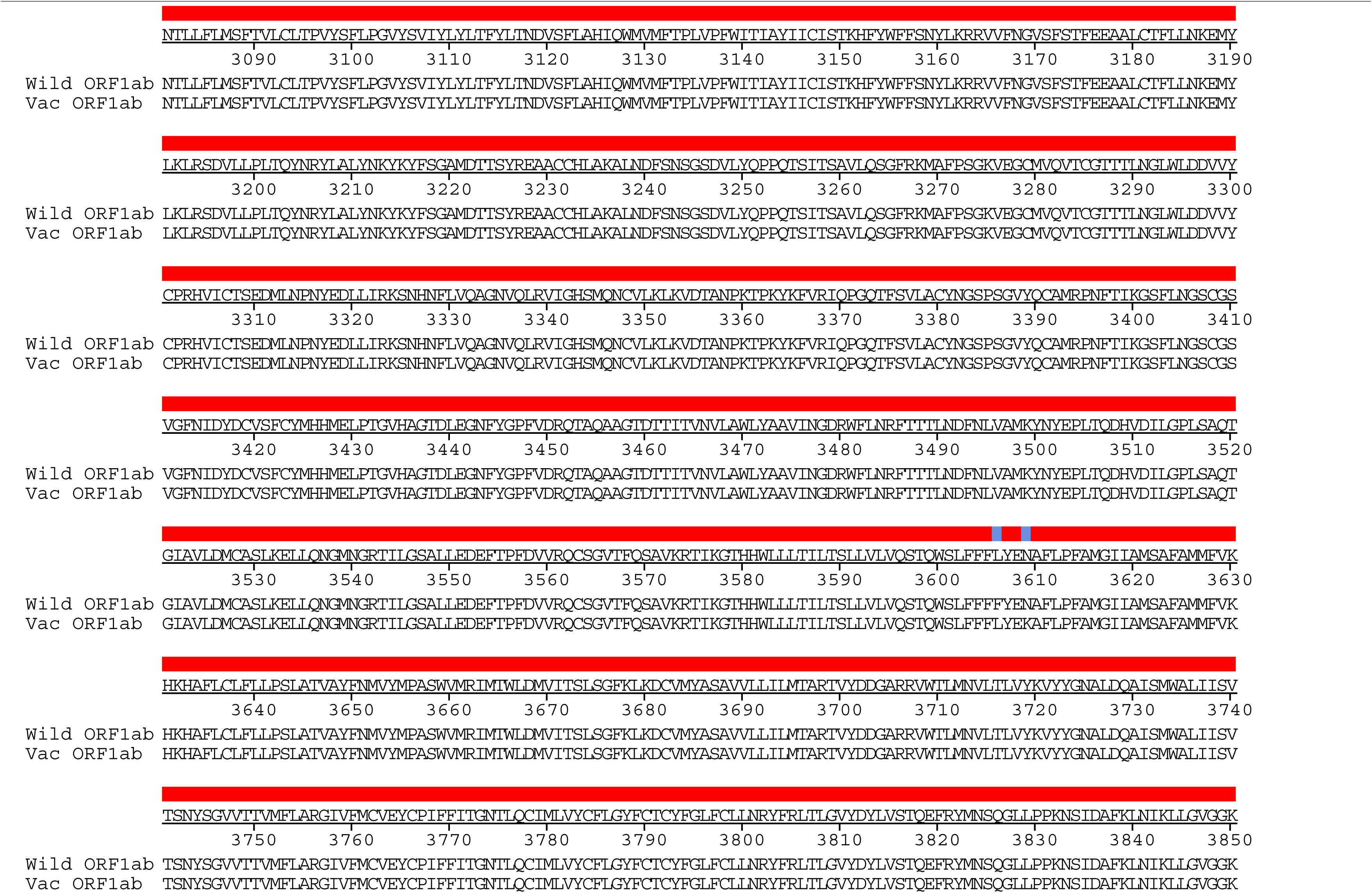
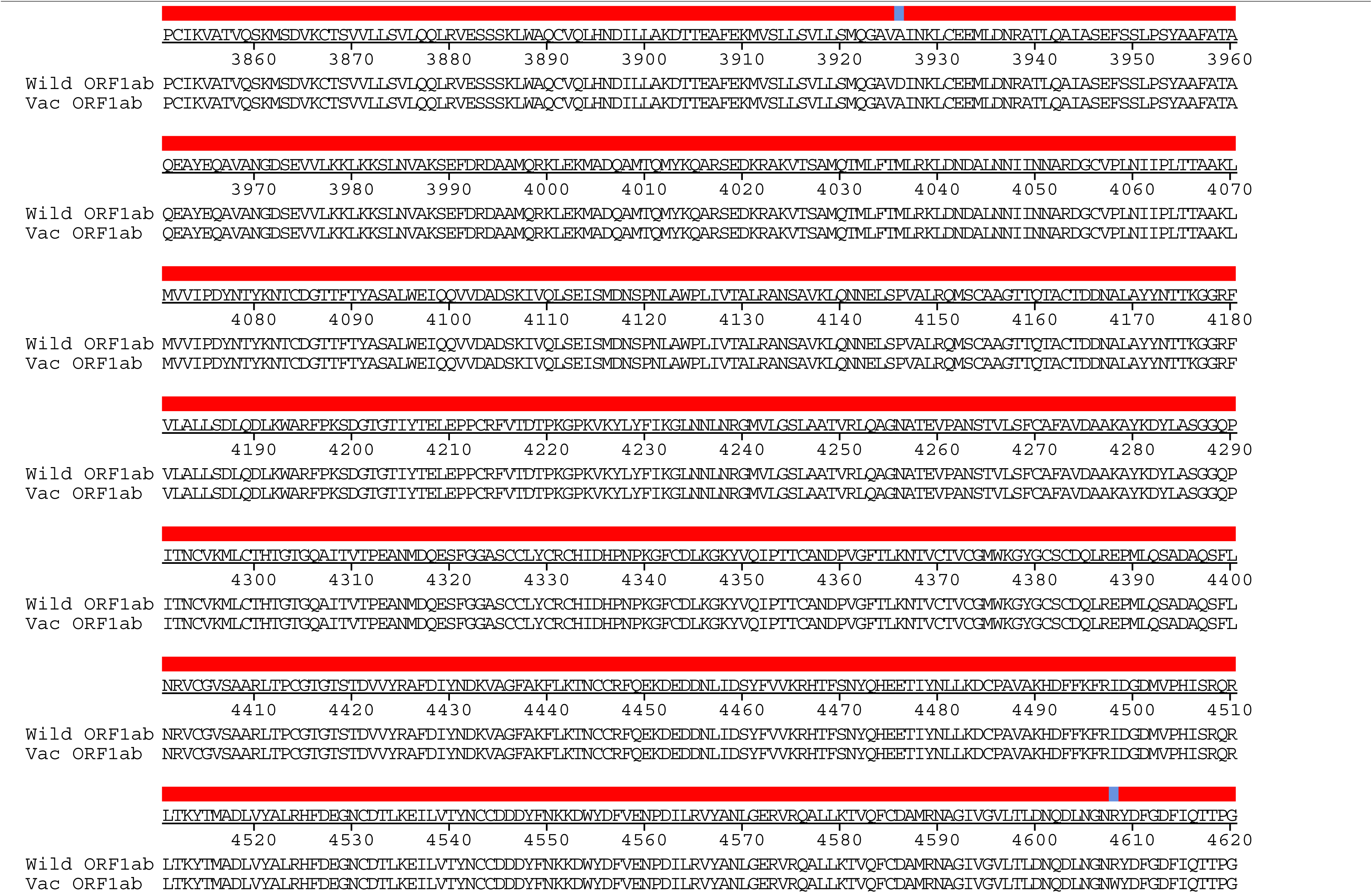
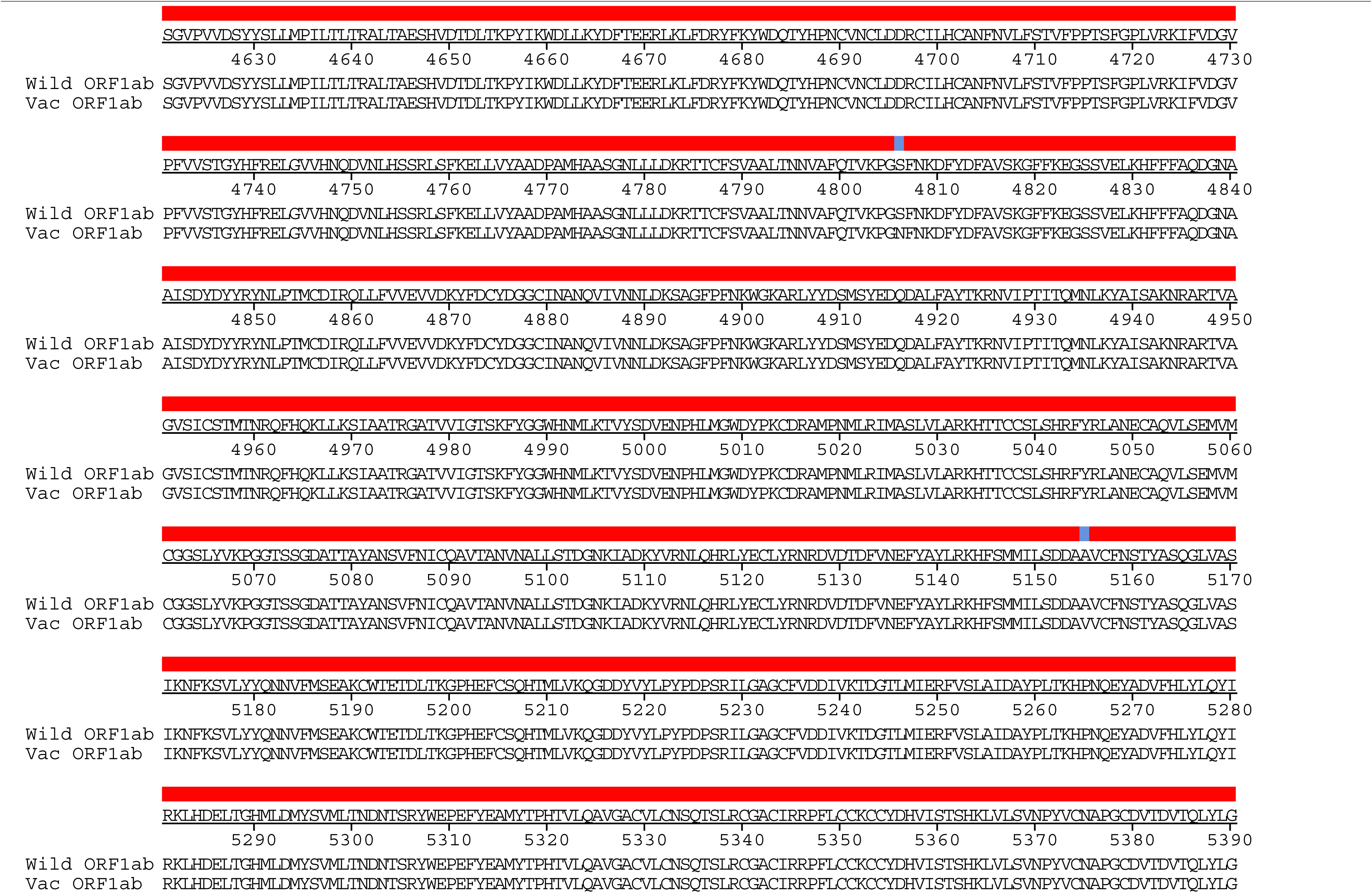
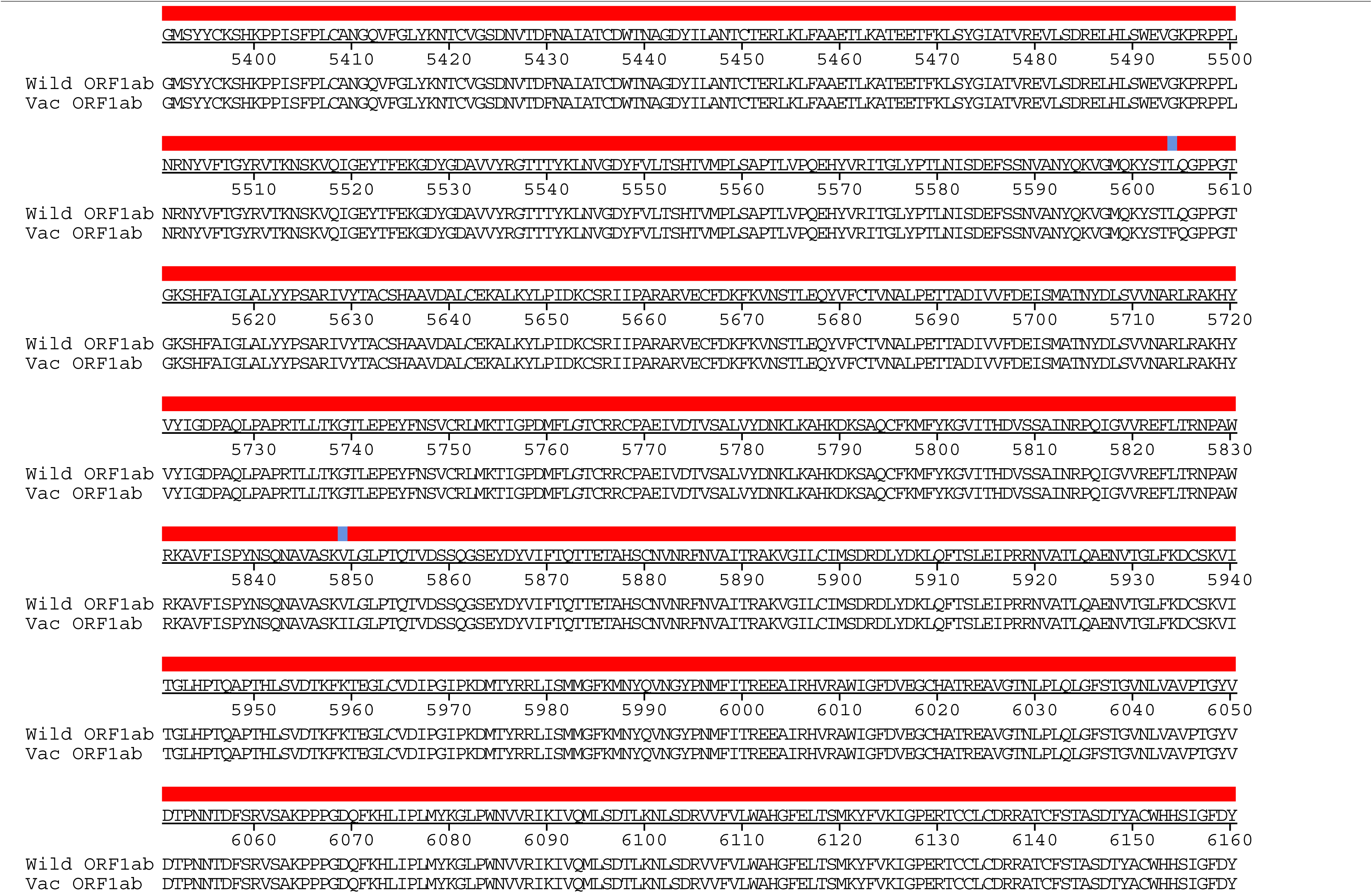
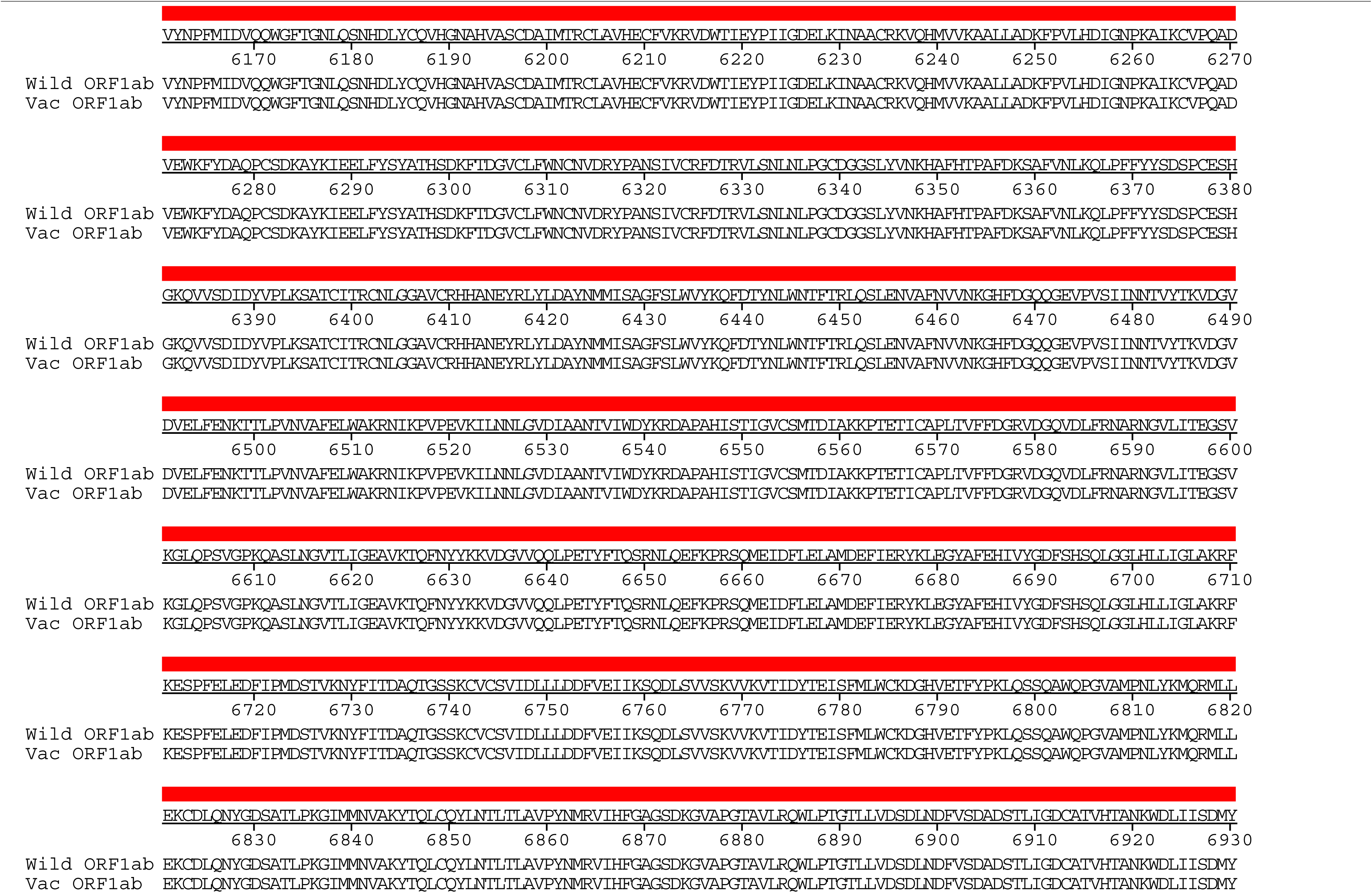
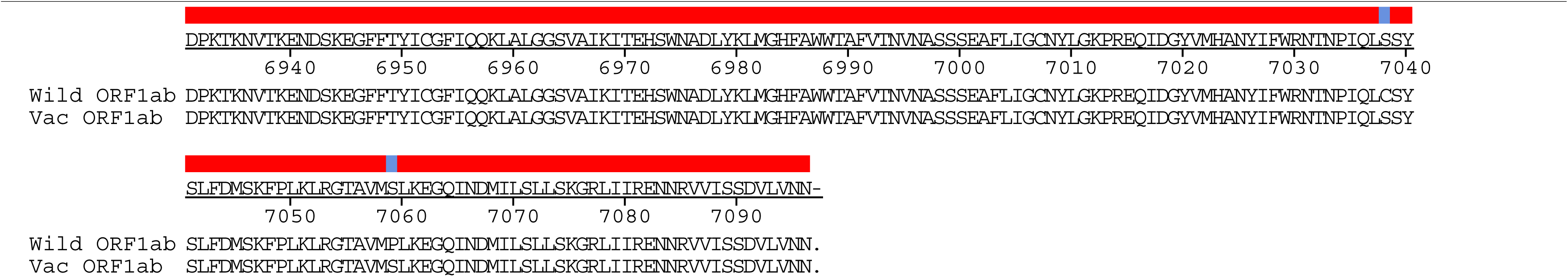
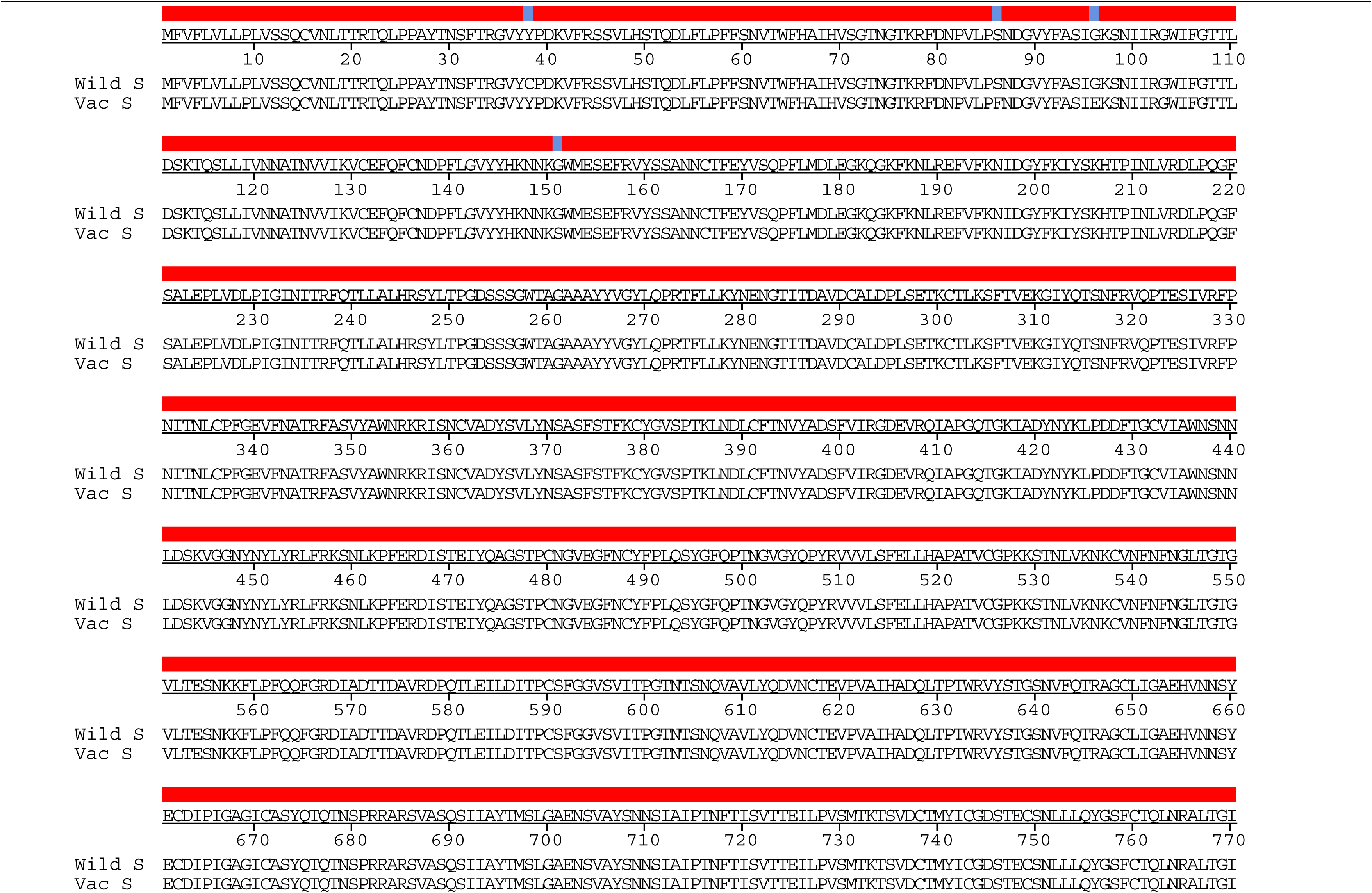
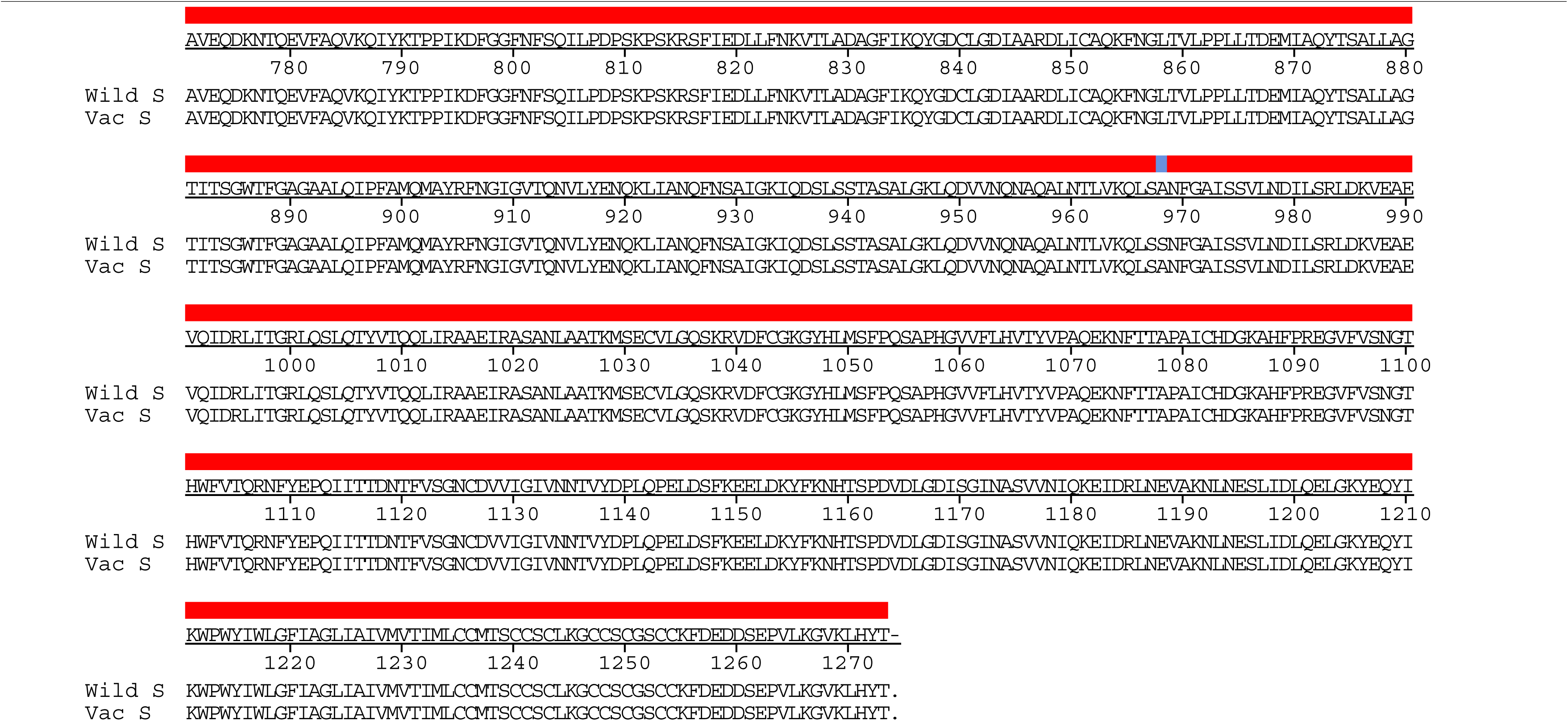
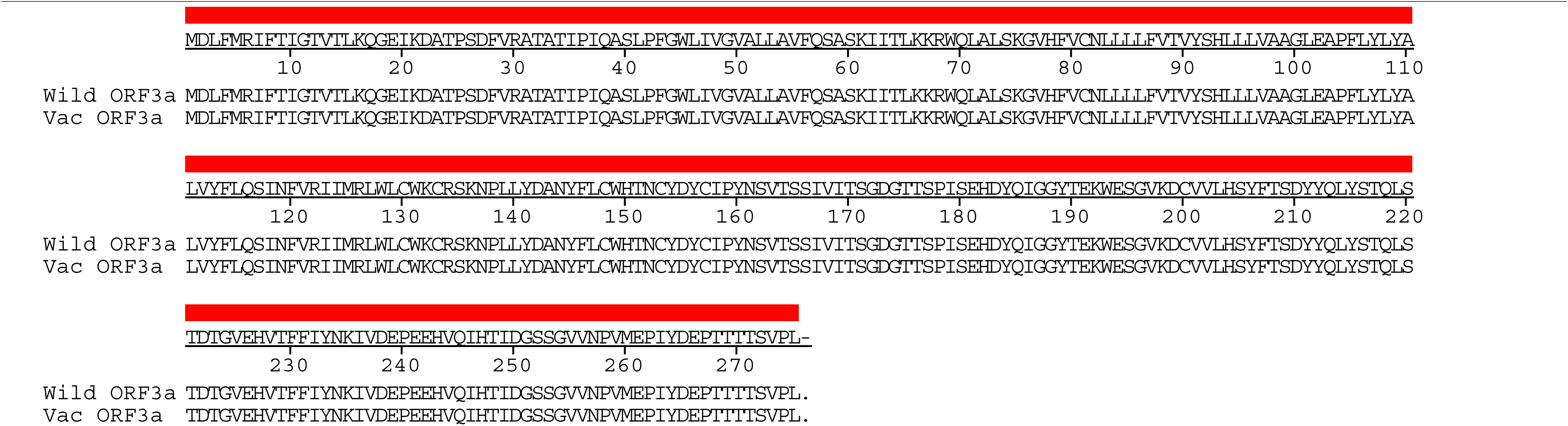
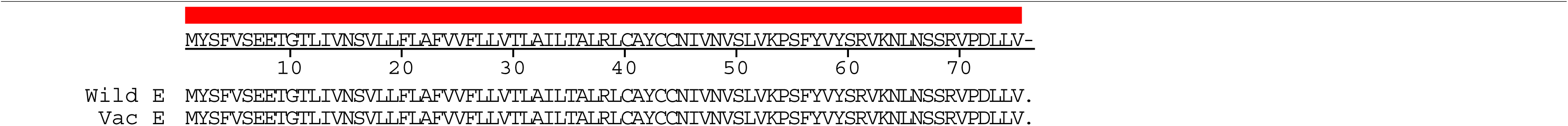
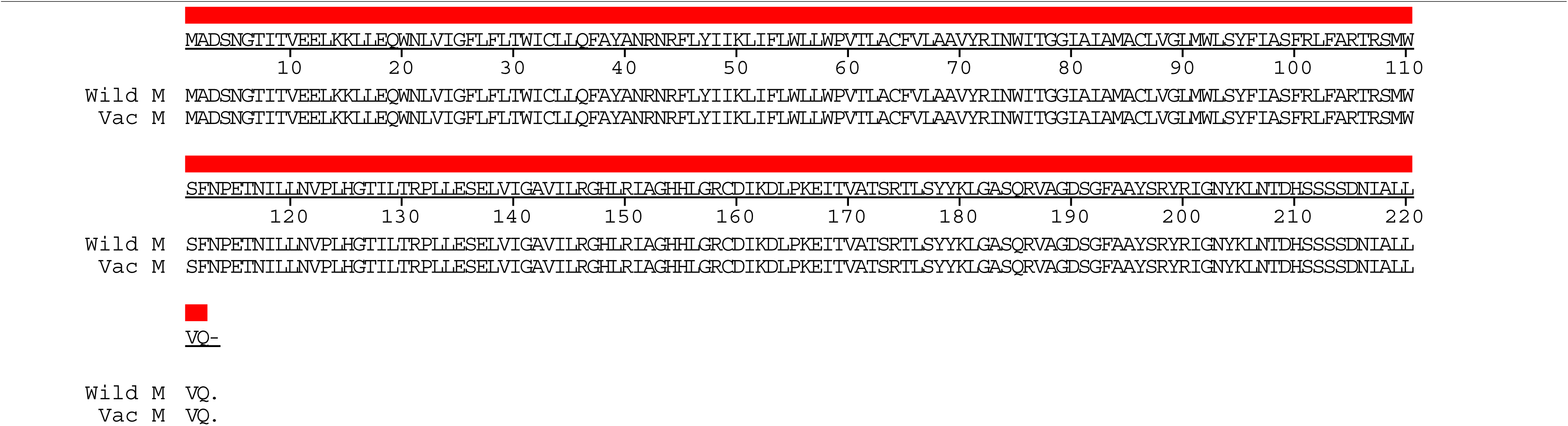
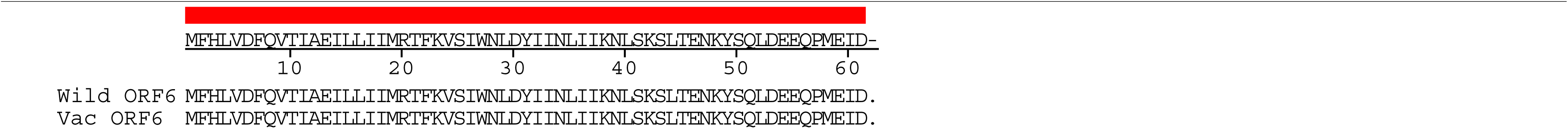
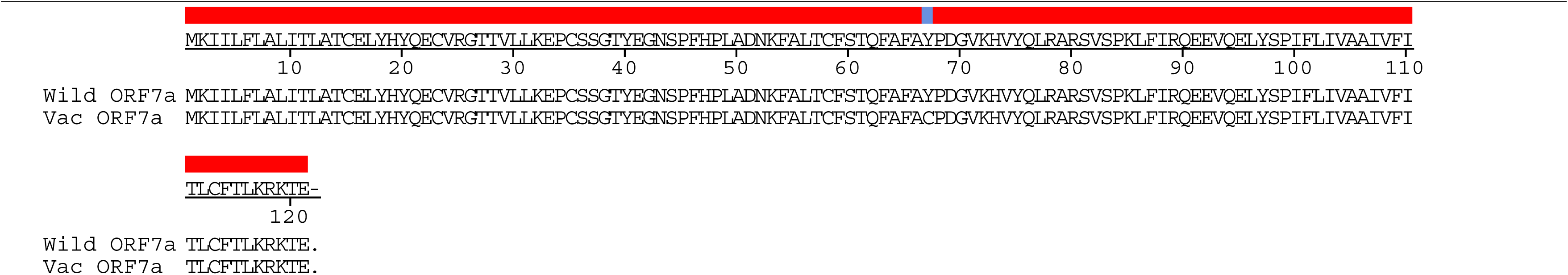
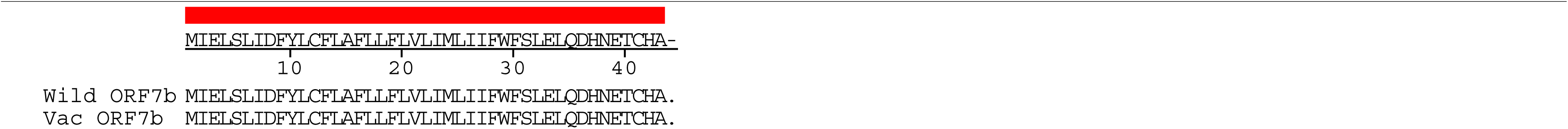
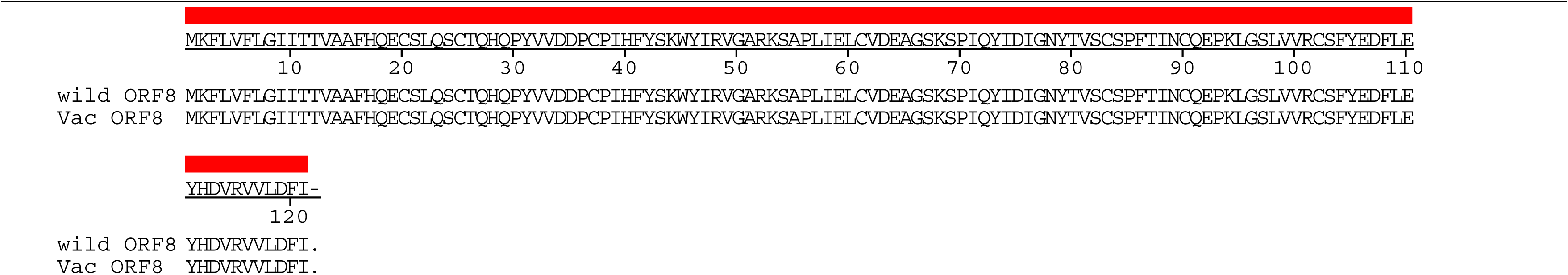
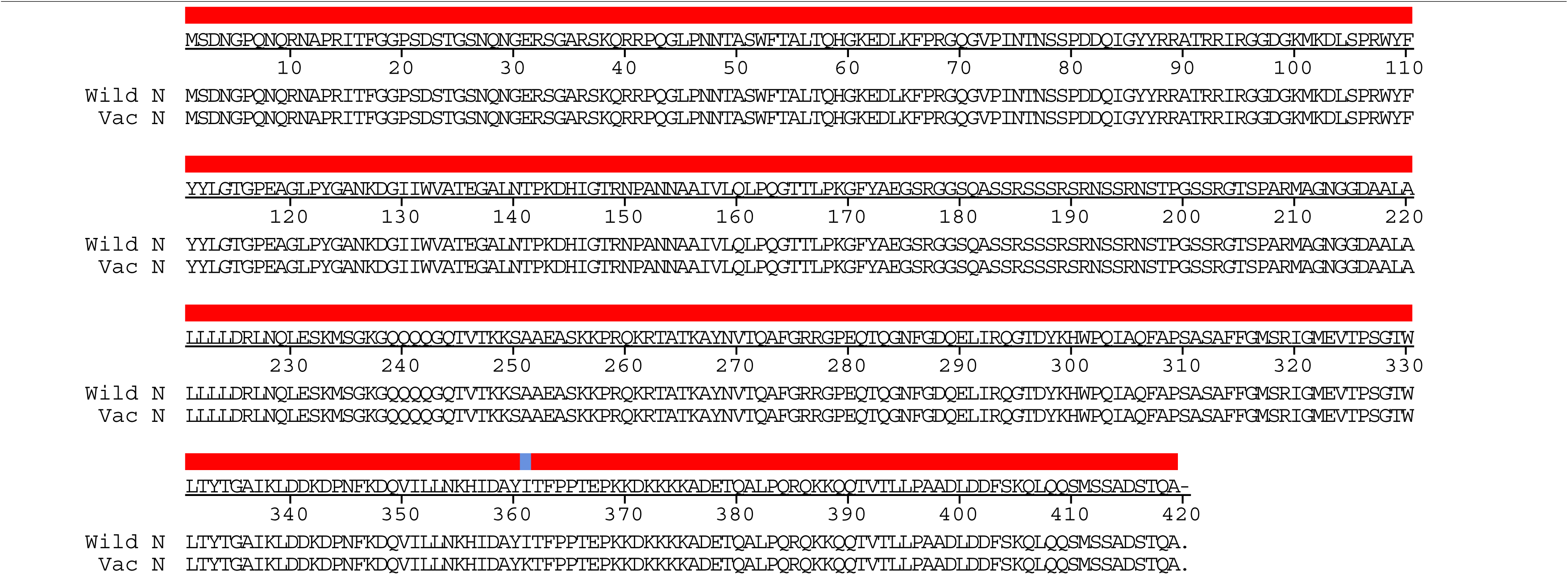
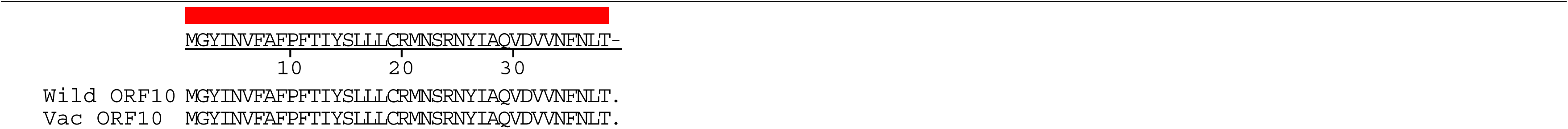
Amino acid alignment of cold-adapted live attenuated SARS-CoV-2 vaccine strain. Full amino acidsequences of cold-adapted live attenuated SARS-CoV-2 vaccine strain (CoV-2-CNUHV03-CA22℃) were aligned with that of wild-type SARS-CoV-2 (CoV-2-CNUHV03) using DNASTAR Lasergene.

